# A spatially aware likelihood test to detect sweeps from haplotype distributions

**DOI:** 10.1101/2021.05.12.443825

**Authors:** Michael DeGiorgio, Zachary A. Szpiech

## Abstract

The inference of positive selection in genomes is a problem of great interest in evolutionary genomics. By identifying putative regions of the genome that contain adaptive mutations, we are able to learn about the biology of organisms and their evolutionary history. Here we introduce a composite likelihood method that identifies recently completed or ongoing positive selection by searching for extreme distortions in the spatial distribution of the haplotype frequency spectrum along the genome relative to the genome-wide expectation taken as neutrality. Furthermore, the method simultaneously infers two parameters of the sweep: the number of sweeping haplotypes and the “width” of the sweep, which is related to the strength and timing of selection. We demonstrate that this method outperforms the leading haplotype-based selection statistics. As a positive control, we apply it to two well-studied human populations from the 1000 Genomes Project and examine haplotype frequency spectrum patterns at the *LCT* and MHC loci. We also apply it to a data set of brown rats sampled in NYC and identify genes related to olfactory perception. To facilitate use of this method, we have implemented it in user-friendly open source software.

## Introduction

The identification and classification of genomic regions undergoing positive selection in populations has been of long standing interest for studying organisms across the tree of life. By investigating regions containing putative adaptive variation, one can begin to shed light on a population’s evolutionary history and the biological changes well-suited to cope with various selection pressures.

The genomic footprint of positive selection is generally characterized by long high-frequency haplotypes and low nucleotide diversity in the vicinity of the adaptive locus, the result of linked genetic material “sweeping” to high frequency faster than mutation and recombination can introduce novel variation. These selective sweeps are often described by two paradigms—“hard sweeps” and “soft sweeps”. Whereas a hard sweep is the result of a beneficial mutation that brings a single haplotype to high frequency [Przeworski, 2002], soft sweeps are the result of selection on multiple haplotype backgrounds, often the result of selection on standing variation or a high adaptive mutation rate. Soft sweeps are thus characterized by multiple sweeping haplotypes rising to high frequency [Hermisson and Pennings, 2005, Pennings and Hermisson, 2006a].

Many statistics have been proposed to capture these patterns to make inferences about recent or ongoing positive selection [Sabeti et al., 2002, Voight et al., 2006, Sabeti et al., 2007, Ferrer-Admetlla et al., 2014, Garud et al., 2015, Field et al., 2016, Harris et al., 2018, Torres et al., 2018, Stern et al., 2019, Harris and DeGiorgio, 2020, Szpiech et al., 2021, Szpiech, 2021, Kim and Stephan, 2002, Nielsen et al., 2005, Chen et al., 2010, Huber et al., 2015, Vy and Kim, 2015, DeGiorgio et al., 2016, Racimo, 2016, Lee and Coop, 2017, Setter et al., 2020], many of which focus on summarizing patterns of haplotype homozygosity in a local genomic region. A particularly novel approach, the *T* statistic implemented in `LASSI` [Harris and DeGiorgio, 2020], employs a likelihood model based on distortions of the haplotype frequency spectrum (HFS). In this framework, Harris and DeGiorgio [2020] model a shift in the HFS toward one or several high-frequency haplotypes as the result of a hard or soft sweep in a local region of the genome. In addition to the likelihood test statistic *T*, for which larger values suggest more support for a sweep, `LASSI` also infers the parameter *m*. This parameter estimates the number of sweeping haplotypes in a genomic region, and *m >* 1 indicates support for a soft sweep.

A drawback of the original formulation of the *T* statistic implemented in `LASSI` is that it does not account for or make use of the genomic spatial distribution of haplotypic variation expected from a sweep. Specifically, Harris and DeGiorgio [2020] demonstrated that if the spatial distribution of *T* was directly accounted for in the machine learning approach (*Trendsetter*) of Mughal and DeGiorgio [2019], the power for detecting sweeps was greatly enhanced. Indeed, modern statistical learning machinery to detect sweeps has been greatly enhanced by incorporating spatial distributions of summary statistics [Lin et al., 2011, Schrider and Kern, 2016, Sheehan and Song, 2016, Kern and Schrider, 2018, Mughal and DeGiorgio, 2019, Mughal et al., 2020]. However, these machine learning methods need extensive simulations under an accurate and explicit demographic model to train the classifier. An alternative approach is to directly integrate this spatial distribution into the likelihood model, as has been performed for site frequency spectrum (SFS) composite likelihood methods to detect sweeps [Kim and Stephan, 2002, Nielsen et al., 2005, Chen et al., 2010, Huber et al., 2015, Vy and Kim, 2015, DeGiorgio et al., 2016, Racimo, 2016, Lee and Coop, 2017, Setter et al., 2020]. Here we incorporate the spatial distribution along the genome of HFS variation into the `LASSI` framework and introduce the Spatially Aware Likelihood Test for Improving `LASSI`, or `saltiLASSI`. For easy application to genomic datasets, we implement `saltiLASSI` in the open source program `lassip` along with `LASSI` [Harris and DeGiorgio, 2020], and other HFS-based statistics H12, H2/H1, G123, and G2/G1 [Garud et al., 2015, Harris et al., 2018]. `lassip` is available at https://www.github.com/szpiech/lassip.

After validating `saltiLASSI` through simulations and comparing it to other popular haplotype-based selection scans, we apply `saltiLASSI` to whole genome data from two different species. These data include two well-studied human populations (CEU and YRI) from the 1000 Genomes Project [The 1000 Genomes Project Consortium, 2015] and a population of brown rats sampled across the island of Manhattan in New York City (NYC), USA [Harpak et al., 2021]. Our analysis of the two human populations serves as a positive control in an empirical dataset with a well-studied demographic history. We reproduce several well-known signals of selection in the European CEU population and the African YRI population, including the *LCT* (CEU), MHC (CEU and YRI), and *APOL1* (YRI) loci, demonstrating that this method works well in real data. Our analysis of the NYC brown rat data serves as an example of applying the `saltiLASSI` method to a dataset with haplotype phase unknown and a poorly calibrated demographic history making neutral simulations contraindicated (see Harpak et al. [2021] on this point). Here, we find strong selection signals among clusters of genes related to olfactory perception.

## Results

In this section we begin by developing a new likelihood ratio test statistic, termed Λ, that evaluates spatial patterns in the distortion of the HFS as evidence for sweeps. We then demonstrate that Λ has substantially higher power than competing single-population haplotype-based approaches, across a number of model parameters related to the underlying demographic and adaptive processes. Similar to the *T* statistic implemented in the `LASSI` framework of Harris and DeGiorgio [2020], we also show that Λ is capable of approximating the softness of a sweep by estimating the current number of high-frequency haplotypes 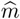. We then apply the Λ statistic to whole-genome sequencing data from two human populations from the 1000 Genomes Project [The 1000 Genomes Project Consortium, 2015] and a population of brown rats from NYC [Harpak et al., 2021].

### Definition of the statistic

Here we extend the `LASSI` maximum likelihood framework for detecting sweeps based on haplotype data [Harris and DeGiorgio, 2020], by incorporating the spatial pattern of haplotype frequency distortion in a statistical model of a sweep. Recall that Harris and DeGiorgio [2020] defined a genome-wide background *K*-haplotype truncated frequency spectrum vector

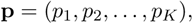

which they assume represents the neutral distribution of the *K* most-frequent haplotypes, with *p*_1_ ≥ *p*_2_ *≥· · ≥ p_K_ ≥* 0 and normalization such that 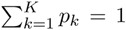. Harris and DeGiorgio [2020] then define the vector

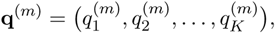

with 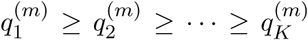 and 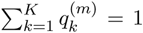. This represents a distorted *K*-haplotype truncated frequency spectrum vector in a particular genomic region with a distortion consistent with *m* sweeping haplotypes. To create the these distorted haplotype spectra, Harris and DeGiorgio [2020] used the equation

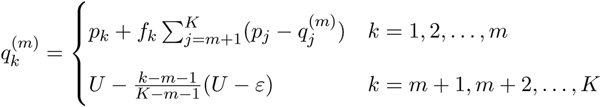

where *f_k_ ≥* 0 for *k ∈ {*1, 2, …, *m}* and 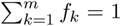, defines the way by which mass is distributed to the *m* “sweeping” haplotypes from the *K − m* non-sweeping haplotypes with frequencies *p_m_*_+1_, *p_m_*_+2_, …, *p_K_*. The variables *U* and *ε* are associated with the amount of mass from non-sweeping haplotypes that are converted to the *m* sweeping haplotypes [see Harris and DeGiorgio, 2020]. We choose to set *U* = *p_K_*, and then vary *ε ≤ U* during optimization. Harris and DeGiorgio [2020] propose several reasonable choices of *f_k_*, and for all computations here we use 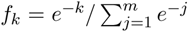. The schematic in Figure 1A illustrates the `LASSI` framework of generating the distorted haplotype spectra.

**Figure 1:**
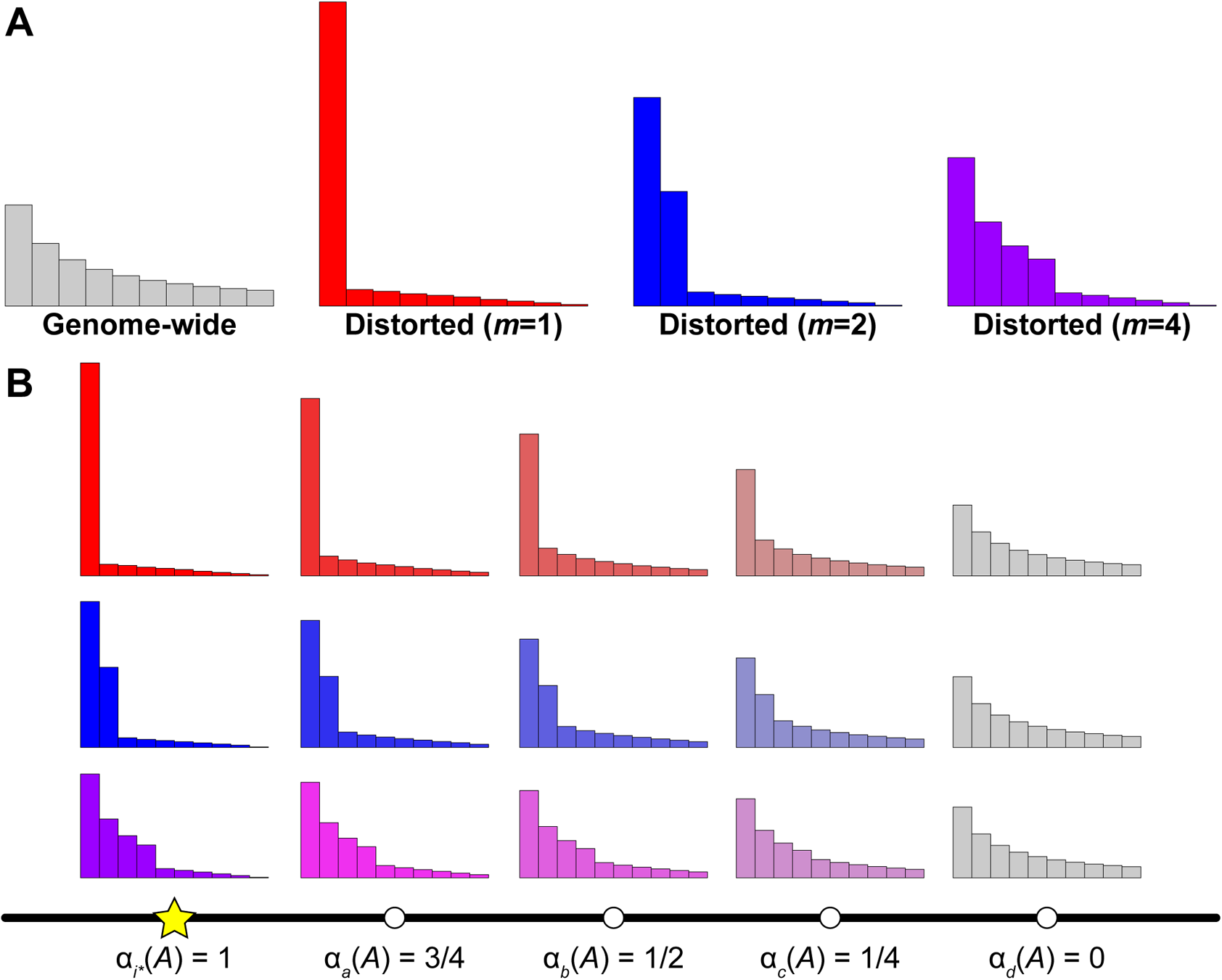
Schematic of the `saltiLASSI` mixture model framework. (A) Generation of distorted haplotype frequency spectra (HFS) for *m* = 1 (red), 2 (blue), and 4 (purple) sweeping haplotypes from a genome-wide (gray) neutral HFS under the `LASSI` framework of Harris and DeGiorgio [2020]. (B) Generation of spatially-distorted HFS under the `saltiLASSI` framework for a window *i* (white circles) with increasing distance from the sweep location (yellow star). When the window is on top of the sweep location, the HFS is identical to the distorted `LASSI` HFS, and *α_i_*(*A*) = 1. When a window is far from the sweep location, the HFS is identical to the genome-wide (neutral) HFS, and *α_i_*\* (*A*) = 0. For windows at intermediate distances from the sweep location, the HFS is a mixture of the distorted and genome-wide HFS, with the distorted HFS contributing *α_i_*(*A*) and the genome-wide HFS contributing 1 *α_i_*(*A*). We show example spectra at windows *a*, *b*, *c*, and *d* that are of increasing distances from the sweep location *i^*^*, with *i^*^ < a < b < c < d*.

To incorporate the spatial distribution haplotypic variation into the `LASSI` framework, consider an index set 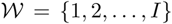 of 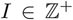 contiguous (potentially overlapping) windows such that window 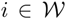 has position along a chromosome denoted *z_i_*. This position could be in physical units (such as bases), in genetic map units (such as centiMorgans), in number of polymorphic sites [such as employed by *nS_L_* in Ferrer-Admetlla et al., 2014], or in window number. We model the relative contribution of a sweep with *m* sweeping haplotypes at target window with index 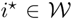 by a parameter *α_i_ ∈* [0, 1] on window 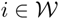 and the relative contribution of neutrality by 1 *− α_i_*.

Following a similar powerful framework introduced by Cheng and DeGiorgio [2020] for modeling balancing selection, we employ a mixture model to model the *K*-haplotype truncated frequency spectrum in window *i*, with a proportion

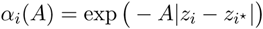

deriving from a sweep model and a proportion 1 *− α_i_*(*A*) deriving from the genome-wide background haplotype spectrum to represent neutrality. Here, *A* is a parameter that we optimize over, describing the rate of decay of the effect of the sweep at target window *i^*^* on the flanking windows a certain distance away. Specifically, we model the *K*-truncated haplotype spectrum in window *i* as the vector

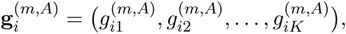

where

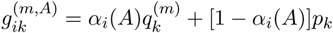

for *k* = 1, 2, …, *K* and 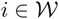. Note here that for target window *i*, *α_i*_* (*A*) = 1, and hence 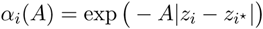 *i.e.*, the target window is on top of the sweep, and so it is entirely determined by the distorted *m*-sweeping haplotype spectrum. However given a fixed *A* value, for windows *i* far enough away from the central window *i^*^*, we have the *α_i_*(*A*) = 0, and therefore 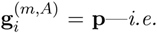, the expectation of a neutral window. Based on these trends, windows far from the putatively selected target window are modeled as neutral, and windows close to the target window are heavily distorted due to the sweep. Moreover, because *α_i_*(*A*) tends to zero for windows far enough away for the central window, the model of neutrality is nested within our proposed sweep model. The schematic in Figure 1B illustrates the `saltiLASSI` framework of generating the spatially-distorted haplotype spectra.

Assume that in window 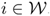, there is a *K*-truncated vector of counts

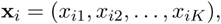

which are the observed counts of the *K* most-frequent haplotypes, with *x_i_*_1_ *≥ x_i_*_2_ *≥ · · · ≥ x_iK_ ≥* 0 and normalized such that 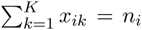, where *n_i_* is the total number of sampled haplotypes in window *i*. Following Cheng and DeGiorgio [2020] and Harris and DeGiorgio [2020], we then compute the log composite likelihood ratios for null hypothesis of neutrality at target window *i^*^* as

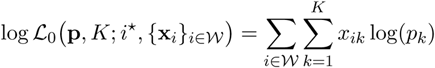

and for the alternative hypothesis of *m* sweeping haplotypes at target window *i^*^* as

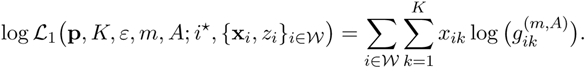

Using these log likelihoods, we follow Harris and DeGiorgio [2020] and construct a log likelihood ratio test statistic of a sweep at target window *i^*^* as

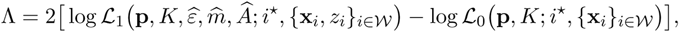

where

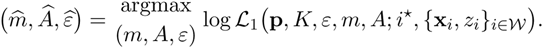

We note that this approach treats windows as independent in the null and alternative hypotheses, thus making it a composite likelihood method that ignores recombination.

### Computing the likelihood

To apply the `saltiLASSI` method, we compute Λ at each window in the genome, where each window is considered the target window *i^*^* in turn, and the likelihood is maximized independently for each target window. That is, all parameters (*m*, *A*, and *ε*) are optimized at each target window *i^*^*, thereby permitting the footprint size *A* of the sweep to vary across the genome, adjusting for initial linkage disequilibrium and local recombination rates that could impact sweep signals. Similar to the way SweepFinder [Nielsen et al., 2005], SweepFinder2 [DeGiorgio et al., 2016], and `LASSI` [Harris and DeGiorgio, 2020] approach maximization, we optimize the likelihood via a grid search across *m* ∈ {1, 2, …, *K}*, *ε ∈* [1*/*(100*K*), *U*], and *A ∈ {A*_min_, …, *A*_max_}. Here, *A*_min_ = *−* ln 0.99999*/d*_min_, representing a value of *A* with a slow decay with distance; *A*_max_ = *−* ln 0.00001*/d*_min_, representing a value of *A* with a fast decay with distance; and *d*_min_ is the smallest distance between any two windows genome-wide. We make 100 equally spaced (in log-space) steps between *A*_min_ and *A*_max_. Furthermore, in order to reduce computational burden, we pre-compute 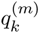 values across this grid for all windows.

### Power to detect sweeps

The power to detect sweeps will depend on a number of factors, including window size used to compute a statistic, whether phasing information for genotypes is used, the selection strength of the beneficial mutation *s*, the age of the sweep *t* (*i.e.*, time at which the selected mutation became beneficial), the number of selected haplotypes *ν*, and the underlying demographic history. To explore the power of Λ, we evaluate its power to detect sweeps of varying strengths, softness, and ages. For sweep settings, we considered only simulations in which the beneficial mutation established by reaching a frequency of at least 0.1, but we did not condition on fixation. Under each setting, we interrogated its robustness to demographic history, both through idealized constant-size histories and histories with recent severe bottlenecks. Moreover we gauged whether Λ yields false sweep signals under settings of background selection. Furthermore, for each setting described, we investigated the power and robustness of using unphased multilocus genotypes as input to Λ instead of phased haplotypes. In addition, we evaluated the effect of sample size *n*, number of haplotypes *K* to truncate the HFS, and recombination rate variation on the power of Λ to detect sweeps. Finally, we compared Λ to competing contemporary methods that use the same type of input data, using the *T* statistic of Harris and DeGiorgio [2020] for phased and unphased input data, and also considered the H12 [Garud et al., 2015], *nS_L_* [Ferrer-Admetlla et al., 2014], and iHS [Voight et al., 2006] statistics for phased data and the G123 statistic [Harris et al., 2018] for unphased data. The simulation protocol for all settings is described in the *Methods* section.

To begin, we compare the performance of Λ to *T*, H12, *nS_L_*, and iHS under a constant-size demographic history with diploid effective size of *N* = 10^4^ diploid individuals. The Λ, *T*, and H12 statistics were computed for different window sizes, consisting of 51, 101, or 201 SNPs per window. Figures 2A and S1 show that across sweeps of varying degrees of softness (beneficial mutation on ν ∈ {1, 2, 4, 8, 16} distinct haplotypes) and for sweeps of varying per-site per-generation strengths of s ∈ {0.01, 0.1}, the method with highest power regardless of time of selection (*t ∈ {*500, 100, 1500, 2000, 2500, 3000} generations prior to sampling) is Λ, thereby outperforming the competing methods. Interestingly, Λ applied to 51 SNP windows has generally higher power than with 101 and 201 SNP windows. Furthermore, smaller window sizes enable Λ to achieve high power even for old sweeps—with this elevated power often substantially higher than the closest competing method. This result recapitulates a finding of Harris and DeGiorgio [2020], where they observed that if the spatial distribution of the *T* statistic was used within a machine learning framework, computing the *T* statistic in a greater number of small windows yielded higher power for ancient sweeps than when a smaller number of large windows was used. This is an intriguing result, because smaller windows have poorer estimates of the distortion of the HFS, yet it appears that for detecting ancient sweeps what matters is capturing the overall spatial trend of the distortion of the HFS. That is, when using too large of windows, Λ is averaging the HFS across too large of a region, which has likely been broken up over time due to recombination for ancient sweeps. Instead, smaller windows focus on genomic segments with less shuffling of haplotype variation due to recombination events, such that distortions in the HFS are due to the effect of a sweep at a nearby selected site.

**Figure 2:**
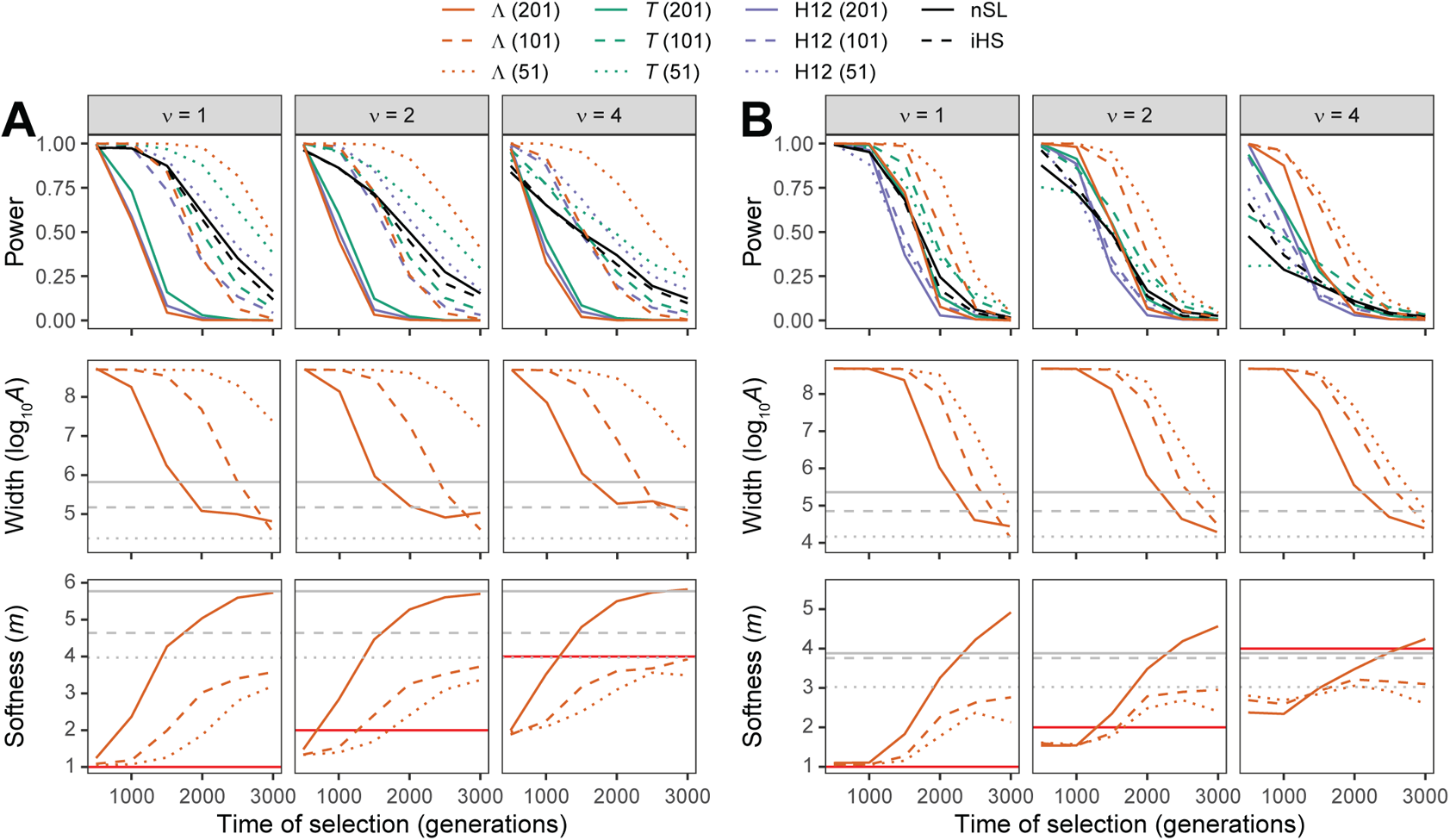
Performance of detecting and characterizing sweeps for applications of Λ, *T*, and H12 with windows of size 51, 101, and 201 SNPs, as well *nS_L_* and iHS under simulations of (A) a constant-size demographic history or (B) the human central European (CEU) demographic history of Terhorst et al. [2017]. Results are based on a sample of *n* = 50 diploid individuals and the haplotype frequency spectra for the Λ and *T* statistics truncated at *K* = 10 haplotypes. (Top row) Power at a 1% false positive rate as a function of selection start time. (Middle row) Estimated sweep width illustrated by mean estimated genomic size influenced by the sweep (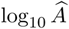) as a function of selection start time. Gray solid, dashed, and dotted horizontal lines are the corresponding mean 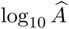 values for Λ applied to neutral simulations. (Bottom row) Estimated sweep softness illustrated by mean estimated number of sweeping haplotypes (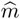) as a function of selection start time. Gray solid, dashed, and dotted horizontal lines are the corresponding mean 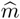 values for Λ applied to neutral simulations, and the red solid horizontal lines correspond to the number of sweeping haplotypes *ν* ∈ {1, 2, 4} assumed in sweep simulations. Sweep scenarios consist of hard (*ν* = 1) and soft (*ν* ∈ {2, 4}) sweeps with per-generation selection coefficient of *s* = 0.1 that started at *t* ∈ 500, 1000, 1500, 2000, 2500, 3000 generations prior to sampling. Results expanded across wider range of simulation settings can be found in Figures S1–S3 and S7–S9 as well as results for application to unphased multilocus genotype data in Figures S4–S6 and S10–S12.

Figure S1 also highlights a key distinction among sweeps of different strengths. Specifically, regardless of method considered, each achieves its highest power when sweeps of strength *s* = 0.1 are recent, whereas for sweeps of strength *s* = 0.01, highest power for each method is shifted farther in the past toward more ancient sweep. This pattern was also found previously for H12 [Harris et al., 2018] and *T* [Harris and DeGiorgio, 2020]. The likely reason for this result is that sweeps of strength *s* = 0.01 require more time for the beneficial allele to reach high frequency and leave a conspicuous genomic footprint, with this greater time to reach high frequency associated with increased chance that recombination and mutation act to break up high-frequency haplotypes. In contrast, sweeps of strength *s* = 0.1 create an immediate selection signature to appear in the genome due to the rapid rise in frequency of a beneficial mutation, but traces of this sweep pattern erode over time due to recombination, mutation, and drift. However, regardless, the Λ statistic paired with a small window size yields uniformly better or comparable sweep detection ability than the other approaches we examined. We also found that all methods performed poorly when selection strength was *s* = 0.001.

During a scan with Λ, the composite likelihood ratio is optimized over the number of high frequency (sweeping) haplotypes *m* and the footprint size of the sweep *A*, leading to respective estimates 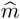 and 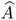. Therefore, at a genomic location with evidence for a sweep (high Λ value), we may better understand properties of the putative sweep by evaluating its softness through 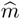 and its strength or age through 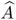. Figure S2 shows that for sweeps of strength *s* = 0.01, the estimated number of sweeping haplotypes 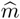 is considerably different from the actual number of initially-selected haplotypes *ν*, regardless of window size used or age of the sweep. In contrast, Figures 2A and S2 reveal that for hard sweeps (*ν* = 1) of strength *s* = 0.1, the estimate of the number of sweeping haplotypes when using 51 SNP windows is often consistent with hard sweeps (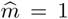) provided that the sweep is recent enough (within the last 500 generations). Similarly, under these same settings but with soft sweeps of *ν ∈ {*2, 4, 8, 16} selected haplotypes (Figures 2A and S2), the estimated number of sweeping haplotypes tends to be underestimated (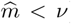) but is still consistent with a soft sweep (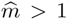). Therefore, provided that a sweep is recent enough, when using 51 SNP windows the value of the estimated number of sweeping haplotypes can be used to lend evidence of a hard (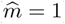) or a soft (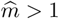) sweep.

Similarly, the other parameter estimate 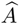 may also help characterize identified sweeps. Specifically, Figures 2A and S3 show that the footprint size of the sweep (measured as 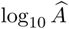) is substantially elevated compared to expectation for neutral simulations for sweep times at which there is high power to detect sweeps (Figures 2A and S1). Interestingly, the shape of the curves relating the mean sweep footprint size over time mirror the power of the Λ statistic with corresponding window size as a function of sweep initiation time (*t*), sweep softness (*ν*), and sweep strength (*s*). These results suggest that the estimate of the sweep footprint size (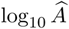) can be used to learn about the age or strength of a candidate sweep (the signatures of which appear to be confounded between the two parameters). Coupled with an estimate of the sweep softness (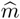), our `saltiLASSI` framework provides a means to not only detect sweeps with high power, but to also learn the underlying parameters that may have shaped the adaptive evolution of candidate sweep regions.

**Figure 3:**
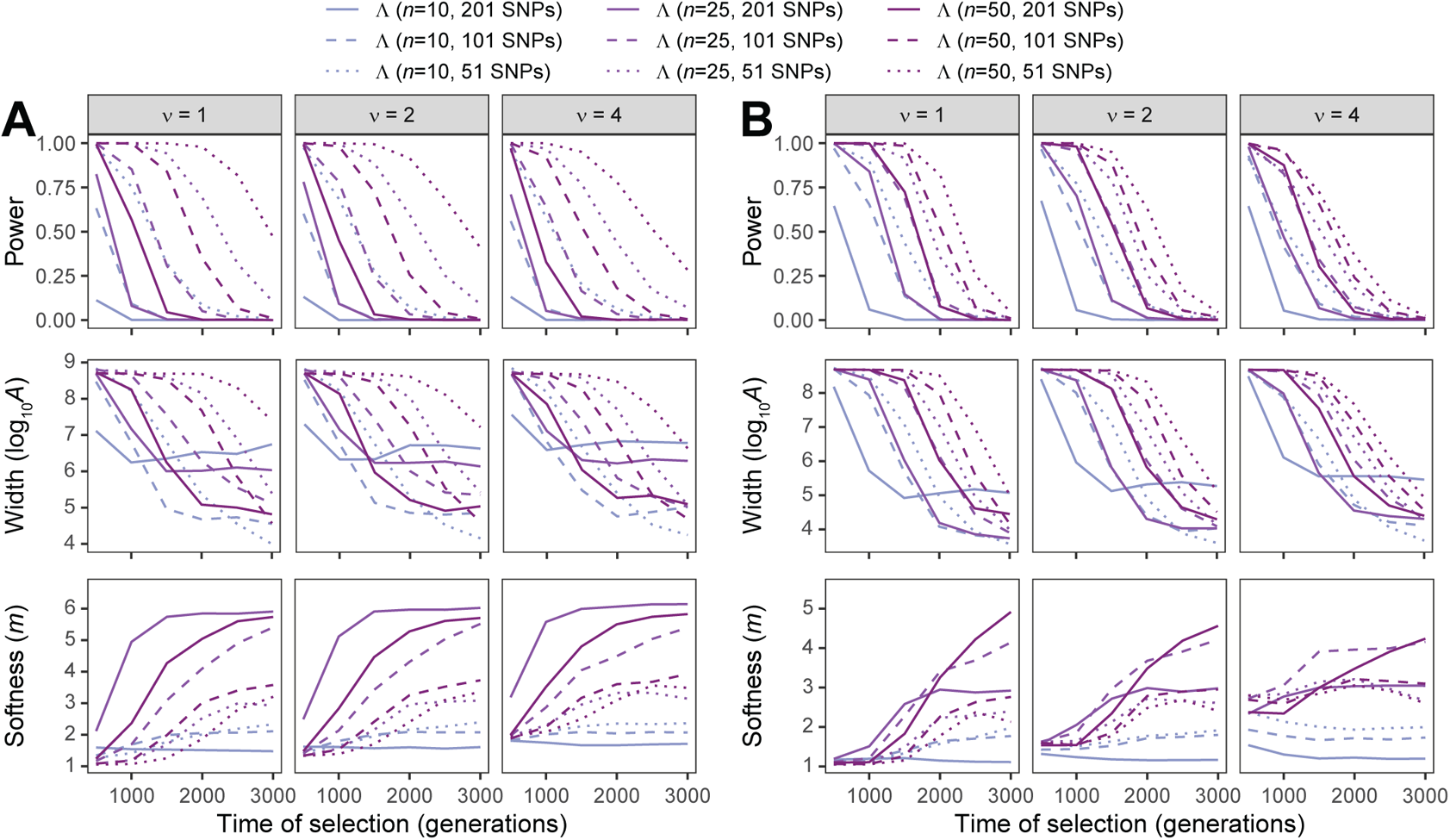
Performance of detecting and characterizing sweeps for applications of Λ with windows of size 51, 101, and 201 SNPs under simulations of (A) a constant-size demographic history or (B) the human central European (CEU) demographic history of Terhorst et al. [2017] and sample size of *n ∈* {10, 25, 50} diploid individuals. Results are based on the haplotype frequency spectra for the Λ statistic truncated at *K* = 10 haplotypes. (Top row) Power at a 1% false positive rate as a function of selection start time. (Middle row) Estimated sweep width illustrated by mean estimated genomic size influenced by the sweep (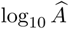) as a function of selection start time. (Bottom row) Estimate sweep softness illustrated by mean estimated number of sweeping haplotypes (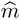) as a function of selection start time. Sweep scenarios consist of hard (*ν* = 1) and soft (*ν* ∈ 2, 4) sweeps with per-generation selection coefficient of *s* = 0.1 that started at *t* ∈ 500, 1000, 1500, 2000, 2500, 3000 generations prior to sampling. Results expanded across wider range of simulation settings can be found in Figures S20–S22 and S26–S28 as well as results for application to unphased multilocus genotype data in Figures S23–S25 and S29–S31.

Obtaining phased haplotypes for input to Λ represents an error-prone step that, without sufficient reference panels or high-enough quality genotypes, may make identification of sweeps difficult or potentially impossible for a number of diverse study systems. It is therefore beneficial if the favorable performance of Λ transfers to datasets that have not been phased. Similar to prior studies [*e.g.*, Harris et al., 2018, Kern and Schrider, 2018, Harris and DeGiorgio, 2020, Harpak et al., 2021], we sought to evaluate the power of Λ when applied to unphased multilocus genotype data, and to compare its performance with the *T* statistic and G123 [analogue of H12 for use with unphased data; Harris et al., 2018], both of which are also applied to unphased multilocus genotypes. Figure S4 shows that Λ maintains high power to detect sweeps of differing ages, strengths, and softness. Consistent with the results on haplotype data (Figures 2A and S1), Λ generally displays higher power than, or comparable power to, *T* and G123, with the best performance deriving from Λ with a small window size of 51 SNPs, and with substantially higher power for old sweeps compared to other approaches. An exception is that for recent (*t ≤* 1000 generations) and highly soft (*ν* = 16) sweeps, using a window size of 101 SNPs for Λ had substantially higher power than using the smaller 51 SNP window. Moreover, for highly soft (*ν* = 16) and ancient (*t ≥* 2000) sweeps with strength *s* = 0.1, the power of Λ is much lower with unphased multilocus genotypes compared to phased haplotypes (compare Figures S1 and S4). Interpretation of 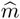 is more difficult for multilocus genotypes compared to haplotypes. However, consistent with the results for haplotypes (Figure S2), Figure S5 shows that when using 51 SNP windows, Λ tends to estimate a small number of sweeping multilocus genotypes (smaller 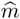) for harder sweeps (smaller *ν*) than for softer sweeps (larger *ν*).

**Figure 4:**
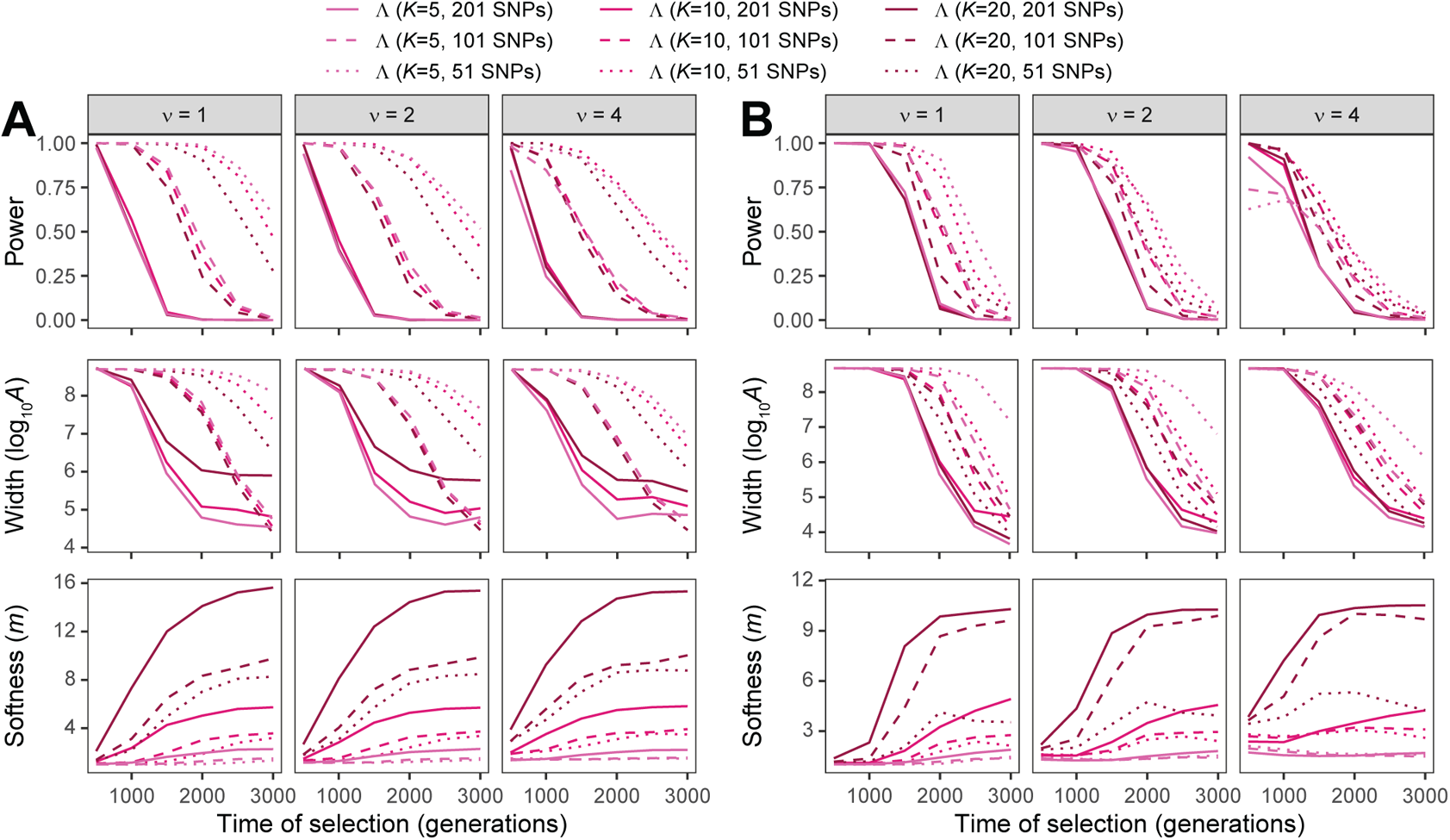
Performance of detecting and characterizing sweeps for applications of Λ with windows of size 51, 101, and 201 SNPs under simulations of (A) a constant-size demographic history or (B) the human central European (CEU) demographic history of Terhorst et al. [2017] and the haplotype frequency spectra for the Λ statistic truncated at *K* 5, 10, 20 haplotypes. Results are based on a sample of *n* = 50 diploid individuals. (Top row) Power at a 1% false positive rate as a function of selection start time. (Middle row) Estimated sweep width illustrated by mean estimated genomic size influenced by the sweep (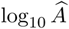) as a function of selection start time. (Bottom row) Estimated sweep softness illustrated by mean estimated number of sweeping haplotypes (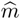) as a function of selection start time. Sweep scenarios consist of hard (*ν* = 1) and soft (*ν* ∈ {2, 4}) sweeps with per-generation selection coefficient of *s* = 0.1 that started at *t* ∈ {500, 1000, 1500, 2000, 2500, 3000} generations prior to sampling. Results expanded across wider range of simulation settings can be found in Figures S32–S34 and S38–S40 as well as results for application to unphased multilocus genotype data in Figures S35–S37 and S41–S43.

**Figure 5:**
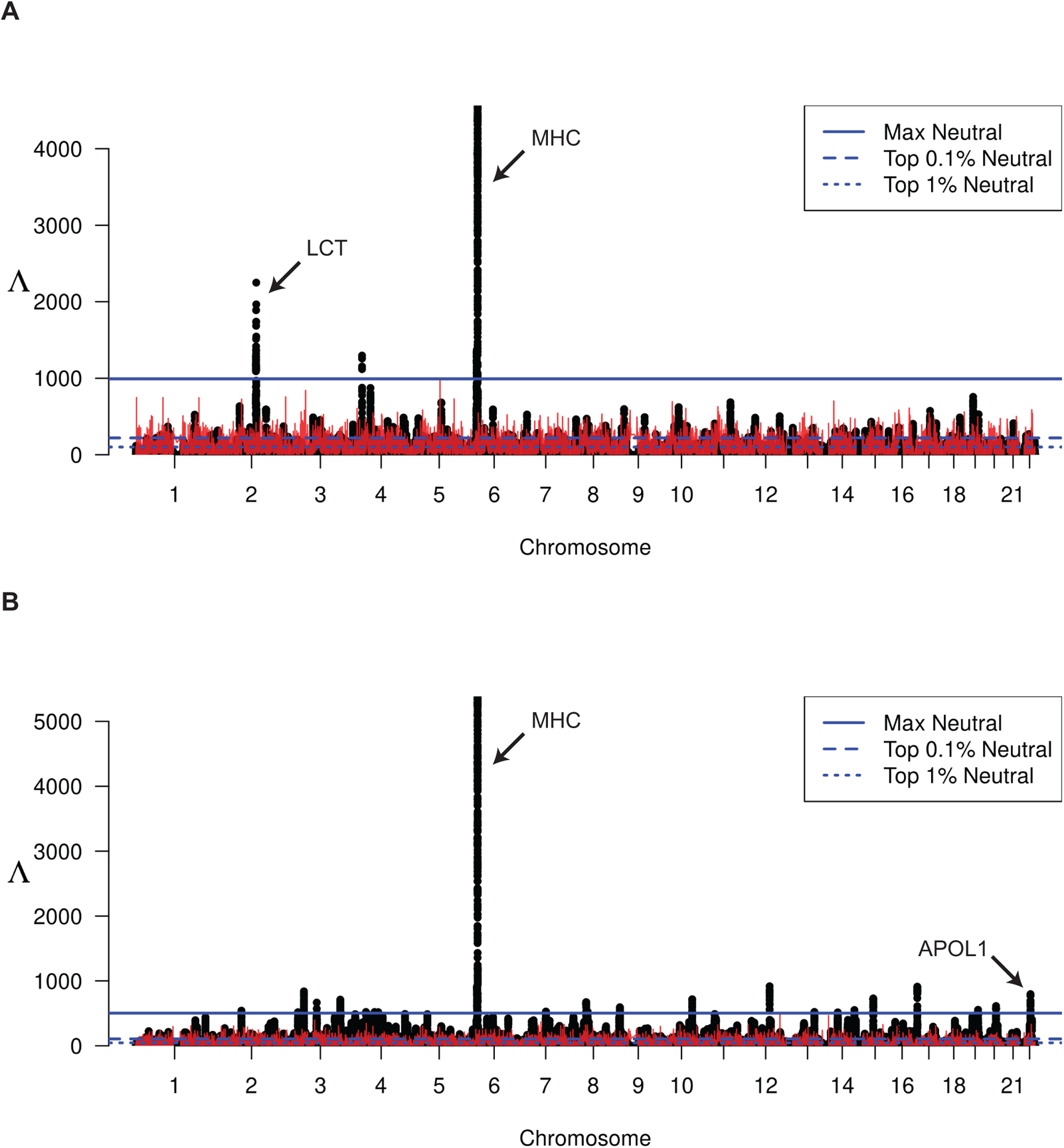
Manhattan plot of Λ-statistics for the (A) CEU and (B) YRI populations from the 1000 Genomes Project. Each point represents a single 201-SNP window along the genome. Horizontal lines represent the top 1%, top 0.1%, and maximum observed Λ statistic across all windows in demography-matched neutral simulations. Red line indicates the maximum observed Λ among 100 replicate simulations at that location in the genome.

While adaptive processes generally affect variation locally in the genome, neutral processes such as demographic history influence overall levels of genome diversity. Specifically, it is common to consider that demographic processes impact the mean value of genetic diversity, and numerous likelihood approaches for detecting sweeps [Kim and Stephan, 2002, Nielsen et al., 2005, Chen et al., 2010, Huber et al., 2015, Vy and Kim, 2015, DeGiorgio et al., 2016, Racimo, 2016, Lee and Coop, 2017, Harris and DeGiorgio, 2020, Setter et al., 2020] and other forms of natural selection [DeGiorgio et al., 2014, Cheng and DeGiorgio, 2019, 2020] have been created to specifically account for this average effect of demographic history on genome diversity. However, demographic processes, such as recent severe bottlenecks, not only alter mean diversity but also influence higher-order moments of diversity, potentially making it insufficient to account solely for the mean effect of diversity [Barton, 1998, Jensen et al., 2005, Pavlidis et al., 2008]. Given that Λ does not account for higher moments than the mean effect of demographic history on the HFS, we sought to evaluate its properties under recent strong bottlenecks—a setting that has proven challenging for other sweep statistics in the past.

The Λ statistic generally exhibits superior power to *T*, H12, *nS_L_*, and iHS when applied to haplotype data (Figures 2B and S7) or to *T* and G123 when applied to unphased multilocus genotype data (Figure S10). Moreover, the general trends in method power as a function sweep strength, softness, and age observed for the constant-size history (Figures 2A, S1, and S4) hold for this complex demographic setting (Figures 2B, S7, and S10), with the caveat that, as expected, power for all methods is generally lower under the bottleneck compared to the constant-size history. A clear difference between these two demography settings is that, whereas Λ had exhibited uniformly superior or comparable power with smaller 51 SNP windows compared to larger 101 or 201 SNP windows (Figures 2A and S1), under the bottleneck model the best window size depends on age of the sweep (Figures 2B and S7). In particular, recent sweeps often had highest power with 201 SNP windows, sweeps of intermediate age with 101 SNPs, and ancient sweeps with 51 SNPs. Therefore, under complex demographic histories, choice of window size for Λ is more nuanced than with constant-size histories. This result is consistent with those of Harris and DeGiorgio [2020] who demonstrated that, when accounting for the spatial distribution of the *T* statistic in a machine learning framework (referred to as *T* -*Trendsetter*), power to detect recent sweeps is higher for larger windows and power to detect ancient sweeps is higher for smaller windows under the bottleneck history considered here.

In addition to demographic history, a pervasive force acting to reduce variation across the genome is background selection [McVicker et al., 2009, Lohmueller et al., 2011, Comeron, 2014, Wilson Sayres et al., 2014], which is the loss of genetic diversity at neutral sites due to negative selection at nearby loci [Charlesworth et al., 1993, Hudson and Kaplan, 1995a, Charlesworth, 2012]. Background selection has been demonstrated to alter the neutral SFS [Charlesworth et al., 1993, 1995, Seger et al., 2010, Nicolaisen and Desai, 2013], and masquerade as false signals of positive selection [Charlesworth et al., 1993, 1995, Hudson and Kaplan, 1995a,b, Nordborg et al., 1996, McVean and Charlesworth, 2000, Boyko et al., 2008, Akashi et al., 2012, Charlesworth, 2012, Huber et al., 2015]. However, because this process does not generally lead to haplotypic variation consistent with sweeps [Enard et al., 2014, Fagny et al., 2014, Schrider, 2020], like prior studies developing haplotype approaches for detecting sweeps [Harris et al., 2018, Harris and DeGiorgio, 2020] we sought to evaluate the robustness of Λ to background selection. We find that under both simple and complex demographic histories, using either phased haplotype or unphased multilocus genotype data, all methods considered here demonstrate robustness to background selection by not falsely attributing genomic regions evolving under background selection as sweeps (Figure S19).

Throughout our experiments, we have considered a per-site per-generation recombination rate of *r* = 10*^−^*^8^ for each simulation replicate. However, recombination rate is known to vary across the genome [Smukowski and Noor, 2011], and it is therefore important to evaluate the performance of Λ compared to other methods when recombination rate varies across genomic regions. To evaluate the effect of recombination rate variation on method performance, we drew per-site per-generation recombination rate from an exponential distribution with mean 10*^−^*^8^ (see *Methods*) for reach replicate neutral and sweep simulation under the bottleneck demographic history [Terhorst et al., 2017]. Figures S13 and S16 indicate that the Λ statistic generally has greater power than *T*, H12 (or G123), *nS_L_*, and iHS under phased haplotypes and unphased multilocus genotypes settings. These results further highlight the robustness of the Λ statistic to realistic genomic characteristics often encountered in empirical studies.

Finally, the number *n* of sampled individuals as well the number *K* of haplotypes used to truncate the HFS should affect the resolution at which we can model the distortion of the HFS due to a sweep, and thus would likely result in alterations of power of Λ to detect sweeps. As expected, Figure 3 shows that increasing sample size generally increases power of Λ to detect sweeps, with highest power typically obtained with the largest *n* and smallest window size combination (*i.e.*, *n* = 50 with 51-SNP windows) and the lowest power with the smallest *n* and largest window size combination (*i.e.*, *n* = 10 with 201-SNP windows). Moreover, as sample size increases, Λ is better able to detect sweeps of older age, and for extremely small samples (*i.e.*, *n* = 10), the estimates 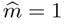 of the number *ν* of sweeping haplotypes are poor. In contrast to changing sample size n, changing the number of haplotypes K to truncate the HFS does not have a substantial effect on the power of Λ to detect sweeps (Figure 4, with the power curves for a specific window size mostly the same across *K ∈ {*5, 10, 20}. This result mirrors that in Figure S5 of Harris and DeGiorgio [2020] for the *T* statistic, whereby changing *K* had little effect on method power. Instead, choice of *K* seems to more strongly influence the estimates 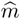 of the number ν of sweeping haplotypes, with larger values of K permitting a wider range of estimates of m. This result mimics those observed for the *T* statistic by Harris and DeGiorgio [2020], in that the choice of *K* has a larger effect on the resolution to classify sweeps as hard or soft than it did on the ability to detect sweeps.

## Application to empirical data

### Humans from the 1000 Genomes Project

The 1000 Genomes Project Phase 3 [The 1000 Genomes Project Consortium, 2015] published the whole genomes of 2504 humans across 26 populations around the world. To illustrate the use of the `saltiLASSI` framework in a context where the populations of interest have well-studied demographic histories, we calculate Λ in two populations: a European population (CEU; *n* = 99) and an African population (*n* = 108). Furthermore, as patterns of recent selection have been extensively studied in these populations, the results will allow us to confirm that the method returns sensible results.

We plot the genome-wide Λ statistics for the CEU population in Figure 5A and the YRI population in Figure 5B. We find several conspicuous peaks of notably large Λ values, which indicates strong support for a highly distorted HFS in these regions compared to the genome-wide mean HFS. We plot the local maximum Λ observed across simulations as a red line, the over-all maximum score (horizontal solid blue line), over-all top-0.1% (horizontal dashed blue line), and over-all top-1% (horizontal dotted blue line); see *Methods* for details.

As this statistic is a composite likelihood ratio test that ignores recombination, we expected that Λ values may be negatively correlated with recombination rate. And, indeed, we find that the max Λ observed in a window across all replicates tends to be larger for low-recombination regions S46. With this in mind, we chose a conservative threshold for determining significance by only calling regions as under selection when the observed Λ is greater than the over-all genome-wide maximum observed Λ from neutral simulations. Taking this approach, we identify several regions in both populations with scores consistently above this threshold, including five regions in the CEU population (Table 1) and 29 in the YRI population (Table 2). Among these regions, we find several well-studied genes that are known to have been under selection in these populations. These include the lactase gene [*LCT*; Tishkoff et al., 2007, Field et al., 2016, Śegurel and Bon, 2017, Taliun et al., 2021], the major histocompatibility complex [MHC; Field et al., 2016, Pierini and Lenz, 2018, Taliun et al., 2021], and the apolipoprotein L1 [*APOL1*; Ko et al., 2013]. We next conduct a gene ontology over-representation test for molecular function using PANTHER16 [Mi et al., 2020] for each population separately. We find that each population’s putatively selected genes are generally representing similar molecular functions (Tables S2 and S3), including MHC class II receptor activity, MHC class II protein complex binding, and peptide antigen binding, further underscoring the evidence for immune system adaptation in human populations around the world [Nédélec et al., 2016, Field et al., 2016, Pierini and Lenz, 2018, Taliun et al., 2021].

**Table 1:** Regions of extreme Λ values in the CEU population and the genes contained therein. 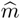 is the inferred number of sweeping haplotypes, and 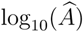 is the estimated sweep width.

| Chr | Start (bp) | Stop (bp) | $\hat{m}$ | $\log_{10}(\hat{A})$ | Max $\Lambda$ | Genes |
| --- | --- | --- | --- | --- | --- | --- |
| 2 | 135,517,106 | 136,318,189 | 1 | 7.817 | 1889.370 | *ACMSD*, *MIR5590*, *CCNT2-AS1*, *CCNT2*, *MAP3K19*, *RAB3GAP1*, *ZRANB3*, *R3HDM1* |
| 2 | 136,524,766 | 136,816,336 | 1 | 7.817 | 2250.050 | *UBXN4*, *LCT*, *LOC100507600*, *MCM6*, *DARS*, *DARS-AS1* |
| 4 | 34,296,435 | 34,400,578 | 1 | 8.252 | 1296.390 | – |
| 6 | 29,782,470 | 29,996,854 | 9 | 7.817 | 1366.550 | *HLA-G*, *HLA-H*, *HCG4B*, *HLA-A*, *HCG9*, *ZNRD1-AS1*, *HLA-J*, *HCG8* |
| 6 | 32,384,933 | 32,723,916 | 7 | 7.817 | 4565.340 | *HLA-DRA*, *HLA-DRB5*, *HLA-DRB6*, *HLA-DRB1*, *HLA-DQA1*, *HLA-DQB1*, *HLA-DQB1-AS1*, *HLA-DQA2*, *MIR3135B*, *HLA-DQB2* |

**Table 2:** Regions of extreme Λ values in the YRI population and the genes contained therein. 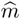 is the inferred number of sweeping haplotypes, and 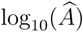 is the estimated sweep width.

| Chr | Start (bp) | Stop (bp) | $\hat{m}$ | $\log_{10}(\hat{A})$ | Max $\Lambda$ | Genes |
| --- | --- | --- | --- | --- | --- | --- |
| 2 | 89,247,854 | 89,309,423 | 8 | 7.817 | 542.403 | – |
| 3 | 27,184,951 | 27,213,780 | 7 | 8.252 | 518.153 | *NEK10* |
| 3 | 46,080,758 | 46,384,651 | 6 | 8.252 | 839.195 | *CCR1, CCR3* |
| 3 | 87,273,225 | 87,328,290 | 6 | 8.252 | 665.802 | *MIR4795, CHMP2B, POU1F1* |
| 3 | 162,536,420 | 162,681,511 | 6 | 8.252 | 711.747 | – |
| 3 | 163,811,348 | 163,832,931 | 7 | 8.252 | 515.029 | – |
| 4 | 46,828,990 | 46,881,572 | 8 | 8.252 | 521.042 | *COX7B2* |
| 4 | 74,441,768 | 74,503,196 | 7 | 8.252 | 520.893 | *RASSF6* |
| 4 | 86,144,358 | 86,185,751 | 9 | 8.252 | 517.750 | – |
| 6 | 31,193,373 | 31,387,666 | 11 | 7.817 | 918.650 | *HLA-C, HLA-B, MIR6891, MICA* |
| 6 | 32,370,521 | 32,750,144 | 8 | 7.817 | 5385.050 | *BTNL2, HLA-DRA, HLA-DRB5, HLA-DRB6, HLA-DRB1, HLA-DQA1, HLA-DQB1, HLA-DQB1-AS1, HLA-DQA2, MIR3135B, HLA-DQB2* |
| 6 | 32,973,878 | 33,206,733 | 8 | 7.817 | 1196.890 | *HLA-DOA, HLA-DPA1, HLA-DPB1, HLA-DPB2, COL11A2, RXRB, SLC39A7, HSD17B8, MIR219A1, RING1* |
| 7 | 80,311,522 | 80,335,123 | 7 | 8.252 | 525.732 | – |
| 7 | 80,335,124 | 80,390,838 | 7 | 8.252 | 527.827 | *SEMA3C* |
| 8 | 50,054,576 | 50,219,674 | 6 | 8.252 | 672.218 | – |
| 8 | 54,726,028 | 54,770,306 | 7 | 8.252 | 526.734 | *ATP6V1H, RGS20* |
| 9 | 11,767,302 | 11,832,748 | 8 | 8.252 | 594.456 | – |
| 10 | 102,156,400 | 102,295,419 | 5 | 8.252 | 717.760 | *WNT8B, SEC31B, NDUFB8* |
| 12 | 79,566,706 | 79,800,343 | 6 | 8.252 | 917.585 | *SYT1* |
| 13 | 89,191,303 | 89,233,281 | 8 | 8.252 | 521.733 | *LINC00433* |
| 14 | 48,798,901 | 48,833,362 | 8 | 7.817 | 504.040 | – |
| 14 | 48,833,363 | 48,853,450 | 7 | 7.817 | 515.715 | – |
| 14 | 102,173,879 | 102,216,165 | 7 | 7.817 | 551.532 | *LINC00239* |
| 15 | 55,137,170 | 55,292,403 | 7 | 8.252 | 730.824 | – |
| 17 | 3,515,275 | 3,672,429 | 6 | 8.252 | 911.490 | *SHPK, CTNS, TAX1BP3, P2RX5-TAX1BP3, EMC6, P2RX5, ITGAE, GSG2* |
| 19 | 38,859,266 | 38,939,066 | 7 | 8.252 | 555.198 | *CATSPERG, PSMD8, GGN, SPRED3, FAM98C, RASGRP4, RYR1* |
| 19 | 39,176,160 | 39,201,469 | 7 | 8.252 | 504.612 | *ACTN4* |
| 20 | 37,392,189 | 37,490,122 | 5 | 8.686 | 613.034 | *ACTR5, PPP1R16B* |
| 22 | 36,567,890 | 36,756,255 | 6 | 8.252 | 796.478 | *APOL4, APOL2, APOL1, MYH9, MIR6819* |

We next explore two peaks in detail, the *LCT* and MHC loci (Figure 6), to illustrate the spatial structure of the HFS in these regions of strong signal in one (*LCT*) or both (MHC) populations. The *LCT* locus has been previously identified as under selection in some northern European populations and eastern African populations [Tishkoff et al., 2007]. As the CEU population has largely northern European ancestry and the YRI population is from western Africa, we expect to find a peak near *LCT* in CEU but not in YRI. Indeed, this is what we see in Figure 6A, which plots Λ statistics in the vicinity of the *LCT* locus on Chromosome 2. Furthermore, we examine the truncated HFS among eleven windows spanning *LCT* in both YRI (Figure 6B) and CEU (Figure 6C). We see in Figure 6B that YRI has haplotype frequencies similar to the genome-wide mean (plotted and highlighted on the left), whereas Figure 6C shows that the CEU population is dominated largely by a single haplotype near 80% frequency. Indeed, the `saltiLASSI` method also infers a 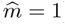 in this region (Table 1), indicating a single sweeping haplotype (i.e., a hard sweep). Furthermore, we can see the HFS in this region trending toward the genome-wide mean as the windows move farther from the sweep’s focal point, illustrating the pattern that the `saltiLASSI` method was designed to capture.

**Figure 6:**
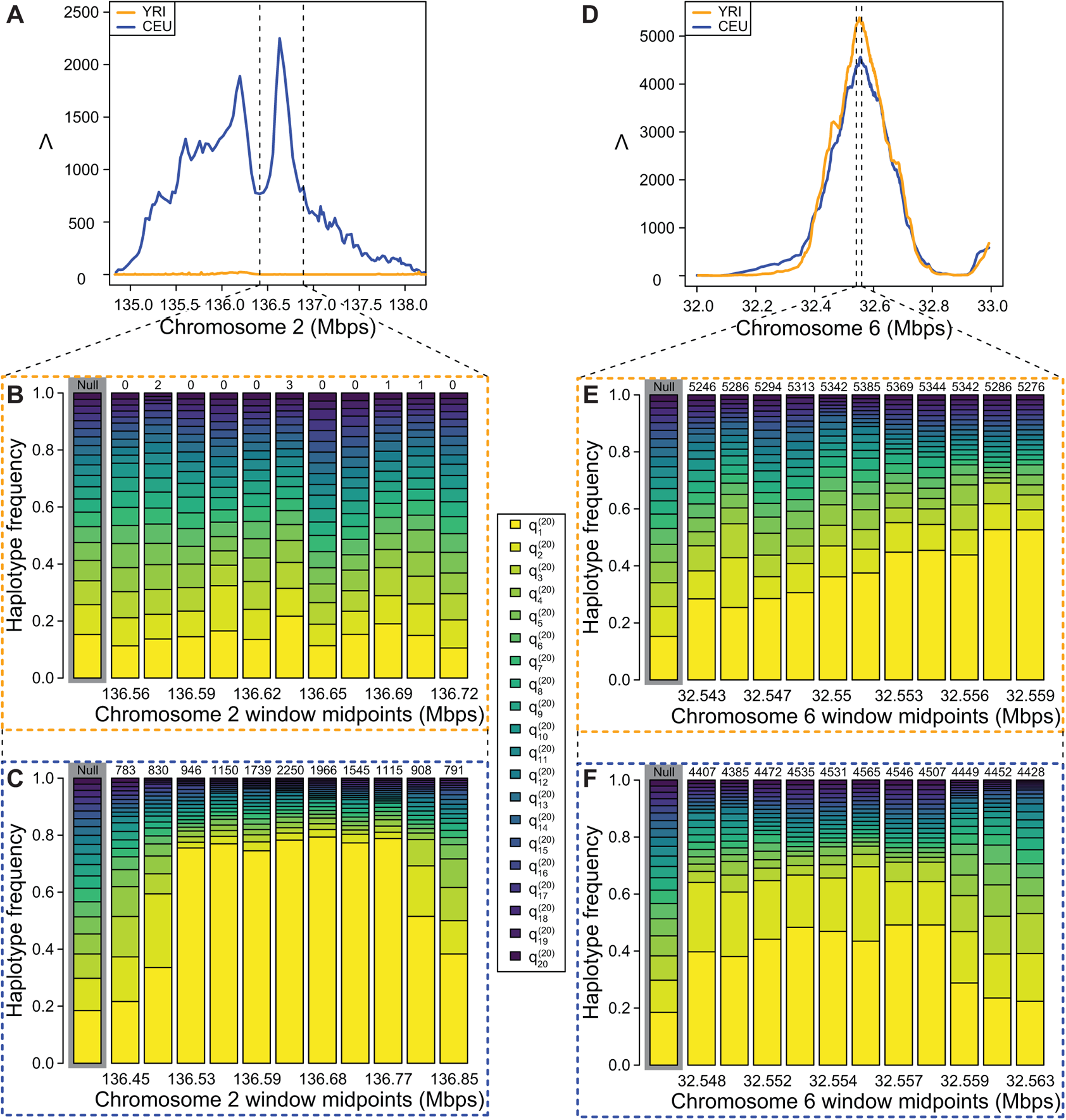
Detailed illustration of Λ statistics and haplotype frequency spectra in CEU and YRI. (A) Λ plotted in the *LCT* region, vertical dotted lines indicate zoomed region shown in (B) and (C). (B) YRI empirical HFS for 11 windows in the *LCT* region. (C) CEU empirical HFS for 11 windows in the *LCT* region. (D) Λ plotted in the MHC region, vertical dotted lines indicate zoomed region shown in (E) and (F). (E) YRI empirical HFS for 11 windows in the MHC region. (F) CEU empirical HFS for 11 windows in the MHC region. In (B), (C), (E), and (F), numbers above HFS are Λ values for the window rounded to the nearest whole number, and the genome-wide average HFS is highlighted in grey. 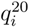 is the frequency of the *i*th most common haplotype truncated to *K* = 20.

Figures 6D-F illustrate the Λ statistics and HFS patterns in the vicinity of the MHC locus. This locus contains a large cluster of immune system genes, and selection at this locus is distinguished from *LCT* in that high diversity is preferred in order for the body to be able to mount a robust response to unknown pathogen exposure. As expected, both populations have extreme Λ values (Figure 6D) and a greatly distorted HFS in this region (Figures 6E and F). However, we note that the HFS is clearly distorted in favor of multiple haplotypes, in contrast to *LCT*, which we expect at a locus that favors diversity. Indeed, the `saltiLASSI` method infers 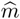 to be between seven and nine in the CEU population and between eight and 11 for YRI (variance due to multiple regions within the MHC being separately identified; Tables 1 and 2).

We repeated our analyses of these two populations and two loci using the unphased multilocus-genotype approach (Figures S44 and S45; Tables S5 and S6), and we find good concordance with the phased haplotype approach.

Finally, we re-compute Λ (phased) in these two populations’ empirical data and all replicates of simulated demography-matched whole-genome data using two distance measures other than physical distance (number of windows and centiMorgans) and find high correlation between Λ values calculated with these alternative distance measures and physical distance (Table S4).

### Rats from New York City

Harpak et al. [2021] published a whole-genome dataset of brown rat samples (*n* = 29) from across the island of Manhattan, New York City, USA to study adaptation to urban environment. In this study, they note that haplotype phase is unknown and that the demographic history for brown rats was not well-calibrated in this population. As such, they chose to use the G123 [Harris et al., 2018] and other statistics, which used multilocus genotypes combined with a gene-based outlier approach to identify putative targets of selection. Here, we re-analyze this data using the `saltiLASSI` framework to illustrate its use in the context of unphased data and a poorly understood demographic history that requires an outlier approach. We plot the genome-wide Λ statistics for the NYC rats in Figure 7, along with blue horizontal lines indicating the top 0.1% (solid), top 1% (dashed), and top 5% (dotted) empirically observed Λ values genome-wide. We identify putatively selected regions as windows with a Λ greater than the top 1% empirical threshold (see *Methods*), with consecutive windows satisfying this condition concatenated together. These regions are then annotated with known genes (RN5 genome build) and presented in Tables S7 and S8.

**Figure 7:**
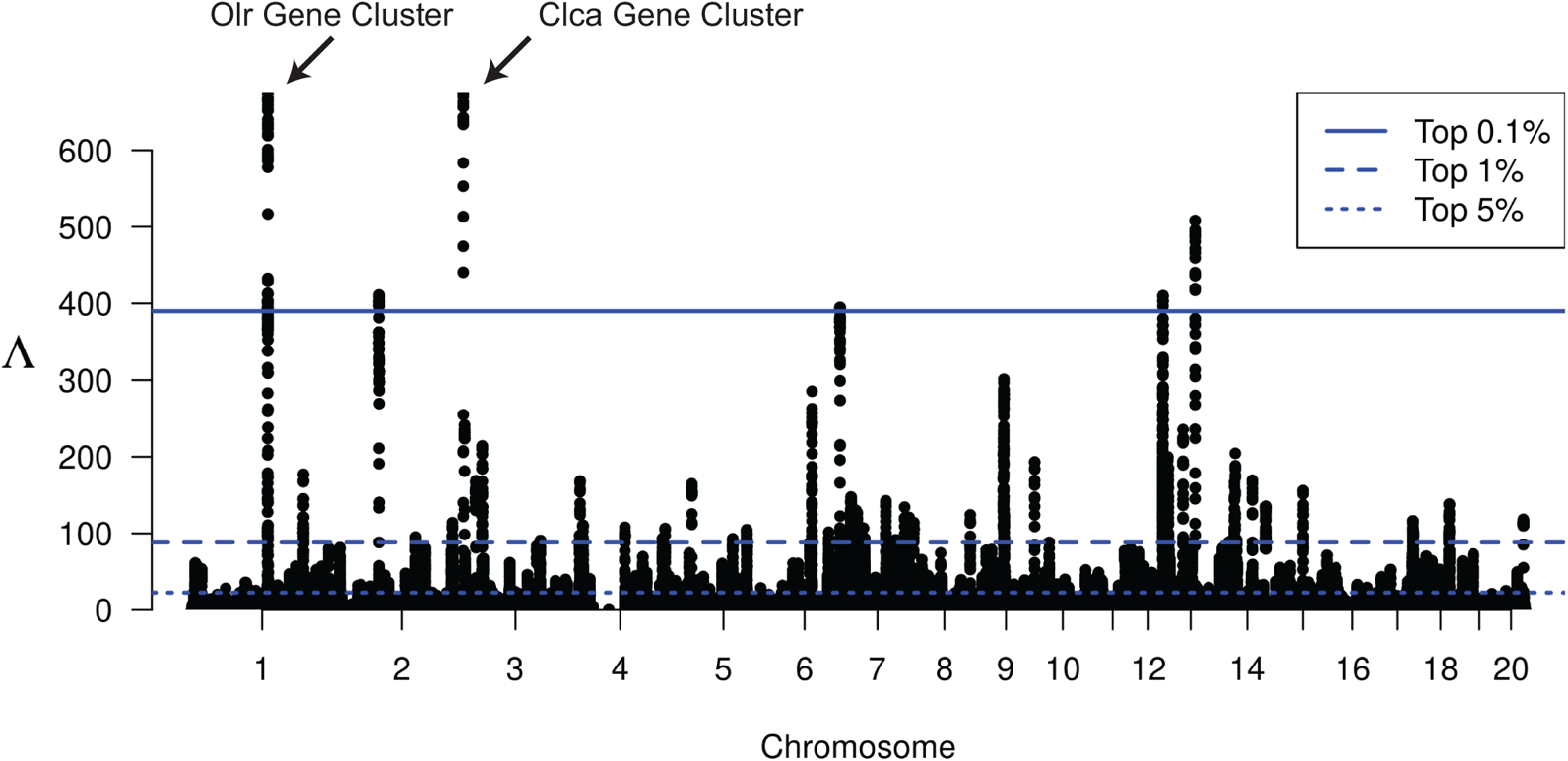
Manhattan plot of Λ-statistics for the New York City rat population. Each point represents a single 201-SNP window along the genome. Horizontal lines represent the top 5%, top 1%, and top 0.1% observed Λ statistic across all windows in the genome.

We note that the two strongest signals in the genome are on chromosomes 1 and 2 (Table S7). The region on chromosome 1 contains a cluster of olfactory receptor genes (*Olr23*, *Olr24*, *Olr25*, *Olr27*, *Olr29*, *Olr30*, *Olr32*, and *Olr34*), and the region on chromosome 2 contains a cluster of calcium-activated chloride channel genes (*Clca2*, *Clca4l*, *Clca4*, *Clca1*, and *Clca5*). Notably, calcium-activated chloride channel genes are expressed in the olfactory nerve layer of mouse brains [Piirsoo et al., 2009]. If these calcium-activated chloride channel genes are similarly expressed in rats, then these two strong selection signals suggest that this urban rat population may be experiencing selection pressures associated with olfactory perception.

Taking the collection of annotated genes present in Table S7, we conduct a gene ontology over-representation test based on molecular function category using PANTHER16 [Mi et al., 2020] with results presented in Table 3. We find that Intracellular Calcium Activated Chloride Channel Activity, Peptidase Activity, and Odorant Binding are statistically over-represented molecular functions among this set of putatively selected genes.

**Table 3:** Gene ontology enrichment analysis of regions with extreme Λ values in the New York City rat population.

| GO molecular function | Fold Enrichment | Raw P-value | FDR |
| --- | --- | --- | --- |
| Intracellular calcium activated chloride channel activity | > 100 | $5.16 \times 10^{-12}$ | $1.23 \times 10^{-8}$ |
| Intracellular chloride channel activity | > 100 | $5.16 \times 10^{-12}$ | $2.46 \times 10^{-8}$ |
| Chloride channel activity | 45.49 | $5.81 \times 10^{-9}$ | $6.93 \times 10^{-6}$ |
| Anion channel activity | 39.07 | $1.37 \times 10^{-8}$ | $1.31 \times 10^{-5}$ |
| Inorganic anion transmembrane transporter activity | 24.06 | $2.14 \times 10^{-7}$ | $1.70 \times 10^{-4}$ |
| Inorganic molecular entity transmembrane transporter activity | 6.39 | $2.96 \times 10^{-5}$ | $9.42 \times 10^{-3}$ |
| Anion transmembrane transporter activity | 11.30 | $1.51 \times 10^{-5}$ | $6.53 \times 10^{-3}$ |
| Ion transmembrane transporter activity | 5.47 | $8.71 \times 10^{-5}$ | $2.60 \times 10^{-2}$ |
| Ion channel activity | 9.07 | $1.10 \times 10^{-5}$ | $5.27 \times 10^{-3}$ |
| Channel activity | 8.12 | $2.23 \times 10^{-5}$ | $7.60 \times 10^{-3}$ |
| Passive transmembrane transporter activity | 8.12 | $2.23 \times 10^{-5}$ | $8.19 \times 10^{-3}$ |
| Ion gated channel activity | 69.19 | $5.57 \times 10^{-10}$ | $8.88 \times 10^{-7}$ |
| Gated channel activity | 11.60 | $2.27 \times 10^{-6}$ | $1.21 \times 10^{-3}$ |
| Metallopeptidase activity | 16.86 | $1.60 \times 10^{-6}$ | $9.56 \times 10^{-4}$ |
| Peptidase activity | 6.92 | $1.68 \times 10^{-5}$ | $6.70 \times 10^{-3}$ |
| Odorant binding | 10.06 | $1.12 \times 10^{-6}$ | $7.62 \times 10^{-4}$ |

## Discussion

In this study, we developed a new likelihood ratio test statistic Λ that examines the spatial distribution of the HFS for evidence of sweeps. We demonstrated that this statistic has high power to detect both hard and soft sweeps, with performance substantially better than competing haplotype-based approaches for the same task. Moreover, while optimizing the model parameters of Λ we obtain estimates of sweep softness *m* and footprint size *A*, which is correlated with age and strength of the sweep. These additional parameters have the potential to further characterize well-supported sweep signals from large Λ values.

In addition to lending exceptional performance on simulated data, application of Λ to whole-genome variant calls from central European and sub-Saharan African individuals recapitulated the well-established signal at the *LCT* gene in Europeans due to lactase persistence [Bersaglieri et al., 2004], as well as sweep footprints at the MHC locus in both populations related to immunity, which have previously been detected with other sweep statistics [Albrechtsen et al., 2010, Goeury et al., 2018, Harris and DeGiorgio, 2020]. Though not novel findings, the clear (Figure 6) and strong (Figure 5) signals at these two loci serve as positive controls to highlight the efficacy of Λ. Furthermore, these findings were similarly recapitulated with unphased multilocus genotype data (Figures S44 and S45), lending support for the utility of Λ when applied to study systems for which obtaining phased haplotypes data is challenging.

Though our identification of the MHC locus in both human empirical scans as a sweep is not novel, it is important to address that the MHC locus comes with a number of technical challenges when assessing genetic variation. Specifically, the MHC locus is known to harbor extensive structural variation, which makes it difficult to assemble [Dilthey et al., 2015] and may lead to downstream errors in variant and genotype calling and in haplotype phasing. Indeed, such difficult to assemble regions may lead to enrichment in heterozygous sites, where in the extreme the majority of individuals are heterozygous. Contiguous SNPs in which individuals have heterozygous genotypes may manifest as a single high-frequency unphased multilocous genotype that stems from two distinct and divergent high-frequency haplotypes. Because Λ only considers the frequency of haplotypes and multilocus genotypes, it may lend support for sweeps in regions where genetic variation is difficult to assay. As with any other sweep detection approach, we recommend that care be taken when pre- and post-processing genomic datasets to attempt to circumvent these issues whenever possible, such as filtering regions with poor mappability, as we have done in this study (see *Methods*).

As the human populations have a well-characterized demographic history, we were able to perform demography-matched neutral simulations to aid in identifying regions of the genome likely affected by selection. When analyzing the New York City brown rat dataset, we had to take an outlier approach as the brown rat demographic history was previously noted to be mis-calibrated for this population [Harpak et al., 2021]. However, our outlier approach notably identified two strong signals of selection among clusters of genes related to olfactory perception. As rats depend heavily on scents for communication and behavior choices [Parmiani et al., 2018, Parsons et al., 2018, 2019], it is reasonable to think that a harsh, noisy, urban environment may present selection pressure on this biological system.

A key parameter that must be chosen when applying Λ is the number of SNPs per window. Specifically, we found that larger windows had greatest power for more recent sweeps, and smaller windows for more ancient sweeps (Figures 2, S1, and S7), mirroring the window size results observed in Figures S8 and S9 of Harris and DeGiorgio [2020] for the spatial distribution of the *T* statistic using a different modeling approach. Therefore, choice of window size may be informed by the time frame of selective events that is being investigated. As highlighted in Figures 2B and S7, the Λ statistic computed within windows of 201 SNPs had highest power of all other tested window sizes within the past 1500 generations under the central European demographic history. Because selective events within this time frame are consistent with adaptive events in recent evolution of modern humans [Gravel et al., 2011, Gronau et al., 2011, Schiffels and Durbin, 2014], we selected this size so that we could recapitulate expected well-established sweeps— *e.g.*, Figures 5 and 6 highlighting the sweep signal at *LCT*. In addition to using simulation results to aid in selecting appropriate window sizes, an alternate method such as choosing sizes based on the expected decay of linkage disequilibrium in the genome has been demonstrated to also work well in practice [*e.g.*, Garud et al., 2015, Harris and DeGiorgio, 2020].

We note that this approach is a composite likelihood statistic, and as such it treats windows as independent, ignoring the effects of recombination. This means that Λ values are likely to be larger in low-recombination regions (Figure S46), and extreme scores found in such regions should be treated with extra scrutiny. However, even in such regions, we have shown that one can employ a simulation based approach to evaluate the uncertainty in the estimated Λ values (Figure 5 and S44)—albeit such an approach can be computationally intensive and would require accurate demographic model and recombination map estimates. An alternative solution to evaluate the uncertainty in Λ while also accounting for recombination would be to perform a block resampling locally in the genome [Lieu and Singh, 1992]. Such an approach would prove valuable for study systems without accurate estimates of demographic models and recombination maps, and would provide an alternative uncertainty metric even for organisms such as humans for which simulations can be employed to evaluate uncertainty.

The *T* statistic of Harris and DeGiorgio [2020] presented the first likelihood approach that evaluated distortions in the HFS to detect selective sweeps, importantly because neutrality and soft sweeps leave similar signatures in the SFS but different within the HFS [Pennings and Hermisson, 2006b]. As demonstrated by Harris and DeGiorgio [2020], using the spatial distribution of the *T* statistic within a machine learning framework enhanced its detection ability, specifically for ancient sweeps. However, machine learning frameworks require extensive simulations to train [*e.g.*, Schrider and Kern, 2016, Sheehan and Song, 2016, Mughal and DeGiorgio, 2019], and these simulations must be based on a set of critical assumptions, such as demographic, mutation rate, and recombination rate parameters. Yet, accurate inferences of these parameters is not always possible, or can be highly error prone, and prior studies have found that these machine learning methods can make highly incorrect predictions if the distribution of training data is different from that of the test or empirical data [Mughal and DeGiorgio, 2019, Mughal et al., 2020]. Furthermore, generation of these training datasets and training the models on them often requires substantial computational time and resources. Instead, our Λ statistic is the first likelihood method to model the spatial distribution of the HFS, providing the power of modeling the spatial distribution of *T* afforded by current machine learning frameworks (*e.g*., compare Figures S1 and S7 with Figures S8 and S9 of Harris and DeGiorgio [2020]). This power comes without having to simulate over a broad range of parameters to train a model, thus saving computational resources, and with predictions not hinging on accurate estimates of genetic and evolutionary model parameters to generate training sets. However, this high power of the Λ statistic to detect candidate sweep regions without simulations is distinct from the requirement that distributions of the statistic from neutral simulations must be generated to reject neutrality at candidate sweep regions. Any sweep statistic, regardless of it being a summary, likelihood, or machine learning approach will require extensive simulations under realistic genetic and evolutionary models to reject the null hypothesis of neutrality.

While optimizing the Λ statistic, we also obtain estimates of the number of presently-sweeping haplotypes *m* and the footprint size *A*. For recent sweeps that are strong enough, estimates of *m* correlate well with the number of initially-selected haplotypes *ν*. For older and less strong sweeps, mutation and recombination events accumulate leading to more distinct haplotypes, thereby inflating *m* estimates. Moreover, estimates of the footprint size *A* correlate with power of Λ, suggesting that the estimated footprint size will be large under scenarios in which sweeps are highly supported. The relationship between *A* and power of Λ is related to prominence of the distortions in the HFS, which also erode due mutation and recombination rates, and this parameter is analogous to the *α* parameter [Durrett and Schweinsberg, 2004] used by other composite likelihood methods to mechanistically model the probability that a lineage escapes a sweep [Nielsen et al., 2005, Setter et al., 2020]. Therefore, though we found that estimates of *m* were not highly accurate under non-ideal sweep settings and that the precise relationship of *A* to the timing and strength of a sweep is unclear, these quantities may still be useful. Specifically, even if the estimates of *m* are not highly accurate proxies for *ν*, estimates of *m* could still be valuable by casting the problem as binary sweep classification with *m* = 1 for hard and *m >* 1 for soft sweeps, as was also suggested for the *T* statistic by Harris and DeGiorgio [2020]. Table 1 highlights that the *LCT* region is identified as a hard sweep (estimated *m* = 1) in the CEU, with inferred soft sweeps (estimated *m >* 1) in the MHC region, which are consistent with the number of prominent high-frequency haplotypes at these regions (Figures 6). Moreover, though not directly associated with population-genetic parameters such as *ν* or the strength *s* and time *t* of a sweep, estimated Λ, *µ*, and *A* values can be used as input features to machine learning regression algorithms to predict underlying evolutionary model parameters of *ν*, *s*, and *t* [Hastie et al., 2009]. Such strategies are typically computationally expensive, but may be required for accurate characterization of sweep footprints, even though they are unnecessary for detecting sweeps due to the already high power of Λ.

The Λ statistic developed here represents an important step in advancing methodology for sweep detection by interrogating the spatial distribution of distortions in the HFS. Prior studies focused either on spatial distributions of the SFS, which cannot distinguish between hard and soft sweeps, or only local distortions in the HFS. Specifically, methods that explore the skews in the SFS typically do so with an explicit analytical population-genetic model [Kim and Stephan, 2002, Nielsen et al., 2005, Huber et al., 2015, Vy and Kim, 2015, DeGiorgio et al., 2016], which are underpowered if the assumed model is incorrect and are underpowered to detect soft sweeps [Pennings and Hermisson, 2006b]. In contrast, analytical population-genetic modeling of distortions in the HFS is difficult, and alternative statistical models that capture relevant features of sweeps are often used, focusing either on local distortions in the HFS [Harris and DeGiorgio, 2020] or haplotype length distributions [Voight et al., 2006, Ferrer-Admetlla et al., 2014]. Instead, our Λ statistic represents a compromise of these two extremes, permitting simultaneous interrogations of haplotype frequency distributions and correlates of their length distributions in a computationally efficient framework that leads to expected patterns that are informed by theoretical results. Our methodological framework therefore provides a foundation for developing tools that can identify other evolutionary processes that may act locally in the genome, enhancing future investigations of sweeps and other forces across a variety of study systems.

## Methods

In this section we outline the methods used to assess the power of a diversity of sweep statistics using simulations. These simulations examine an array of model parameters, including sweep strength, age, and softness as well as the confounding effects of demographic history, background selection, haplotype phasing, and recombination rate variation. We also describe pre- and post-analysis processing for the application of the Λ statistic to our two real-data examples: CEU and YRI human populations and a rat population from New York City.

### Power Analysis

To assess the ability of Λ to detect sweeps, we conducted forward-time simulations using `SLiMv3.2` [Haller and Messer, 2019] for sweeps of varying strength, age, and softness under a constant-size demographic history as well as under a realistic non-equilibrium demographic history inspired by human studies. Specifically, for each simulation scenario, we generated 1000 independent replicates of length 500 kb, so that Λ was able to interrogate the spatial distribution of variation across a large genomic segment. We employed a mutation rate of *µ* = 1.29 *×*10*^−^*^8^ per site per generation [Scally and Durbin, 2012, Adrion et al., 2020] and a recombination rate of *r* = 10*^−^*^8^ per site per generation [Payseur and Nachman, 2000]. For the constant-size demographic history, we considered a population size of *N* = 10^4^ diploid individuals [Takahata, 1993], and to investigate complex non-equilibrium demographic histories, we employed the model inferred in Terhorst et al. [2017] of central European humans (CEU), which incorporates a recent bottleneck with a severe population collapse followed by rapid population expansion. In particular, we used this non-equilibrium model as it was inferred by the contemporary method `SMC++` [Terhorst et al., 2017], which attempts to fit model parameters that can both recapitulate haplotype diversity and allele frequency distributions [Beich-man et al., 2017] observed in genomic data from the CEU population of the 1000 Genomes Project dataset [The 1000 Genomes Project Consortium, 2015]. We also considered a setting in which recombination rate was permitted to vary across simulation replicates under the CEU demographic model, with recombination rate for a given simulated replicate drawn from an exponential distribution with mean *r* = 10*^−^*^8^ per site per generation [*i.e.*, inspired by Schrider and Kern, 2016].

In addition to these genetic and demographic parameters, for selection simulations, we modeled sweeps on *ν ∈ {*1, 2, 4, 8, 16} initially-selected haplotypes, where each of these haplotypes harbored a beneficial allele in the center of the simulated genomic segment with strength *s ∈ {*0.001, 0.01, 0.1} per generation that immediately appeared and became beneficial at time *t ∈ {*500, 1000, 1500, 2000, 2500, 3000} generations prior to sampling. To ensure that a sweep signature had the potential to be uncovered (especially under settings with *s* = 0.001 and 0.01), we required that the beneficial allele established in the population by reaching a frequency of 0.1 in the population. Simulation replicates for which the beneficial allele did not reach a frequency of 0.1 in the population were repeated until the beneficial allele established in the population. All neutral and selection simulations were run for 11*N* generations, where the first 10*N* generations were used as burn-in and *n* = 50 diploid individuals were sampled from the population after 11*N* generations (*i.e.*, the present). Because forward-time simulations are computationally intensive, as is commonly-practiced [Yuan et al., 2012, Ruths and Nakhleh, 2013] we scaled all constant-size demographic history simulations by a factor *λ* = 10 and the European human history by *λ* = 20, such that the selection coefficient, mutation rate, and recombination rate were multiplied by *λ* and the population size at each generation and the total number of simulated generations were divided by *λ*. This scaling leads to a speedup of approximately *λ*^2^ in computing time, such that the constant-size simulations run roughly 100 times faster than without scaling and the CEU model simulations run approximately 400 times faster, making a large-scale simulation study feasible.

When analyzing each simulated replicate, we examined the performance of Λ with the likelihood *T* statistic [Harris and DeGiorgio, 2020] that does not account for the spatial distribution of genomic variation, the summary statistic H12 [Garud et al., 2015] that was developed to detect hard and soft sweeps with similar power, and the standardized iHS [Voight et al., 2006] and *nS_L_* Ferrer-Admetlla et al. [2014] methods that summarize the lengths of haplotypes centered on core SNPs. When applying one of these sweep detection statistics to a simulated replicate, we scanned the entire simulated region, and the score of the applied statistic for that simulated replicate was chosen as the maximum value of that statistic, computed across all test positions within the simulated region. To investigate the effect of window size on the relative powers of Λ, *T*, and H12, we considered their applications in central windows of 51, 101, and 201 SNPs, and analyzed windows every 25 SNPs across a simulated sequence. We chose SNP-delimited windows rather than windows based on physical length as they should be more robust to variation in recombination and mutation rate across the genome, as well as random missing genomic segments due to poor mappability, alignability, or sequence quality. That is, we expect SNP-delimited to be more conservative than windows based on the physical length of an analyzed genomic segment. We also examined the application of Λ, *T*, and G123 [Harris et al., 2018, analogue of H12] to unphased multilocus genotype input data to evaluate the relative powers of these three approaches when applied on study systems for which obtaining phased haplotypes is difficult, unreliable, or impossible [Mallick et al., 2009]. We applied the `lassip` software released with this article for application of the `saltiLASSI` Λ statistic, the `LASSI` *T* statistic, and H12 (and G123), and the `selscan` software [Szpiech and Hernandez, 2014] to compute standardized iHS and *nS_L_*.

### Analysis of 1000 Genomes Data

We extracted the phased genomes of CEU (99 diploids) and YRI (108 diploids) populations, separately, from the full 1000 Genomes Project Phase 3 dataset (2504 diploids) [The 1000 Genomes Project Consortium, 2015]. For each population, we retained only autosomal biallelic SNPs that were polymorphic in the sample. In order to avoid potentially spurious signals, we also filtered any regions with poor mappability as indicated by mean CRG100 *<* 0.9 [Derrien et al., 2012, Huber et al., 2015]. This left 12,400,078 SNPs in CEU and 20,417,698 SNPs in YRI.

We compute `saltiLASSI` Λ statistics for both phased (haplotype-based) and unphased (multilocus-genotype-based) analyses with `lassip`. We use physical distance as the distance measure, and we set `--winsize 201`, `--winstep 100`, and `--k 20` to use the ranked HFS for the top *K* = 20 most frequent haplotypes. By default `lassip` assumes phased data and computes haplotype-based statistics, when the `--unphased` flag is set, all statistics are computed using multilocus genotypes.

To determine significance thresholds, we simulated neutral whole genomes with a realistic recombination map and demographic history using `stdpopsim` [Adrion et al., 2020] and `msprime` [Kelleher et al., 2016]. Using the `OutOfAfrica` `2T12` demographic history [Tennessen et al., 2012] and the `HapMapII` `GRCh37` genetic map [Consortium, 2007] in `stdpopsim`, we simulate 100 replicates of all 22 autosomes for each population separately, sampling 99 diploid individuals for CEU simulations and 108 diploid individuals for YRI simulations. For each replicate, we then compute `saltiLASSI` Λ statistics for both phased and unphased analyses with `lassip`, setting `--winsize 201`, `--winstep 100`, and `--k 20`. As simulated genomes do not simulate variants at the same sites, the windows within which Λ is calculated will not perfectly align with each other or our real-data analysis. In order to compare neutral and real Λ values at local regions of the genome, for each neutral replicate, separately, we align the simulated windows to the windows of our real-data analysis, and then for each real-data window we calculate a weighted mean of all overlapping windows to get a neutral-simulation Λ for that window associated with our real-data. In this way we are able to compute 100 neutral-simulated Λ values for each window in our real-data analyses. We then compute the max Λ, the top-0.1% Λ, and the top-1% Λ across all windows in all replicates for each population and each analysis (phased/unphased), which are given in Table S1. We consider any window with a Λ greater than the max observed across all genome analysis windows from all neutral simulations as a putatively selected region, and we concatenate consecutive windows satisfying this condition into larger regions implicated as being under selection (phased in Tables 1 and 2 and unphased in Tables S5 and S6).

Finally, we also compute Λ for all simulated and empirical data using two other distance measures: number of windows and centiMorgans. For the latter measure we use the `HapMapII` `GRCh37` genetic map [Consortium, 2007] and use the genetic distance between window midpoints. Midpoints for which a genetic position does not exist in the `HapMapII` `GRCh37` genetic map are linearly interpolated based on the nearest surrounding sites. We compare these results to the results calculated using physical distance using Spearman’s rank correlation (Table S4). For simulated data, we compute the mean correlation coefficient across all 100 replicates.

### Analysis of New York City Rats

We extracted the genetic data of 29 rats sampled in New York City [Harpak et al., 2021], retaining only autosomal biallelic SNPs that were polymorphic in the sample. This left 13,532,711 SNPs. As these data are unphased, we use `lassip` to compute `saltiLASSI` Λ statistic using multilocus-genotypes (`--unphased` flag). We set `--winsize 201` and `--winstep 100`, and we choose `--k 20` to use the ranked HFS for the top *K* = 20 most frequent haplotypes.

Harpak et al. [2021] noted that the demographic history for brown rats was likely poorly calibrated for these New York City samples. We therefore take an outlier approach for analyzing the results of the `saltiLASSI` method on these data. We compute the top-0.1% Λ, the top-1% Λ, and the top-5% Λ across all windows genome-wide, getting 389.839, 88.080, and 22.724, respectively. Putatively selected regions were identified by concatenating consecutive windows with Λ greater than the top-1% Λ observed (Tables S7 and S8).

## Data Availability

Software implementing this method is available at https://github.com/szpiech/lassip. The 1000 Genomes Project data is available at https://www.internationalgenome.org/, and the New York City rat data is available at https://doi.org/10.5061/dryad.08kprr4zn. Analysis scripts and intermediate data files used in this study are available from Data Dryad at doi:10.5061/dryad.4qrfj6qbm.

## Acknowledgments

We thank three anonymous reviewers for their constructive feedback that helped strengthen this manuscript. This work was supported by National Institutes of Health grant R35GM128590, by National Science Foundation grants DBI-2130666, DEB-1949268, and BCS-2001063, and by Pennsylvania State University startup funds. Computations for this research were performed using the services provided by Research Computing at the Florida Atlantic University and using the Pennsylvania State University’s Institute for Computational Data Sciences’ Roar supercomputer.

## Supplementary material

**Table S1:** Λ statistic thresholds for TGP analyses as calculated from demography-matched neutral simulations.

| Population | Phased | Top-1% $\Lambda$ | Top-0.1% $\Lambda$ | Max $\Lambda$ |
| --- | --- | --- | --- | --- |
| CEU | Yes | 100.464 | 220.170 | 992.484 |
| CEU | No | 43.444 | 96.525.547 | 425.335 |
| YRI | Yes | 46.092 | 105.728 | 503.981 |
| YRI | No | 19.339 | 38.000 | 145.126 |

**Table S2:** Gene ontology enrichment analysis of regions with extreme Λ values in the European (CEU) human population.

| GO molecular function | Fold Enrichment | Raw P-value | FDR |
| --- | --- | --- | --- |
| MHC class II receptor activity | > 100 | $4.05 \times 10^{-15}$ | $6.59 \times 10^{-12}$ |
| Immune receptor activity | 40.58 | $6.92 \times 10^{-9}$ | $4.22 \times 10^{-6}$ |
| CD8 receptor binding | > 100 | $1.48 \times 10^{-5}$ | $7.22 \times 10^{-3}$ |
| T cell receptor binding | > 100 | $3.29 \times 10^{-7}$ | $1.78 \times 10^{-4}$ |
| Protein-containing complex binding | 5.96 | $2.72 \times 10^{-5}$ | $1.21 \times 10^{-2}$ |
| MHC class II protein complex binding | > 100 | $1.96 \times 10^{-15}$ | $4.79 \times 10^{-12}$ |
| MHC protein complex binding | > 100 | $1.17 \times 10^{-14}$ | $1.43 \times 10^{-11}$ |
| Beta-2-microglobulin binding | > 100 | $4.43 \times 10^{-5}$ | $1.80 \times 10^{-2}$ |
| Peptide antigen binding | > 100 | $4.37 \times 10^{-17}$ | $2.13 \times 10^{-13}$ |
| Peptide binding | 24.29 | $6.89 \times 10^{-10}$ | $5.60 \times 10^{-7}$ |
| Amide binding | 19.57 | $3.64 \times 10^{-9}$ | $2.54 \times 10^{-6}$ |
| Antigen binding | 41.08 | $1.20 \times 10^{-11}$ | $1.17 \times 10^{-8}$ |

**Table S3:** Gene ontology enrichment analysis of regions with extreme Λ values in the African (YRI) human population.

| GO molecular function | Fold Enrichment | Raw P-value | FDR |
| --- | --- | --- | --- |
| MHC class II receptor activity | > 100 | $6.38 \times 10^{-17}$ | $1.04 \times 10^{-13}$ |
| Immune receptor activity | 25.82 | $9.27 \times 10^{-12}$ | $9.05 \times 10^{-9}$ |
| MHC class II protein complex binding | > 100 | $2.49 \times 10^{-18}$ | $1.22 \times 10^{-14}$ |
| MHC protein complex binding | > 100 | $2.86 \times 10^{-17}$ | $6.99 \times 10^{-14}$ |
| Protein-containing complex binding | 3.70 | $3.52 \times 10^{-5}$ | $1.91 \times 10^{-2}$ |
| Peptide antigen binding | 93.61 | $2.79 \times 10^{-15}$ | $3.40 \times 10^{-12}$ |
| Peptide binding | 10.43 | $2.14 \times 10^{-7}$ | $1.49 \times 10^{-4}$ |
| Amide binding | 8.40 | $1.25 \times 10^{-6}$ | $7.60 \times 10^{-4}$ |
| Antigen binding | 17.64 | $2.71 \times 10^{-9}$ | $2.20 \times 10^{-6}$ |

**Table S4:** Spearman correlations of Λ statistics calculated with different distance metrics from demography-matched neutral whole genome simulations with variable recombination rate (mean across 100 replicates) and from empirical data.

| Population | Data Type | # windows vs. bps | cM vs. bps |
| --- | --- | --- | --- |
| CEU | Simulated | 0.929 | 0.912 |
| YRI | Simulated | 0.967 | 0.895 |
| CEU | Empirical | 0.977 | 0.907 |
| YRI | Empirical | 0.967 | 0.909 |

**Table S5:** Regions of extreme Λ values (unphased analysis) in the CEU population and the genes contained therein. 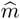 is the inferred number of sweeping haplotypes, and 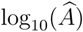 is the estimated sweep width.

| Chr | Start (bp) | Stop (bp) | $\hat{m}$ | $\log_{10}(\hat{A})$ | Max $\Lambda$ | Genes |
| --- | --- | --- | --- | --- | --- | --- |
| 2 | 135,517,503 | 135,699,618 | 2 | 8.686 | 464.366 | *ACMSD*, *MIR5590*, *CCNT2-AS1*, *CCNT2* |
| 2 | 135,699,619 | 135,889,801 | 2 | 8.686 | 469.480 | *CCNT2*, *MAP3K19*, *RAB3GAP1* |
| 2 | 135,936,894 | 136,318,298 | 1 | 8.686 | 677.751 | *ZRANB3*, *R3HDM1* |
| 2 | 136,494,985 | 136,788,904 | 1 | 8.686 | 890.887 | *UBXN4*, *LCT*, *LOC100507600*, *MCM6*, *DARS*, *DARS-AS1* |
| 4 | 34,363,017 | 34,399,713 | 4 | 8.686 | 440.658 | – |
| 4 | 61,210,873 | 61,273,068 | 5 | 8.686 | 448.738 | – |
| 6 | 32,491,945 | 32,694,523 | 7 | 7.817 | 699.735 | *HLA-DRB5*, *HLA-DRB6*, *HLA-DRB1*, *HLA-DQA1*, *HLA-DQB1*, *HLA-DQB1-AS1* |

**Table S6:** Regions of extreme Λ values (unphased analysis) in the YRI population and the genes contained therein. 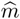 is the inferred number of sweeping haplotypes, and 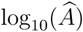 is the estimated sweep width.

| Chr | Start (bp) | Stop (bp) | $\hat{m}$ | $\log_{10}(\hat{A})$ | Max $\Lambda$ | Genes |
| --- | --- | --- | --- | --- | --- | --- |
| 1 | 196,838,929 | 196,882,435 | 9 | 8.252 | 154.62 | *CFHR4* |
| 1 | 223,285,200 | 223,386,011 | 5 | 8.252 | 197.548 | *TLR5* |
| 3 | 27,139,533 | 27,182,014 | 8 | 8.252 | 154.481 | *NEK10* |
| 3 | 46,040,267 | 46,365,291 | 6 | 8.252 | 348.033 | *XCR1, CCR1, CCR3* |
| 3 | 162,438,975 | 162,692,695 | 8 | 8.252 | 296.913 | – |
| 3 | 163,802,717 | 163,857,923 | 8 | 8.252 | 174.977 | – |
| 4 | 12,461,207 | 12,479,264 | 8 | 8.252 | 151.189 | – |
| 4 | 12,503,155 | 12,570,561 | 9 | 8.252 | 168.923 | – |
| 4 | 46,843,639 | 46,884,507 | 9 | 8.252 | 145.785 | *COX7B2* |
| 4 | 74,379,014 | 74,493,240 | 8 | 8.252 | 160.766 | *LOC728040, RASSF6* |
| 6 | 31,171,915 | 31,401,004 | 9 | 7.817 | 352.419 | *HLA-C, HLA-B, MIR6891, MICA* |
| 6 | 32,510,827 | 32,635,225 | 9 | 7.817 | 248.927 | *HLA-DRB6, HLA-DRB1, HLA-DQA1, HLA-DQB1, HLA-DQB1-AS1* |
| 6 | 32,649,806 | 32,694,318 | 10 | 7.817 | 180.873 | – |
| 6 | 32,978,997 | 33,195,684 | 7 | 7.817 | 324.768 | *HLA-DPA1, HLA-DPB1, HLA-DPB2, COL11A2, RXRB, SLC39A7, HSD17B8, MIR219A1, RING1* |
| 6 | 130,541,399 | 130,674,414 | 7 | 7.817 | 244.991 | *SAMD3* |
| 7 | 80,186,004 | 80,428,489 | 6 | 8.252 | 207.636 | *CD36, SEMA3C* |
| 8 | 5,767,335 | 5,824,312 | 10 | 8.252 | 165.733 | – |
| 8 | 9,453,656 | 9,697,983 | 8 | 8.252 | 176.026 | *TNKS* |
| 8 | 50,022,640 | 50,229,742 | 8 | 8.252 | 275.713 | – |
| 9 | 11,774,095 | 11,867,257 | 9 | 8.252 | 161.915 | – |
| 9 | 11,898,626 | 11,940,738 | 7 | 8.252 | 166.824 | – |
| 10 | 102,043,692 | 102,310,074 | 6 | 8.252 | 327.657 | *BLOC1S2, PKD2L1, SCD, OLMALINC, WNT8B, SEC31B, NDUFB8, HIF1AN* |
| 11 | 42,559,011 | 42,612,433 | 8 | 8.252 | 152.469 | – |
| 11 | 42,626,446 | 42,711,242 | 8 | 8.252 | 187.084 | – |
| 12 | 79,543,414 | 79,794,477 | 6 | 8.252 | 334.766 | *SYT1* |
| 12 | 82,748,538 | 82,820,129 | 9 | 8.252 | 173.956 | *CCDC59, METTL25* |
| 12 | 82,859,617 | 82,931,718 | 8 | 8.252 | 151.183 | *METTL25* |
| 13 | 89,192,255 | 89,235,146 | 9 | 8.252 | 157.25 | *LINC00433* |
| 14 | 48,245,871 | 48,321,935 | 8 | 7.817 | 168.862 | *LINC00648* |
| 14 | 48,355,469 | 48,392,903 | 7 | 7.817 | 151.657 | – |
| 14 | 48,656,257 | 48,694,596 | 9 | 7.817 | 156.422 | – |
| 14 | 106,434,952 | 106,471,410 | 5 | 7.817 | 155.709 | *ADAM6* |
| 15 | 55,111,308 | 55,359,448 | 7 | 8.252 | 302.699 | – |
| 17 | 3,530,561 | 3,658,973 | 6 | 8.252 | 294.719 | *SHPK, CTNS, TAX1BP3, P2RX5-TAX1BP3, EMC6, P2RX5, ITGAE, GSG2* |
| 17 | 60,593,430 | 60,624,708 | 6 | 8.252 | 152.548 | *TLK2* |
| 19 | 38,860,718 | 38,951,198 | 8 | 8.252 | 182.638 | *CATSPERG, PSMD8, GGN, SPRED3, FAM98C, RASGRP4, RYR1* |
| 20 | 37,313,016 | 37,490,473 | 7 | 8.686 | 159.738 | *SLC32A1, ACTR5, PPP1R16B* |
| 22 | 36,571,575 | 36,749,069 | 7 | 8.252 | 184.972 | *APOL4, APOL2, APOL1, MYH9, MIR6819* |

**Table S7:** Regions of extreme Λ values in the New York City rat population that contain annotated genes in genome build RN5. 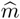 is the inferred number of sweeping haplotypes, and 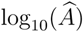 is the estimated sweep width.

| Chr | Start (bp) | Stop (bp) | $\hat{m}$ | $\log_{10}(\hat{A})$ | Max $\Lambda$ | Genes |
| --- | --- | --- | --- | --- | --- | --- |
| 1 | 156,312,419 | 157,034,325 | 3 | 7.765 | 673.695 | *Olr23*, *Olr24*, *Olr25*, *Olr27*,<br>*Olr29*, *Olr30*, *Olr32*, *Olr34*,<br>*Folh1* |
| 1 | 230,123,826 | 230,327,101 | 3 | 7.765 | 177.137 | *Slc22a24* |
| 2 | 247,180,473 | 247,304,944 | 1 | 7.727 | 113.896 | *Ndst3* |
| 2 | 269,229,545 | 269,542,892 | 1 | 7.727 | 675.289 | *Sh3glb1*, *Clca2*, *Clca4l*, *Clca4*,<br>*Clca1*, *Clca5*, *Odf2l* |
| 3 | 10,383,050 | 10,578,622 | 3 | 7.766 | 168.801 | *Abo3* |
| 4 | 41,336,974 | 41,611,989 | 1 | 7.910 | 168.115 | *Foxp2* |
| 4 | 133,838,379 | 133,973,032 | 3 | 7.910 | 107.921 | *Moxd2*, *Prss58*, *Tryx5* |
| 4 | 211,484,812 | 211,533,858 | 4 | 7.910 | 93.3354 | *Clec2e* |
| 4 | 216,573,081 | 216,659,618 | 2 | 7.910 | 106.382 | *Cacna1c* |
| 5 | 107,548,396 | 107,587,543 | 1 | 8.226 | 92.9657 | *Sh3gl2* |
| 6 | 92,808,416 | 92,850,870 | 3 | 7.837 | 89.5456 | *Lrfn5* |
| 7 | 17,187,521 | 17,434,428 | 1 | 7.751 | 147.23 | *Vom2r55* |
| 7 | 27,437,241 | 27,591,701 | 2 | 7.751 | 130.023 | *Nt5dc3*, *Stab2* |
| 7 | 29,959,185 | 30,172,142 | 1 | 7.751 | 126.808 | *Ano4*, *Nr1h4* |
| 7 | 45,385,763 | 45,446,200 | 2 | 7.751 | 93.5232 | *Slc6a15* |
| 8 | 119,571,436 | 119,673,192 | 1 | 8.378 | 124.087 | *Arpp21* |
| 9 | 55,257,470 | 56,770,741 | 1 | 8.134 | 301.09 | *Tmeff2* |
| 13 | 1,901,678 | 2,824,028 | 2 | 7.949 | 410.203 | *Dsel* |
| 13 | 12,195,969 | 12,424,761 | 2 | 7.949 | 199.78 | *Cntnap5c* |
| 13 | 68,568,273 | 68,885,107 | 3 | 7.949 | 508.191 | *Brinp3* |
| 14 | 32,656,671 | 32,892,843 | 1 | 7.740 | 204.445 | *Igfbp7*, *Polr2b* |
| 17 | 80,051,550 | 80,219,722 | 2 | 7.860 | 103.731 | *Fam107b* |

**Table S8:** Regions of extreme Λ values in the New York City rat population that do not contain annotated genes in genome build RN5. 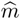 is the inferred number of sweeping haplotypes, and 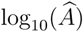 is the estimated sweep width.

| Chr | Start (bp) | Stop (bp) | $\hat{m}$ | $\log_{10}(\hat{A})$ | Max $\Lambda$ | Genes |
| --- | --- | --- | --- | --- | --- | --- |
| 2 | 96,271,808 | 96,560,663 | 2 | 7.727 | 410.932 | — |
| 2 | 96,881,372 | 97,074,854 | 3 | 7.727 | 326.032 | — |
| 2 | 170,378,708 | 170,519,263 | 2 | 7.727 | 95.1473 | — |
| 2 | 272,083,254 | 272,359,881 | 1 | 7.727 | 241.38 | — |
| 3 | 23,325,877 | 23,538,235 | 2 | 7.766 | 213.939 | — |
| 3 | 143,427,703 | 143,448,409 | 1 | 7.766 | 90.6533 | — |
| 4 | 40,399,215 | 40,462,778 | 1 | 7.910 | 97.1179 | — |
| 4 | 47,317,126 | 47,413,412 | 3 | 7.910 | 110.665 | — |
| 4 | 47,479,582 | 47,521,098 | 3 | 7.910 | 91.0695 | — |
| 5 | 23,200,838 | 23,418,591 | 3 | 8.226 | 164.644 | — |
| 5 | 136,100,860 | 136,251,275 | 2 | 8.226 | 103.145 | — |
| 5 | 136,259,254 | 136,263,345 | 2 | 8.226 | 92.3248 | — |
| 5 | 136,271,400 | 136,321,452 | 2 | 8.226 | 104.762 | — |
| 5 | 136,904,389 | 136,961,220 | 1 | 8.226 | 99.4727 | — |
| 5 | 137,107,962 | 137,167,762 | 1 | 8.226 | 91.6863 | — |
| 6 | 93,246,089 | 93,895,837 | 2 | 7.837 | 285.452 | — |
| 6 | 128,257,859 | 128,342,359 | 1 | 7.837 | 101.541 | — |
| 6 | 151,455,228 | 151,704,771 | 1 | 7.837 | 394.659 | — |
| 6 | 151,877,456 | 151,898,077 | 3 | 7.837 | 94.741 | — |
| 7 | 19,461,789 | 19,650,647 | 3 | 7.751 | 133.615 | — |
| 7 | 43,024,984 | 43,049,245 | 2 | 7.751 | 89.1577 | — |
| 7 | 43,095,862 | 43,141,691 | 2 | 7.751 | 106.754 | — |
| 7 | 89,090,760 | 89,282,665 | 2 | 7.751 | 142.529 | — |
| 7 | 109,432,422 | 109,528,692 | 1 | 7.751 | 91.6154 | — |
| 7 | 127,786,183 | 127,890,186 | 1 | 7.751 | 134.217 | — |
| 7 | 137,032,536 | 137,122,391 | 1 | 7.751 | 120.846 | — |
| 8 | 3,092,342 | 3,236,525 | 3 | 8.378 | 113.975 | — |
| 9 | 56,797,610 | 56,900,761 | 2 | 8.134 | 125.534 | — |
| 9 | 57,116,060 | 57,176,933 | 1 | 8.134 | 93.6 | — |
| 9 | 119,697,786 | 119,952,597 | 1 | 8.134 | 192.989 | — |
| 10 | 28,182,853 | 28,209,022 | 2 | 8.240 | 88.2872 | — |
| 13 | 3,073,604 | 3,247,275 | 3 | 7.949 | 185.906 | — |
| 13 | 13,376,314 | 13,603,477 | 1 | 7.949 | 185.505 | — |
| 13 | 14,872,399 | 14,887,445 | 3 | 7.949 | 88.3306 | — |
| 13 | 43,647,576 | 43,944,224 | 4 | 7.949 | 235.378 | — |
| 14 | 20,095,568 | 20,119,479 | 2 | 7.740 | 88.9939 | — |
| 14 | 30,706,707 | 30,902,046 | 1 | 7.740 | 166.608 | — |
| 14 | 30,925,938 | 30,979,611 | 1 | 7.740 | 95.0194 | — |
| 14 | 30,990,277 | 31,067,039 | 2 | 7.740 | 95.3088 | — |
| 14 | 35,268,129 | 35,338,455 | 1 | 7.740 | 91.7566 | — |
| 14 | 67,880,756 | 68,116,102 | 1 | 7.740 | 169.303 | — |
| 14 | 95,349,113 | 95,473,128 | 2 | 4.103 | 135.307 | — |
| 15 | 57,136,691 | 57,510,633 | 1 | 7.968 | 155.564 | — |
| 17 | 79,592,898 | 79,717,397 | 1 | 7.860 | 116.18 | — |
| 18 | 61,104,703 | 61,227,007 | 1 | 8.141 | 123.9 | — |
| 18 | 61,609,058 | 61,788,832 | 1 | 8.141 | 138.145 | — |
| 20 | 54,099,523 | 54,274,713 | 3 | 7.885 | 118.206 | — |

**Figure S1:**
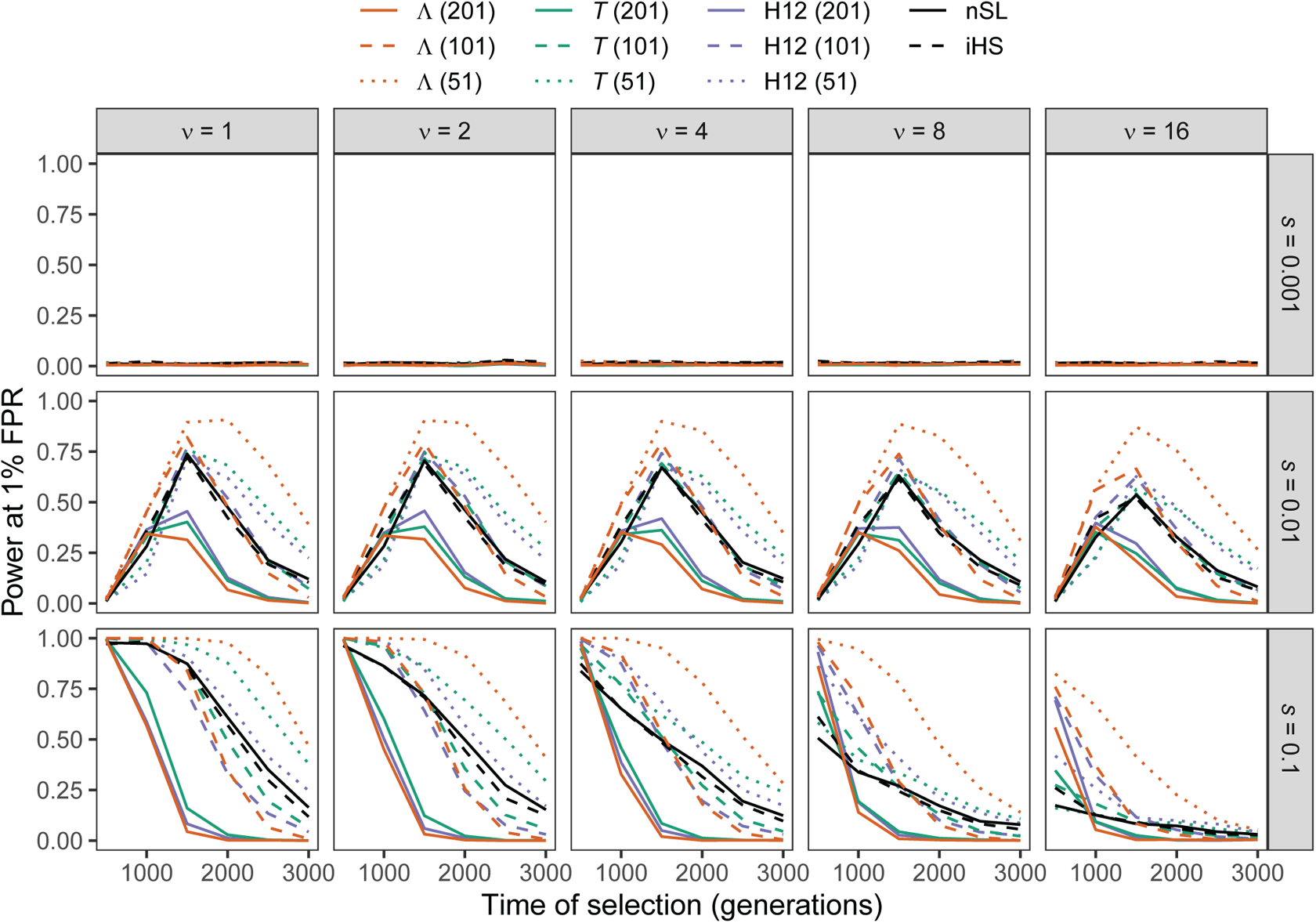
Power at a 1% false positive rate (FPR) as a function of selection start time for applications of Λ, *T*, and H12 with windows of size 51, 101, and 201 SNPs, as well *nS_L_* and iHS under simulations of a constant-size demographic history for per-generation selection coefficients of *s* ∈ {0.001, 0.01, 0.1} on the rows. Classification ability demonstrated for selection start times of *t* ∈ {500, 1000, 1500, 2000, 2500, 3000} generations prior to sampling for *ν* ∈ {1, 2, 4, 8, 16} initially-selected haplotypes (columns). Results are based on a sample of *n* = 50 diploid individuals and the haplotype frequency spectra for the Λ and *T* statistics truncated at *K* = 10 haplotypes.

**Figure S2:**
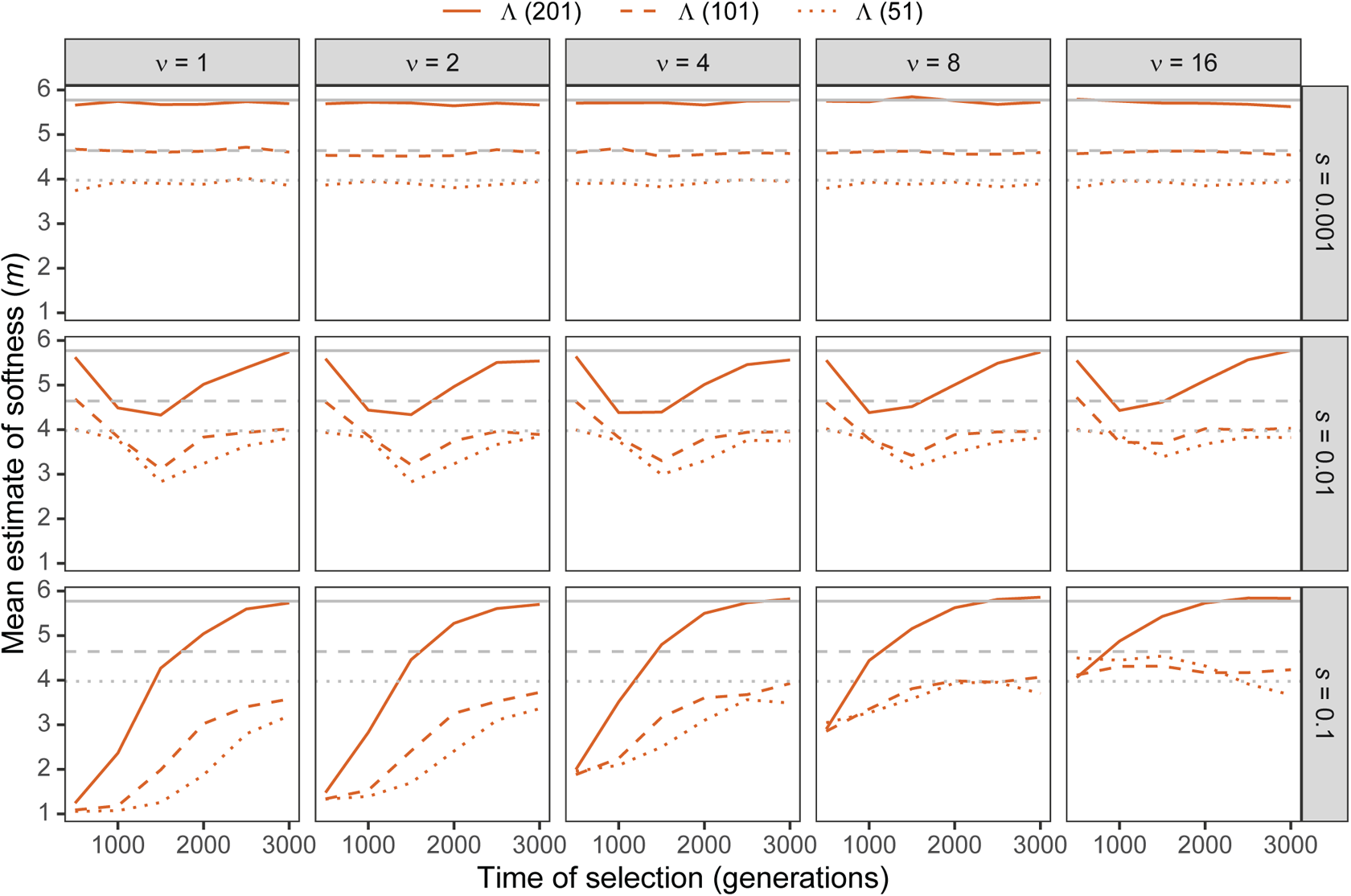
Estimated sweep softness illustrated by mean estimated number of sweeping haplotypes (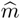) in Λ with windows of size 51, 101, and 201 SNPs under simulations of a constant-size demographic history for per-generation selection coefficients of *s* ∈ {0.001, 0.01, 0.1} on the rows. Mean estimated softness demonstrated for selection start times of t ∈ {500, 1000, 1500, 2000, 2500, 3000} generations prior to sampling for ν ∈ {1, 2, 4, 8, 16} initially-selected haplotypes (columns). Gray solid, dashed, and dotted horizontal lines are the corresponding mean 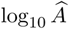 values for Λ applied to neutral simulations. Results are based on a sample of n = 50 diploid individuals and the haplotype frequency spectrum for the Λ statistic truncated at K = 10 haplotypes.

**Figure S3:**
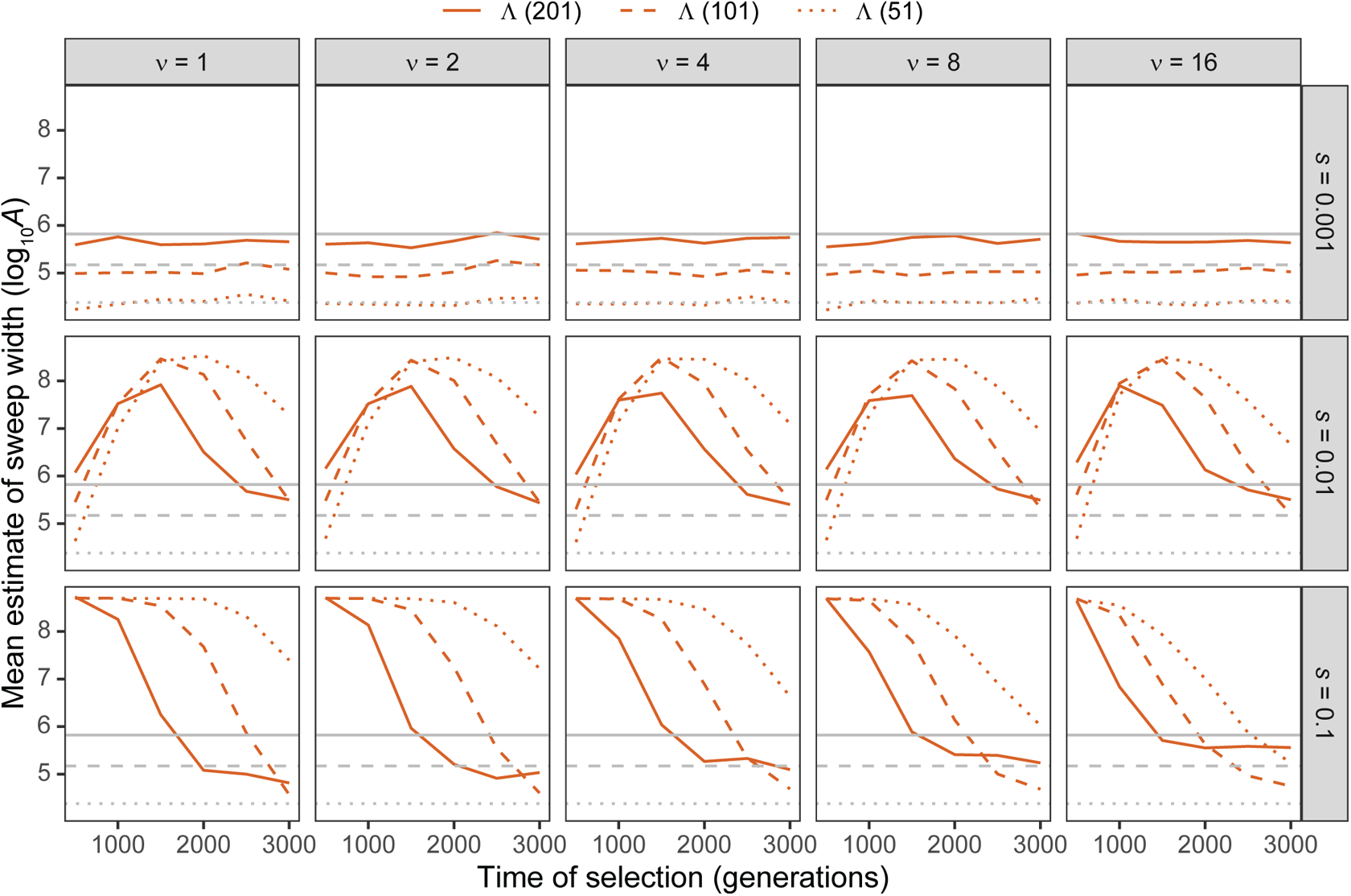
Estimated sweep width illustrated by mean estimated genomic size influenced by the sweep (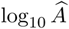) in Λ with windows of size 51, 101, and 201 SNPs under simulations of a constant-size demographic history for per-generation selection coefficients of *s* ∈ {0.001, 0.01, 0.1} on the rows. Mean estimated genomic size influenced by sweeps demonstrated for selection start times of *t* ∈ {500, 1000, 1500, 2000, 2500, 3000} generations prior to sampling for *ν* ∈ {1, 2, 4, 8, 16} initially-selected haplotypes (columns). Gray solid, dashed, and dotted horizontal lines are the corresponding mean 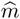 values for Λ applied to neutral simulations. Results are based on a sample of *n* = 50 diploid individuals and the haplotype frequency spectrum for the Λ statistic truncated at *K* = 10 haplotypes.

**Figure S4:**
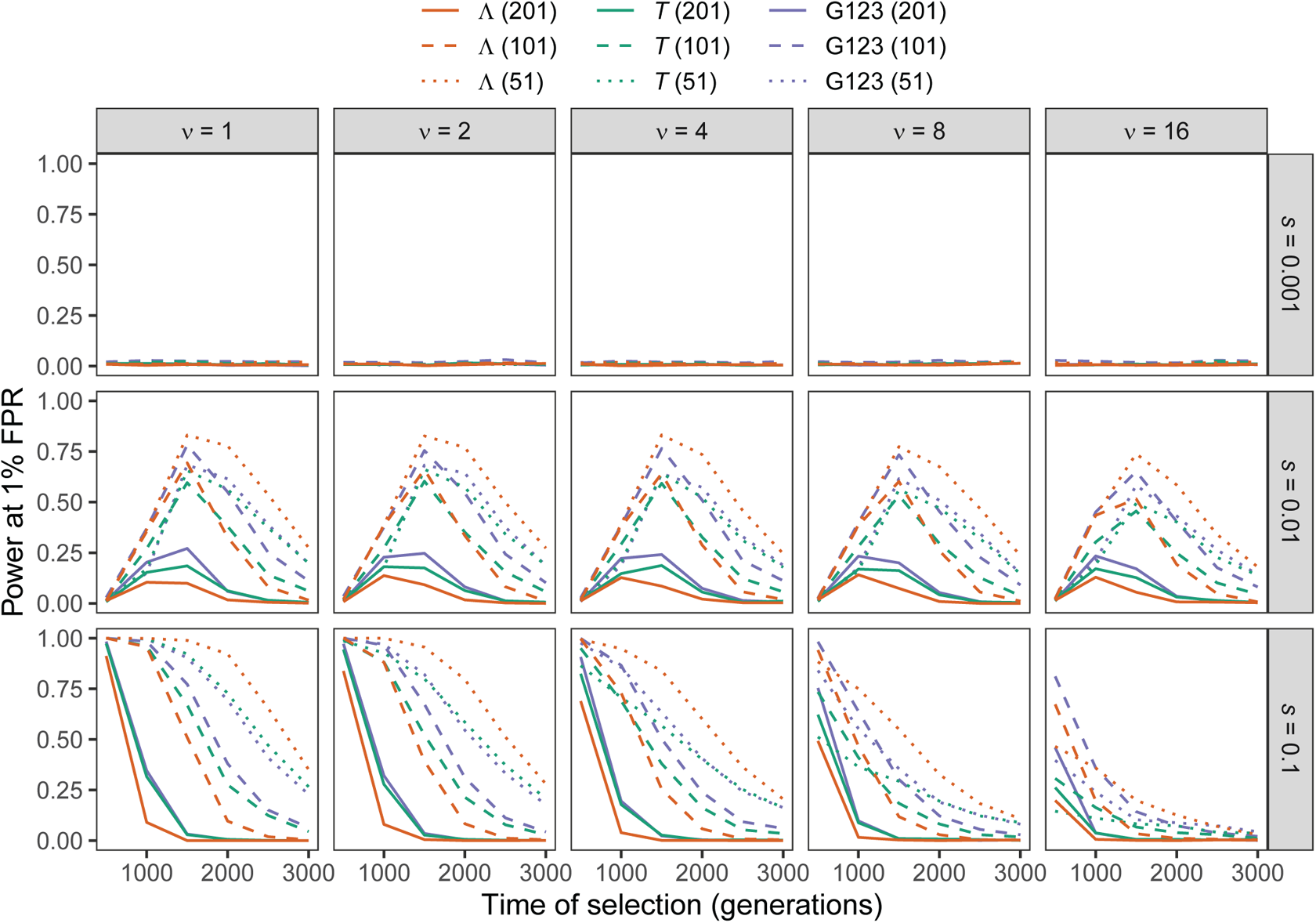
Power at a 1% false positive rate (FPR) as a function of selection start time for applications of Λ, *T*, and G123 with windows of size 51, 101, and 201 SNPs to unphased multilocus genotype input data under simulations of a constant-size demographic history for per-generation selection coefficients of *s* ∈ {0.001, 0.01, 0.1} on the rows. Classification ability demonstrated for selection start times of *t* ∈ {500, 1000, 1500, 2000, 2500, 3000} generations prior to sampling for *ν* ∈ {1, 2, 4, 8, 16} initially-selected haplotypes (columns). Results are based on a sample of *n* = 50 diploid individuals and the multilocus genotype frequency spectra for the Λ and *T* statistics truncated at *K* = 10 multilocus genotypes.

**Figure S5:**
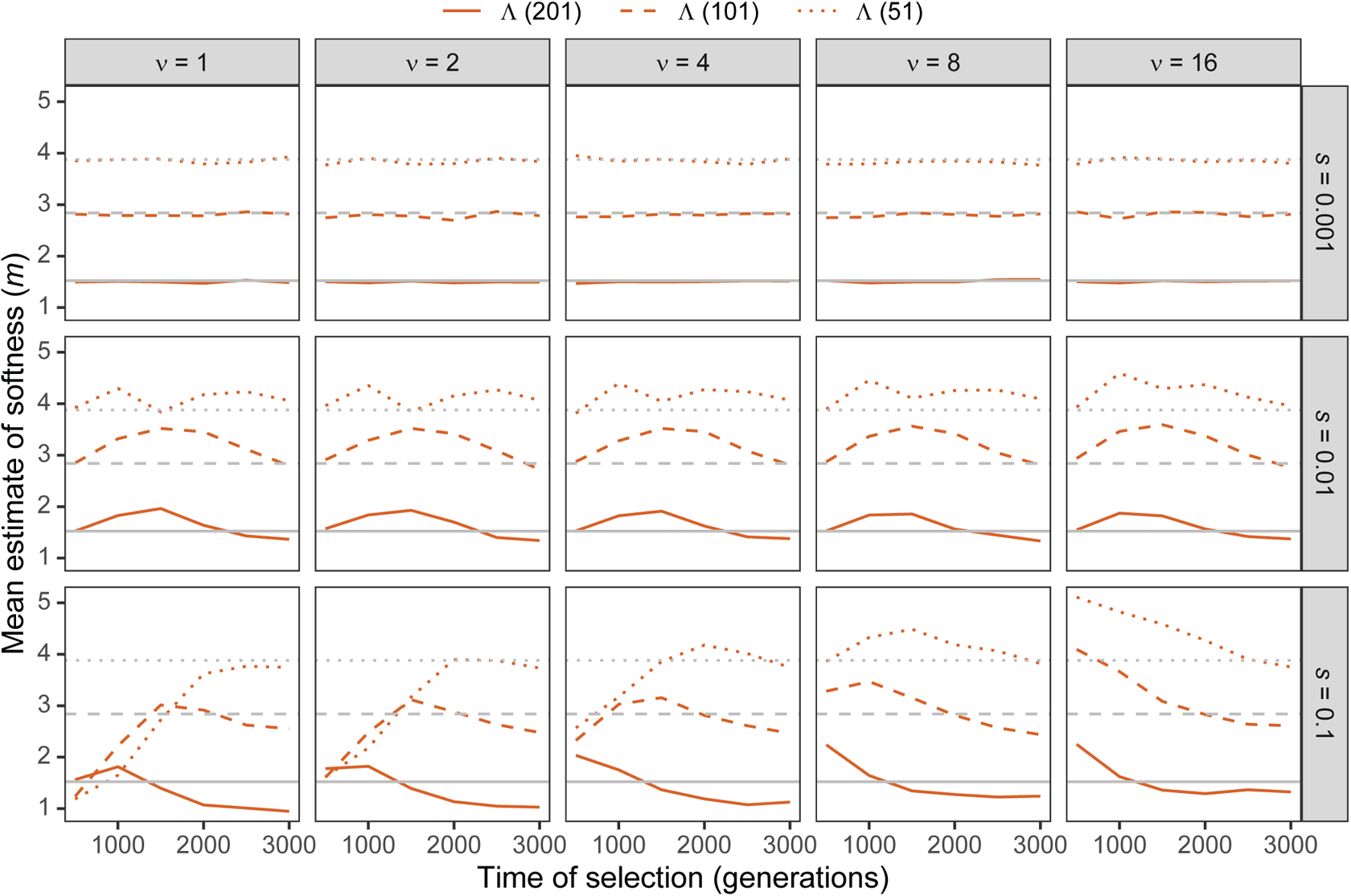
Estimated sweep softness illustrated by mean estimated number of sweeping haplotypes (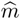) in Λ with windows of size 51, 101, and 201 SNPs applied to unphased multilocus input data under simulations of a constant-size demographic history for per-generation selection coefficients of *s* ∈ {0.001, 0.01, 0.1} on the rows. Mean estimated softness demonstrated for selection start times of *t* ∈ {500, 1000, 1500, 2000, 2500, 3000} generations prior to sampling for *ν* ∈ {1, 2, 4, 8, 16} initially-selected haplotypes (columns). Gray solid, dashed, and dotted horizontal lines are the corresponding mean 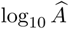 values for Λ applied to neutral simulations. Results are based on a sample of n = 50 diploid individuals and the multilocus genotype frequency spectrum for the Λ statistic truncated at K = 10 multilocus genotypes.

**Figure S6:**
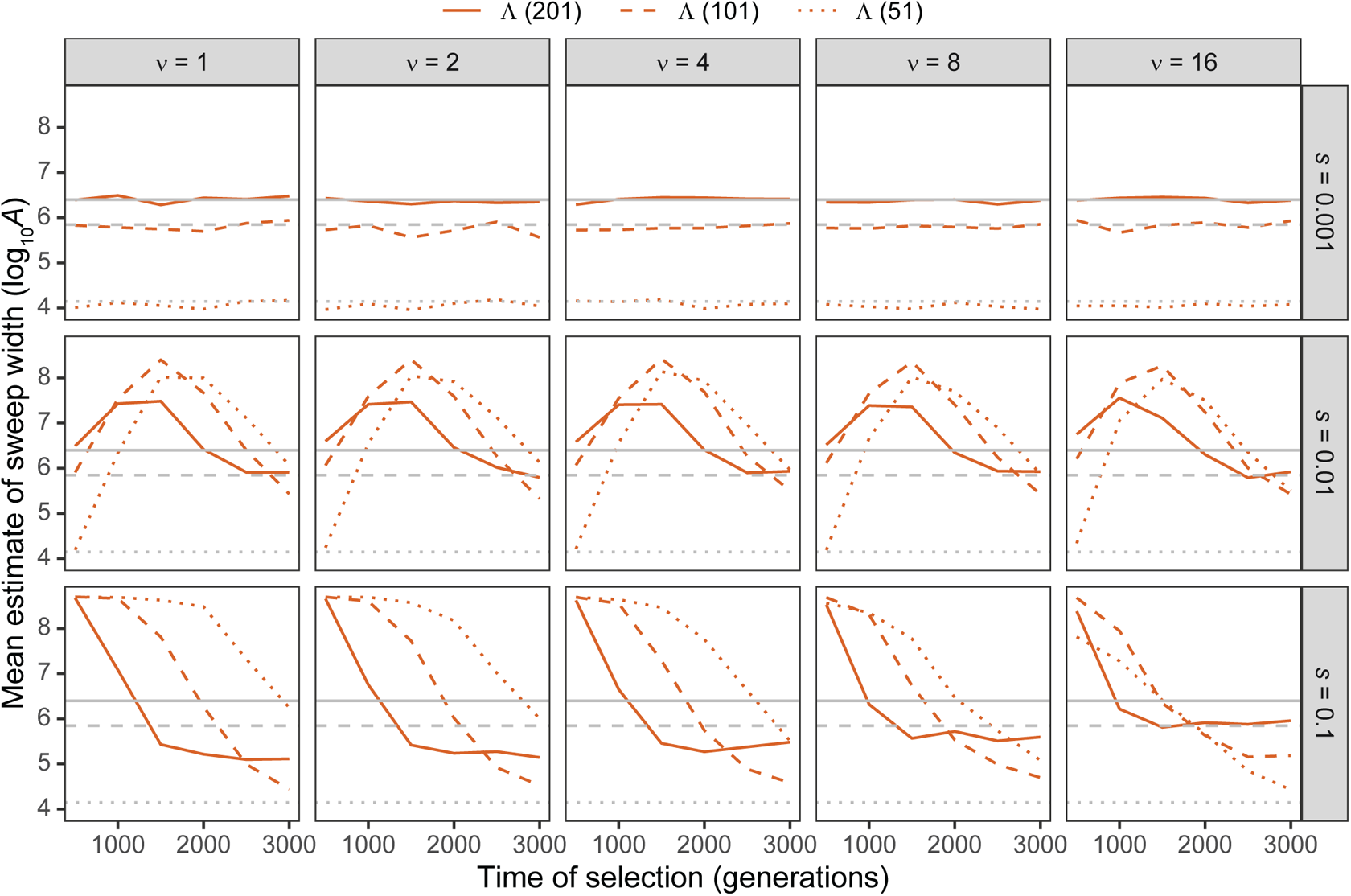
Estimated sweep width illustrated by mean estimated genomic size influenced by the sweep (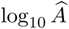) in Λ with windows of size 51, 101, and 201 SNPs applied to unphased multilocus input data under simulations of a constant-size demographic history for per-generation selection coefficients of *s ∈* {0.001, 0.01, 0.1} on the rows. Mean estimated genomic size influenced by sweeps demonstrated for selection start times of *t ∈* {500, 1000, 1500, 2000, 2500, 3000} generations prior to sampling for *ν ∈* {1, 2, 4, 8, 16} initially-selected haplotypes (columns). Gray solid, dashed, and dotted horizontal lines are the corresponding mean 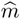 values for Λ applied to neutral simulations. Results are based on a sample of *n* = 50 diploid individuals and the multilocus genotype frequency spectrum for the Λ statistic truncated at *K* = 10 multilocus genotypes.

**Figure S7:**
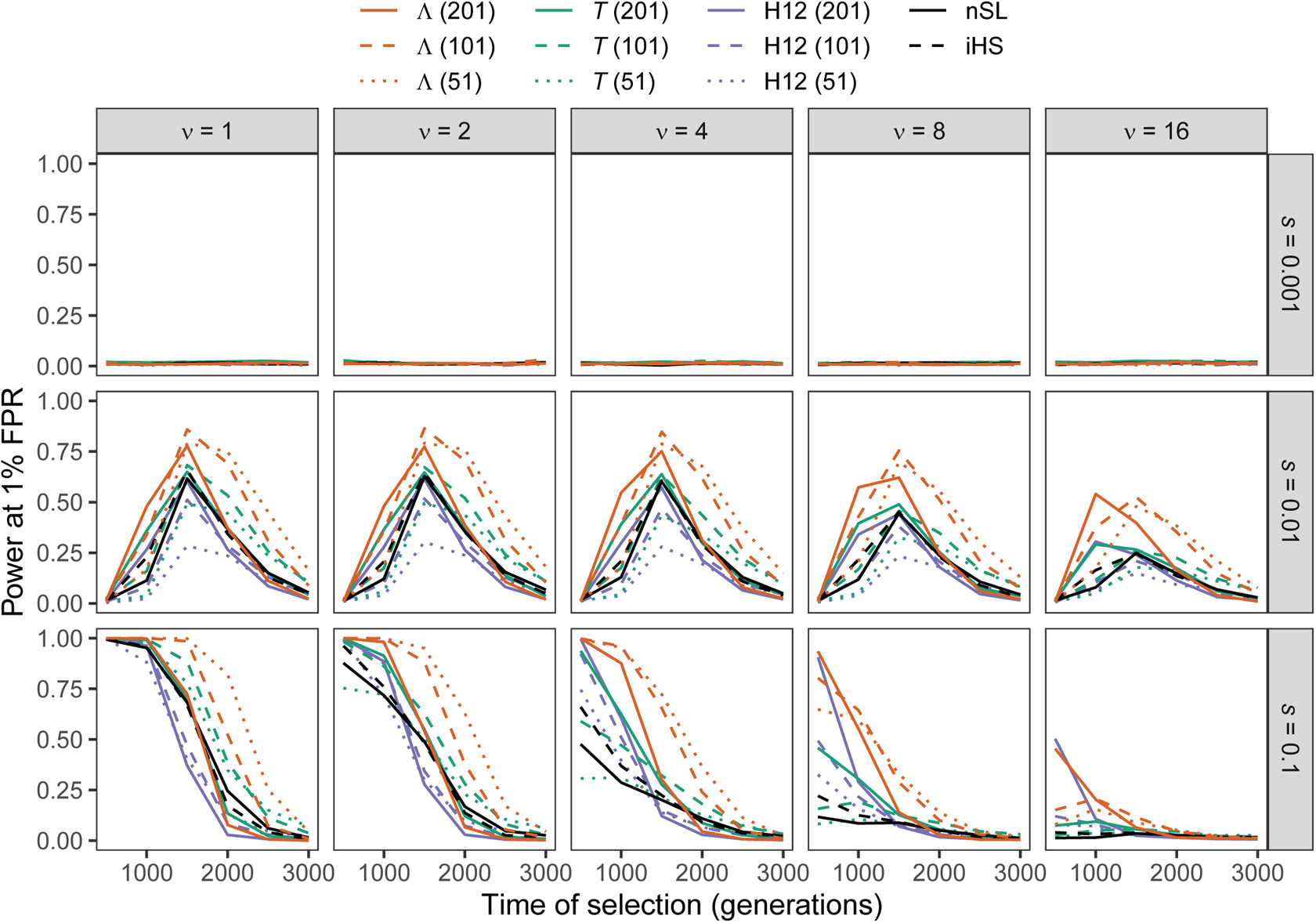
Power at a 1% false positive rate (FPR) as a function of selection start time for applications of Λ, *T*, and H12 with windows of size 51, 101, and 201 SNPs, as well *nS_L_* and iHS under simulations of the human central European (CEU) demographic history of Terhorst et al. [2017] for per-generation selection coefficients of *s ∈* {0.001, 0.01, 0.1} on the rows. Classification ability demonstrated for selection start times of *t ∈* {500, 1000, 1500, 2000, 2500, 3000} generations prior to sampling for *ν ∈* {1, 2, 4, 8, 16} initially-selected haplotypes (columns). Results are based on a sample of *n* = 50 diploid individuals and the haplotype frequency spectra for the Λ and *T* statistics truncated at *K* = 10 haplotypes.

**Figure S8:**
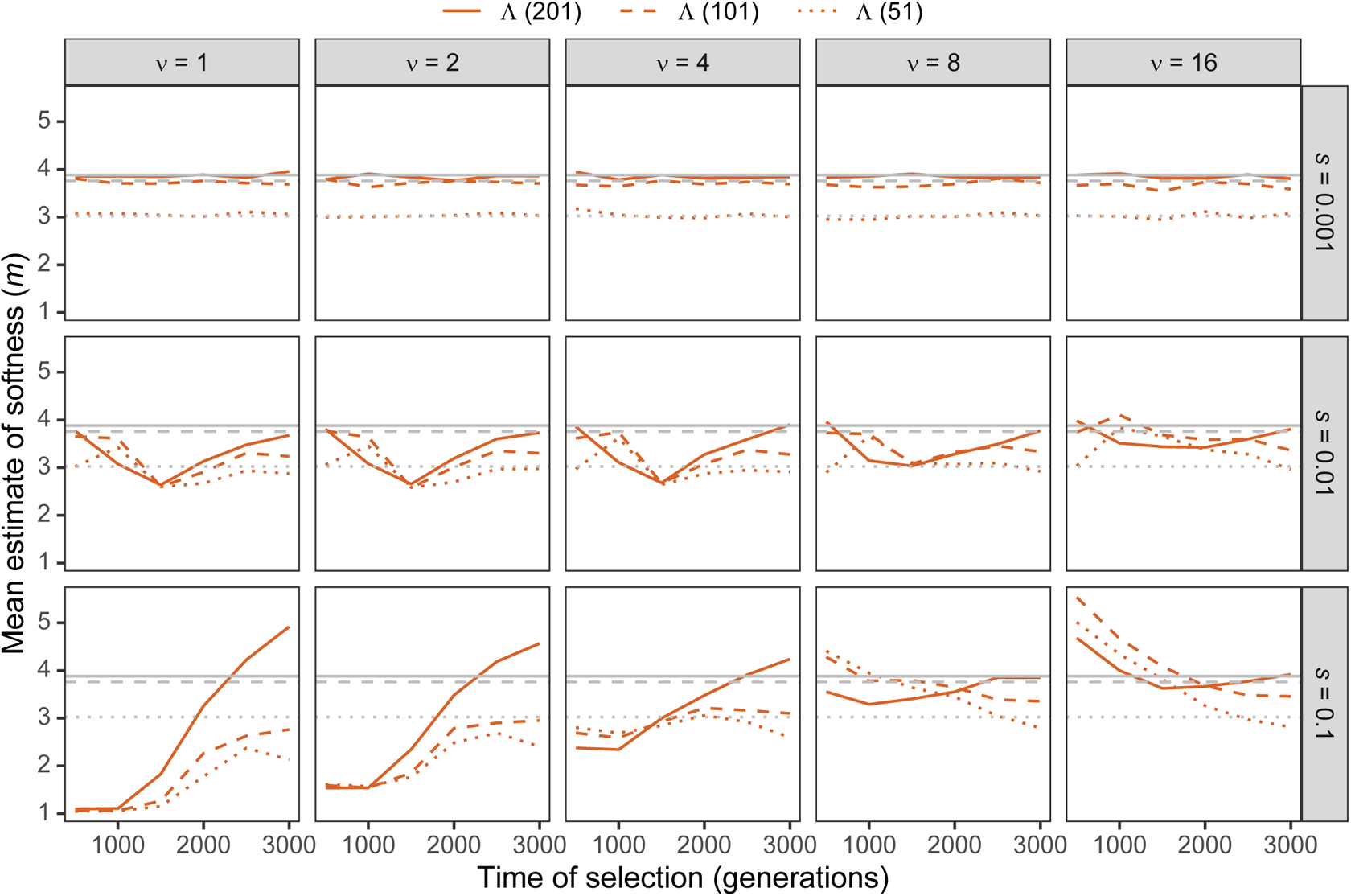
Estimated sweep softness illustrated by mean estimated number of sweeping haplotypes (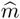) in Λ with windows of size 51, 101, and 201 SNPs under simulations of the human central European (CEU) demographic history of Terhorst et al. [2017] for per-generation selection coefficients of *s ∈* {0.001, 0.01, 0.1} on the rows. Mean estimated softness demonstrated for selection start times of *t ∈* {500, 1000, 1500, 2000, 2500, 3000} generations prior to sampling for *ν ∈* {1, 2, 4, 8, 16} initially-selected haplotypes (columns). Gray solid, dashed, and dotted horizontal lines are the corresponding mean 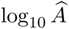 values for Λ applied to neutral simulations. Results are based on a sample of *n* = 50 diploid individuals and the haplotype frequency spectrum for the Λ statistic truncated at *K* = 10 haplotypes.

**Figure S9:**
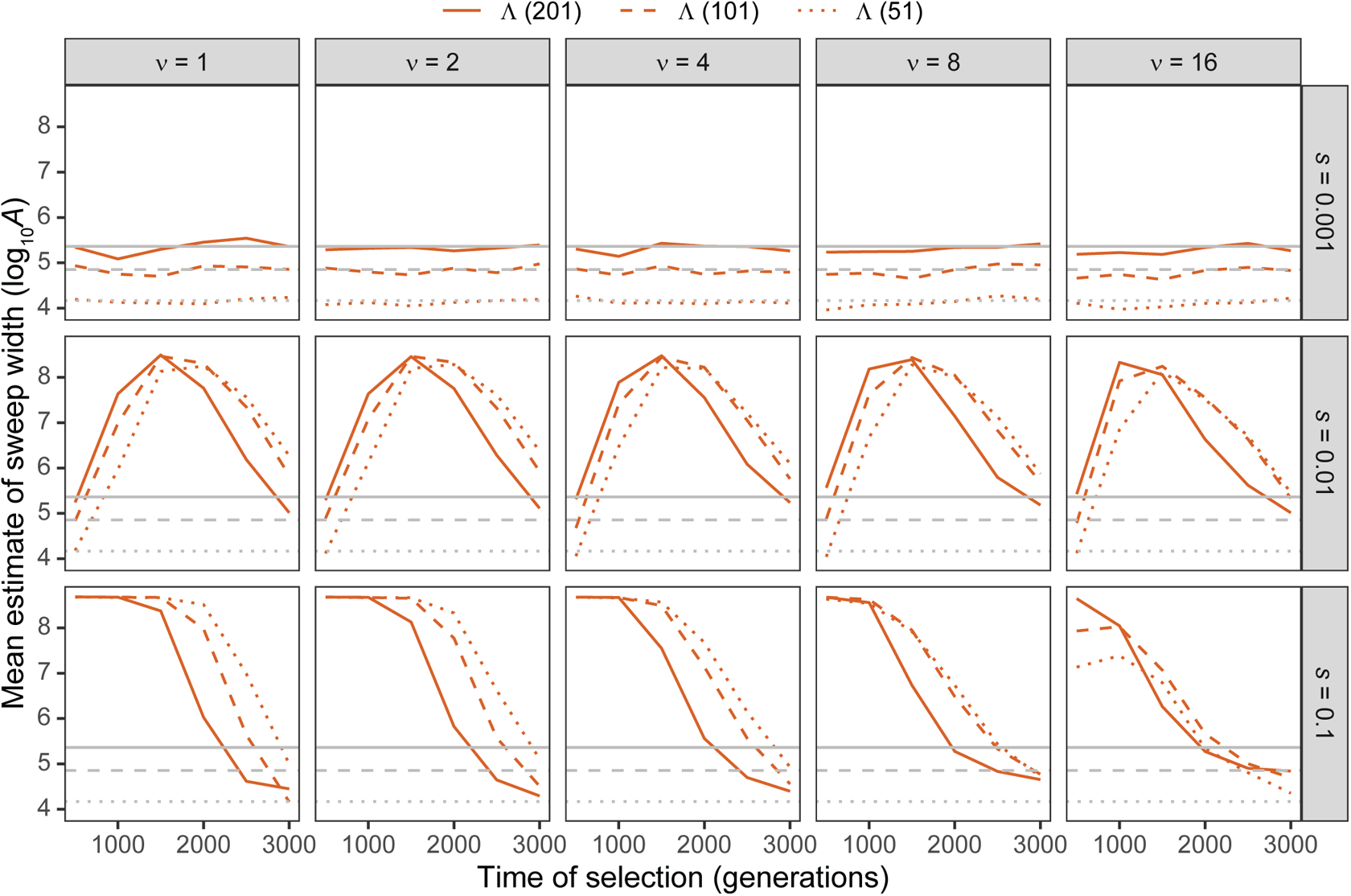
Estimated sweep width illustrated by mean estimated genomic size influenced by the sweep (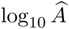) in Λ with windows of size 51, 101, and 201 SNPs under simulations of the human central European (CEU) demographic history of Terhorst et al. [2017] for per-generation selection coefficients of *s ∈* {0.001, 0.01, 0.1} on the rows. Mean estimated genomic size influenced by sweeps demonstrated for selection start times of *t ∈* {500, 1000, 1500, 2000, 2500, 3000} generations prior to sampling for *ν ∈* {1, 2, 4, 8, 16} initially-selected haplotypes (columns). Gray solid, dashed, and dotted horizontal lines are the corresponding mean 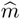 values for Λ applied to neutral simulations. Results are based on a sample of *n* = 50 diploid individuals and the haplotype frequency spectrum for the Λ statistic truncated at *K* = 10 haplotypes.

**Figure S10:**
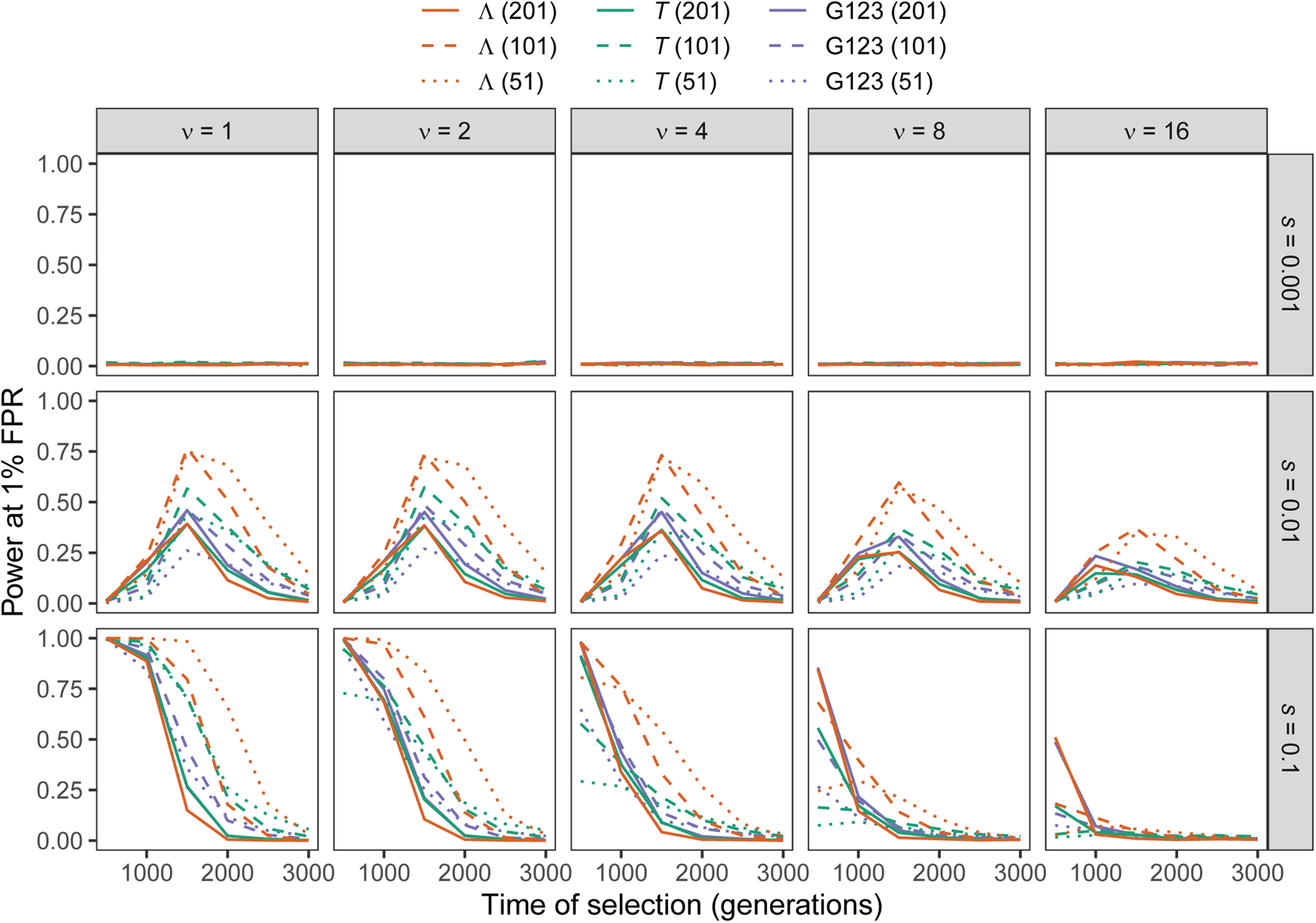
Power at a 1% false positive rate (FPR) as a function of selection start time for applications of Λ, *T*, and G123 with windows of size 51, 101, and 201 SNPs to unphased multilocus genotype input data under simulations of the human central European (CEU) demographic history of Terhorst et al. [2017] for per-generation selection coefficients of *s ∈* {0.001, 0.01, 0.1} on the rows. Classification ability demonstrated for selection start times of *t ∈* {500, 1000, 1500, 2000, 2500, 3000} generations prior to sampling for *ν ∈* {1, 2, 4, 8, 16} initially-selected haplotypes (columns). Results are based on a sample of *n* = 50 diploid individuals and the multilocus genotype frequency spectra for the Λ and *T* statistics truncated at *K* = 10 multilocus genotypes.

**Figure S11:**
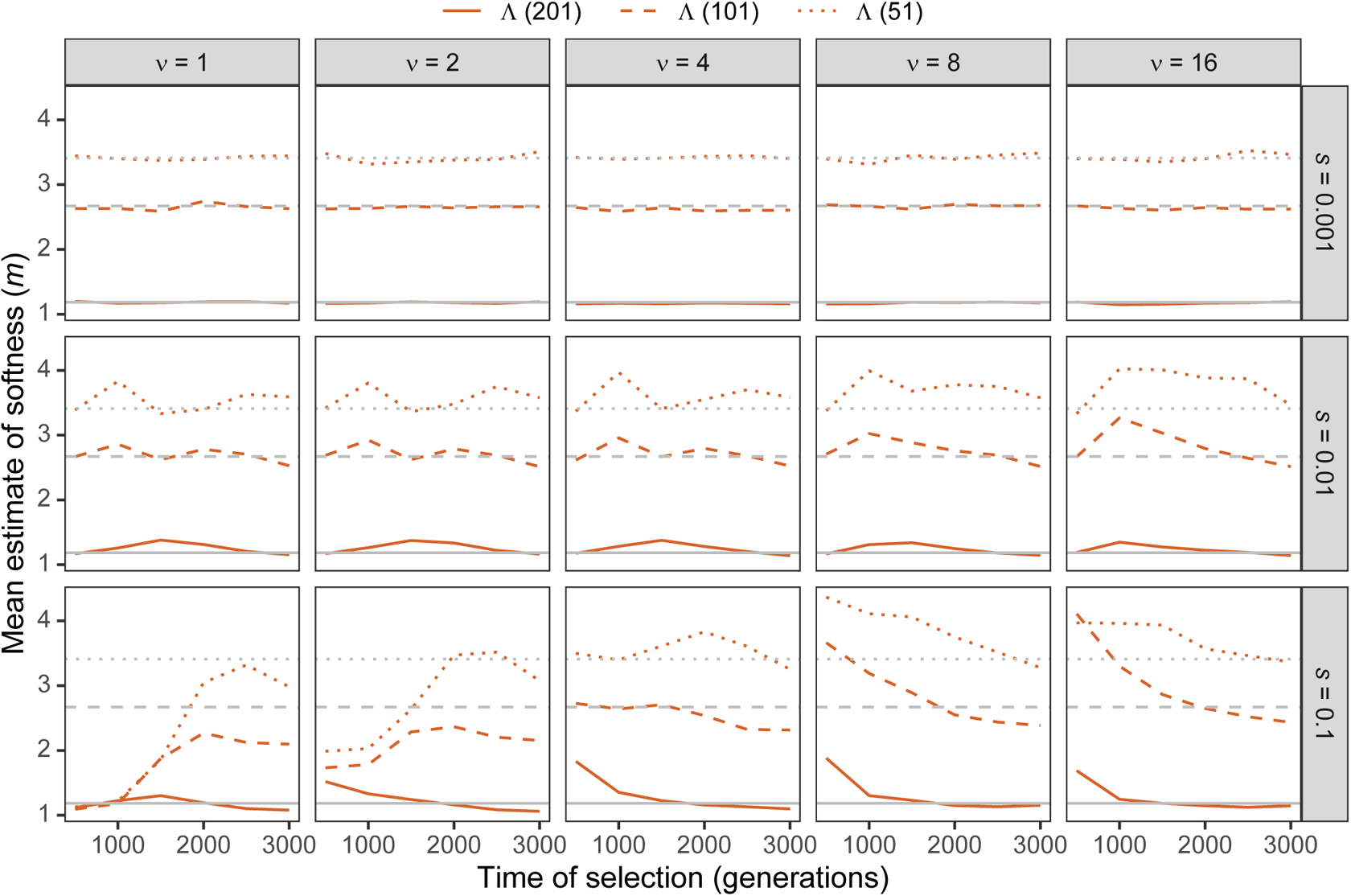
Estimated sweep softness illustrated by mean estimated number of sweeping haplotypes (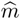) in Λ with windows of size 51, 101, and 201 SNPs applied to unphased multilocus input data under simulations of the human central European (CEU) demographic history of Terhorst et al. [2017] for per-generation selection coefficients of *s ∈* {0.001, 0.01, 0.1} on the rows. Mean estimated softness demonstrated for selection start times of *t ∈* {500, 1000, 1500, 2000, 2500, 3000} generations prior to sampling for *ν ∈* {1, 2, 4, 8, 16} initially-selected haplotypes (columns). Gray solid, dashed, and dotted horizontal lines are the corresponding mean 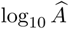 values for Λ applied to neutral simulations. Results are based on a sample of *n* = 50 diploid individuals and the multilocus genotype frequency spectrum for the Λ statistic truncated at *K* = 10 multilocus genotypes.

**Figure S12:**
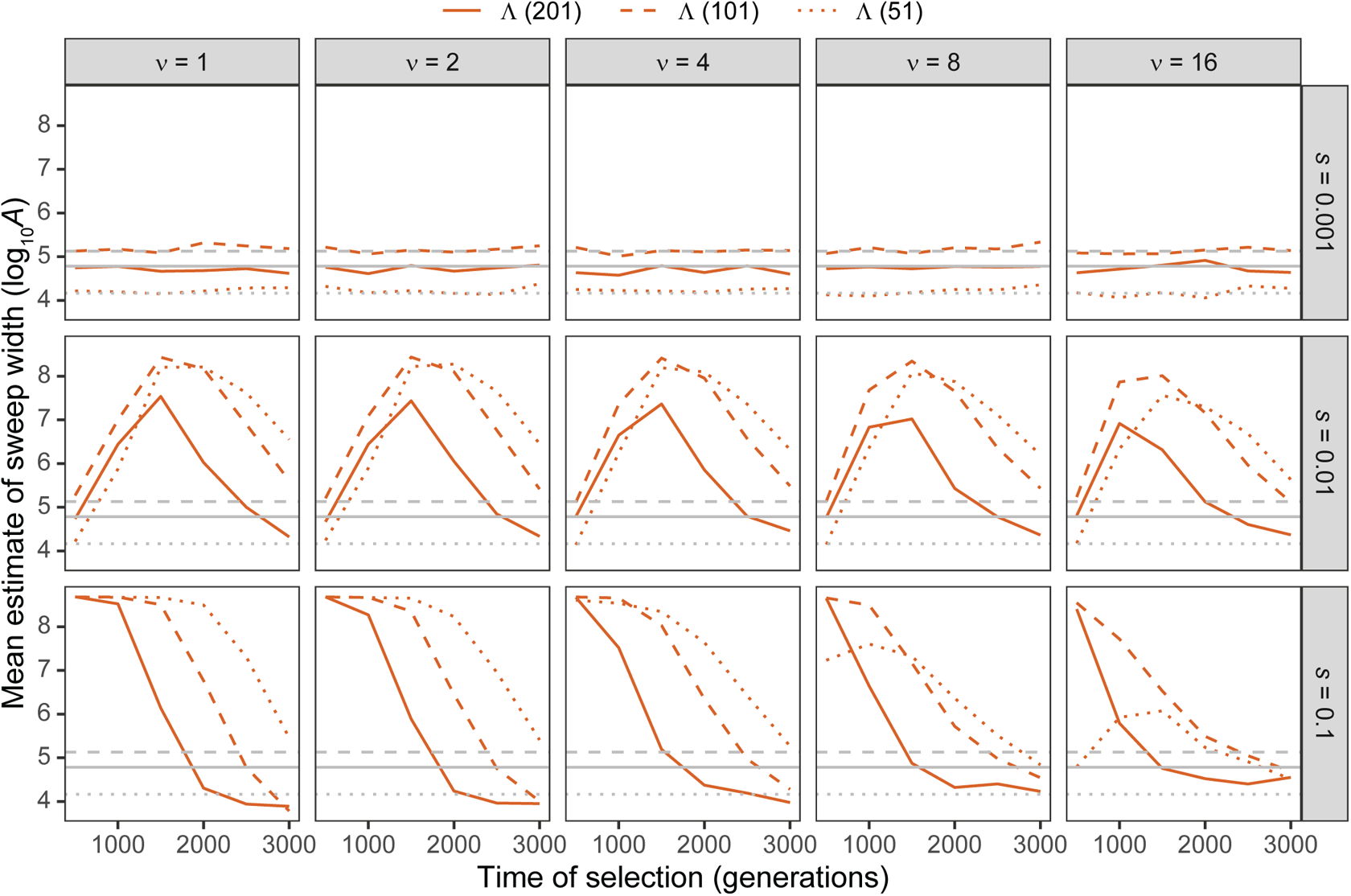
Estimated sweep width illustrated by mean estimated genomic size influenced by the sweep (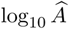) in Λ with windows of size 51, 101, and 201 SNPs applied to unphased multilocus input data under simulations of the human central European (CEU) demographic history of Terhorst et al. [2017] for per-generation selection coefficients of *s ∈* {0.001, 0.01, 0.1} on the rows. Mean estimated genomic size influenced by sweeps demonstrated for selection start times of *t ∈* {500, 1000, 1500, 2000, 2500, 3000} generations prior to sampling for *ν ∈* {1, 2, 4, 8, 16} initially-selected haplotypes (columns). Gray solid, dashed, and dotted horizontal lines are the corresponding mean 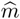 values for Λ applied to neutral simulations. Results are based on a sample of *n* = 50 diploid individuals and the multilocus genotype frequency spectrum for the Λ statistic truncated at *K* = 10 multilocus genotypes.

**Figure S13:**
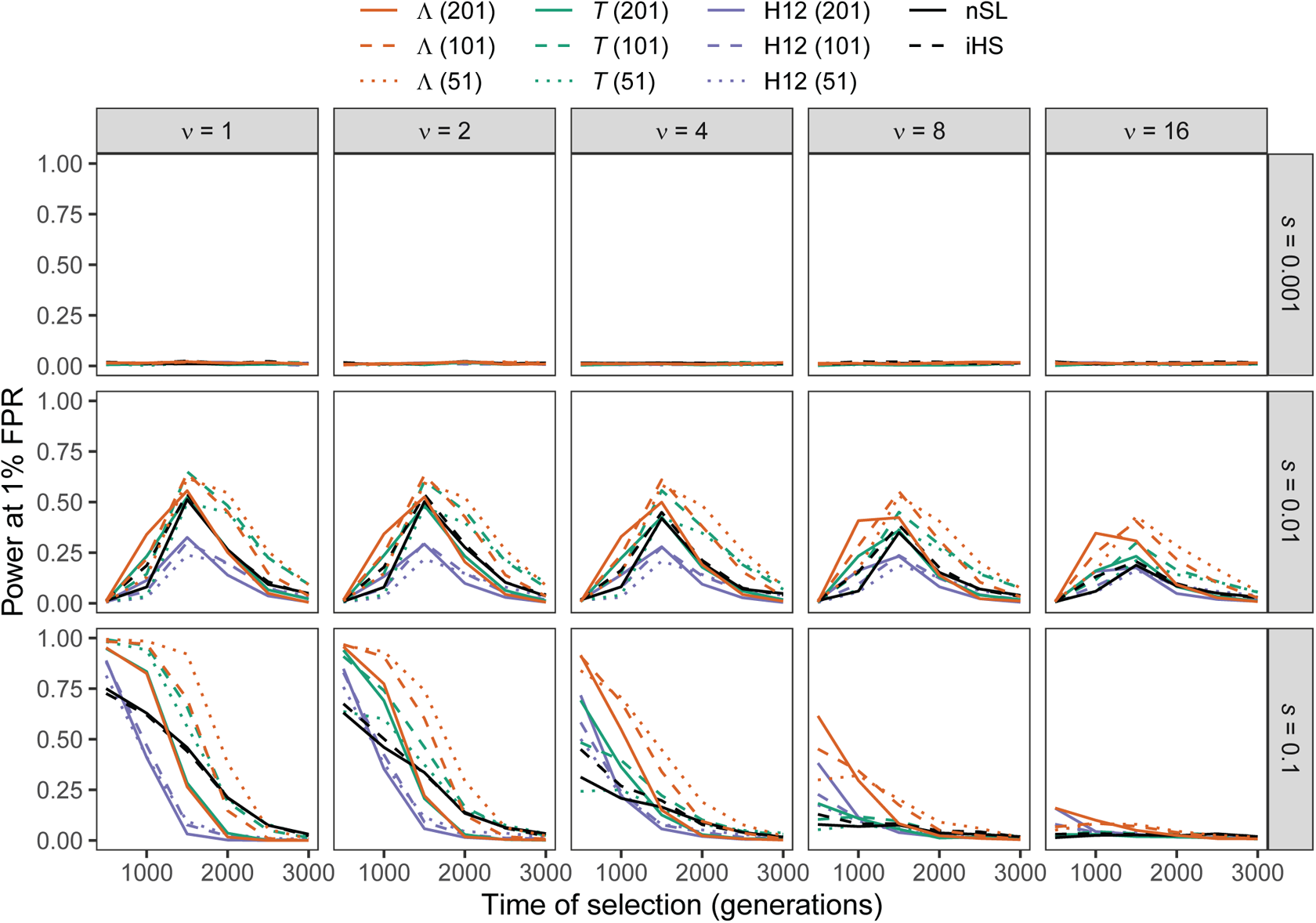
Power at a 1% false positive rate (FPR) as a function of selection start time for applications of Λ, *T*, and H12 with windows of size 51, 101, and 201 SNPs, as well *nS_L_* and iHS under simulations of the human central European (CEU) demographic history of Terhorst et al. [2017] with per-site per-generation recombination rate drawn from an exponential distribution with mean of 10*^−^*^8^ for per-generation selection coefficients of *s ∈* {0.001, 0.01, 0.1} on the rows. Classification ability demonstrated for selection start times of *t ∈* {500, 1000, 1500, 2000, 2500, 3000} generations prior to sampling for *ν ∈* {1, 2, 4, 8, 16} initially-selected haplotypes (columns). Results are based on a sample of *n* = 50 diploid individuals and the haplotype frequency spectra for the Λ and *T* statistics truncated at *K* = 10 haplotypes. Plots displaying patterns in estimated sweep softness and footprint size can be found in Figures S14 and S15, respectively.

**Figure S14:**
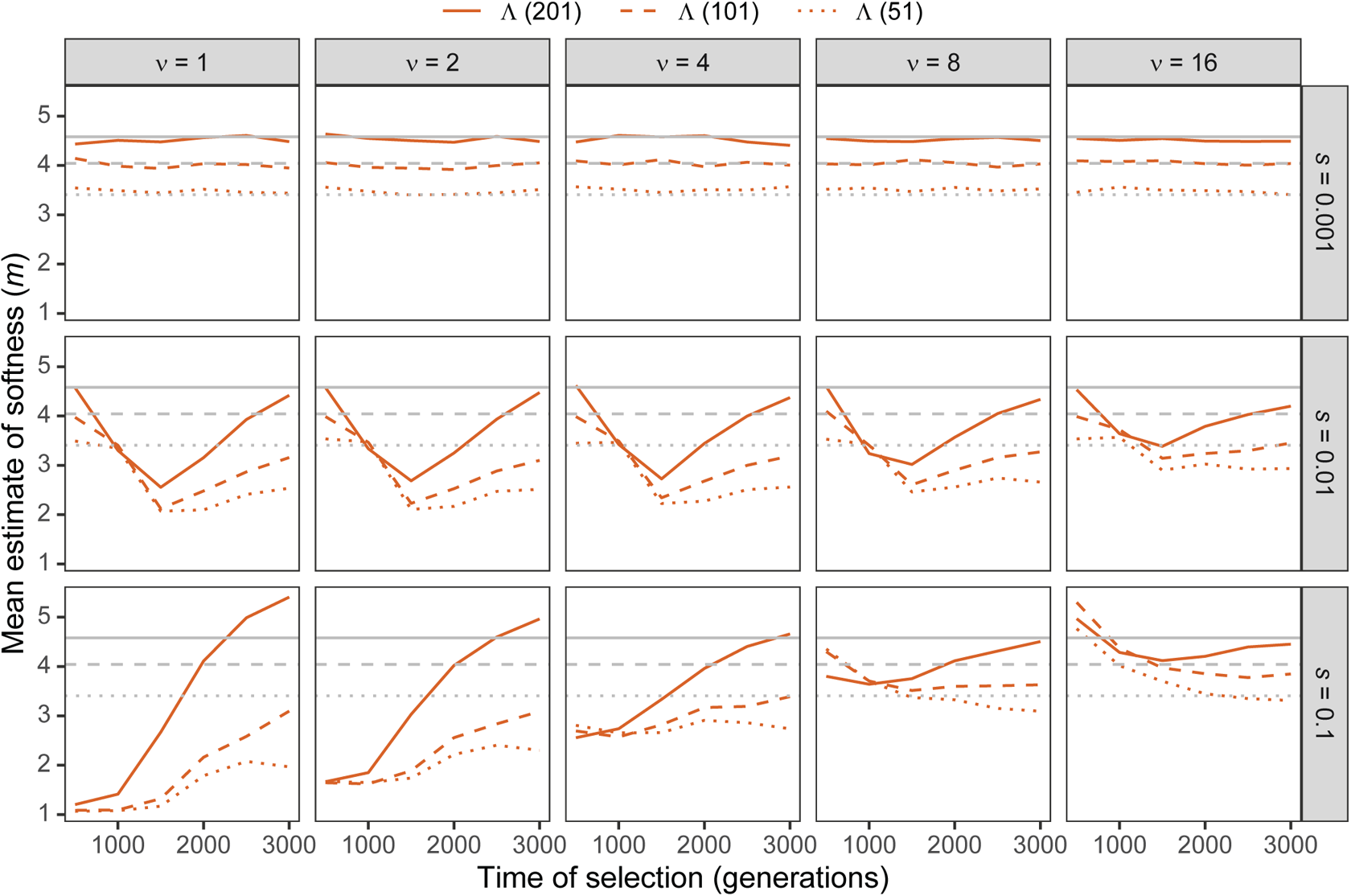
Estimated sweep softness illustrated by mean estimated number of sweeping haplotypes (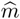) in Λ with windows of size 51, 101, and 201 SNPs under simulations of the human central European (CEU) demographic history of Terhorst et al. [2017] with per-site per-generation recombination rate drawn from an exponential distribution with mean of 10*^−^*^8^ for per-generation selection coefficients of *s ∈* {0.001, 0.01, 0.1} on the rows. Mean estimated softness demonstrated for selection start times of *t ∈* {500, 1000, 1500, 2000, 2500, 3000} generations prior to sampling for *ν ∈* {1, 2, 4, 8, 16} initially-selected haplotypes (columns). Gray solid, dashed, and dotted horizontal lines are the corresponding mean 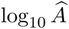 values for Λ applied to neutral simulations. Results are based on a sample of *n* = 50 diploid individuals and the haplotype frequency spectrum for the Λ statistic truncated at *K* = 10 haplotypes.

**Figure S15:**
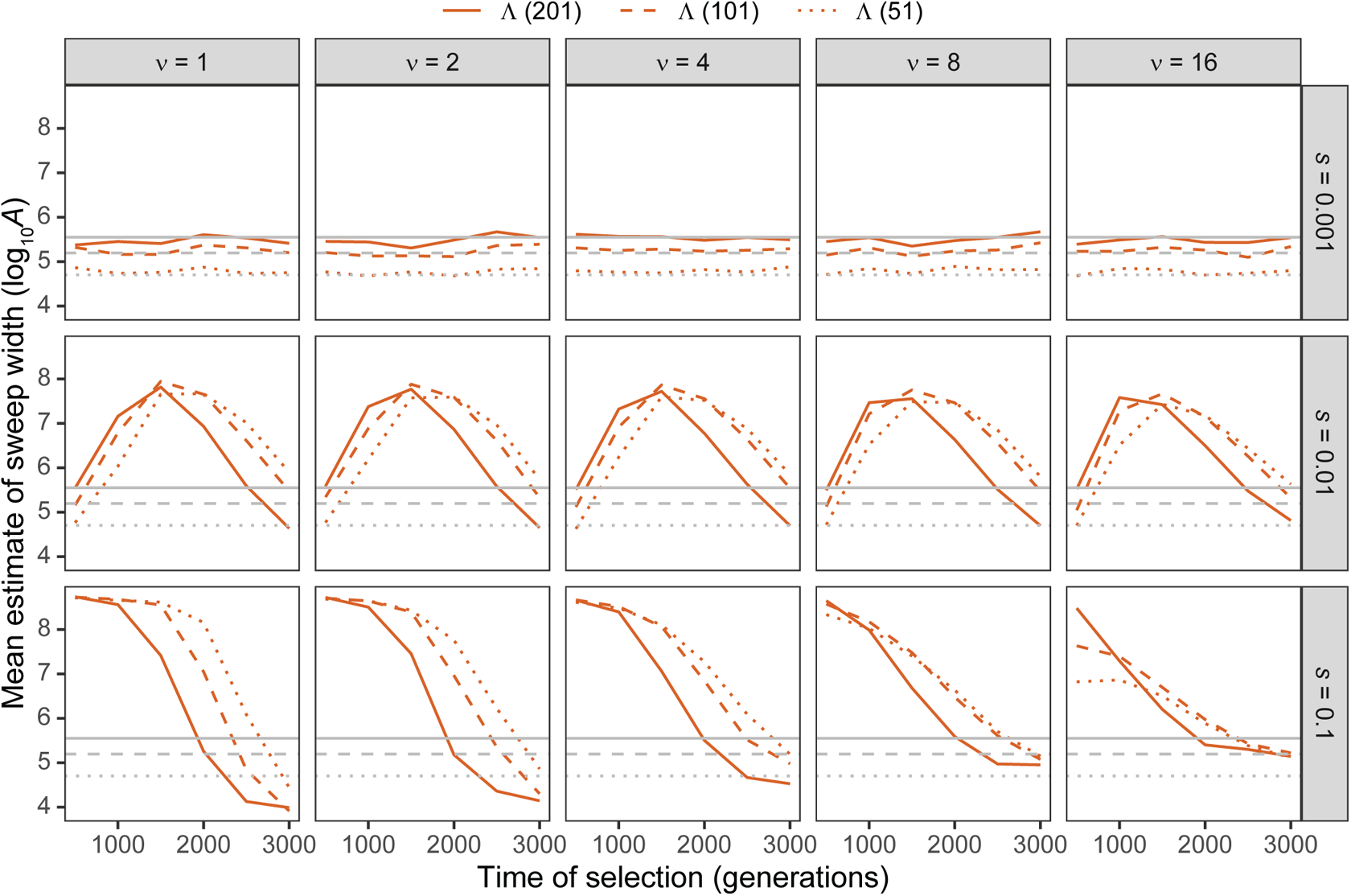
Estimated sweep width illustrated by mean estimated genomic size influenced by the sweep (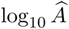) in Λ with windows of size 51, 101, and 201 SNPs under simulations of the human central European (CEU) demographic history of Terhorst et al. [2017] with per-site per-generation recombination rate drawn from an exponential distribution with mean of 10*^−^*^8^ for per-generation selection coefficients of *s ∈* {0.001, 0.01, 0.1} on the rows. Mean estimated genomic size influenced by sweeps demonstrated for selection start times of *t ∈* {500, 1000, 1500, 2000, 2500, 3000} generations prior to sampling for *ν ∈* {1, 2, 4, 8, 16} initially-selected haplotypes (columns). Gray solid, dashed, and dotted horizontal lines are the corresponding mean 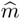 values for Λ applied to neutral simulations. Results are based on a sample of *n* = 50 diploid individuals and the haplotype frequency spectrum for the Λ statistic truncated at *K* = 10 haplotypes.

**Figure S16:**
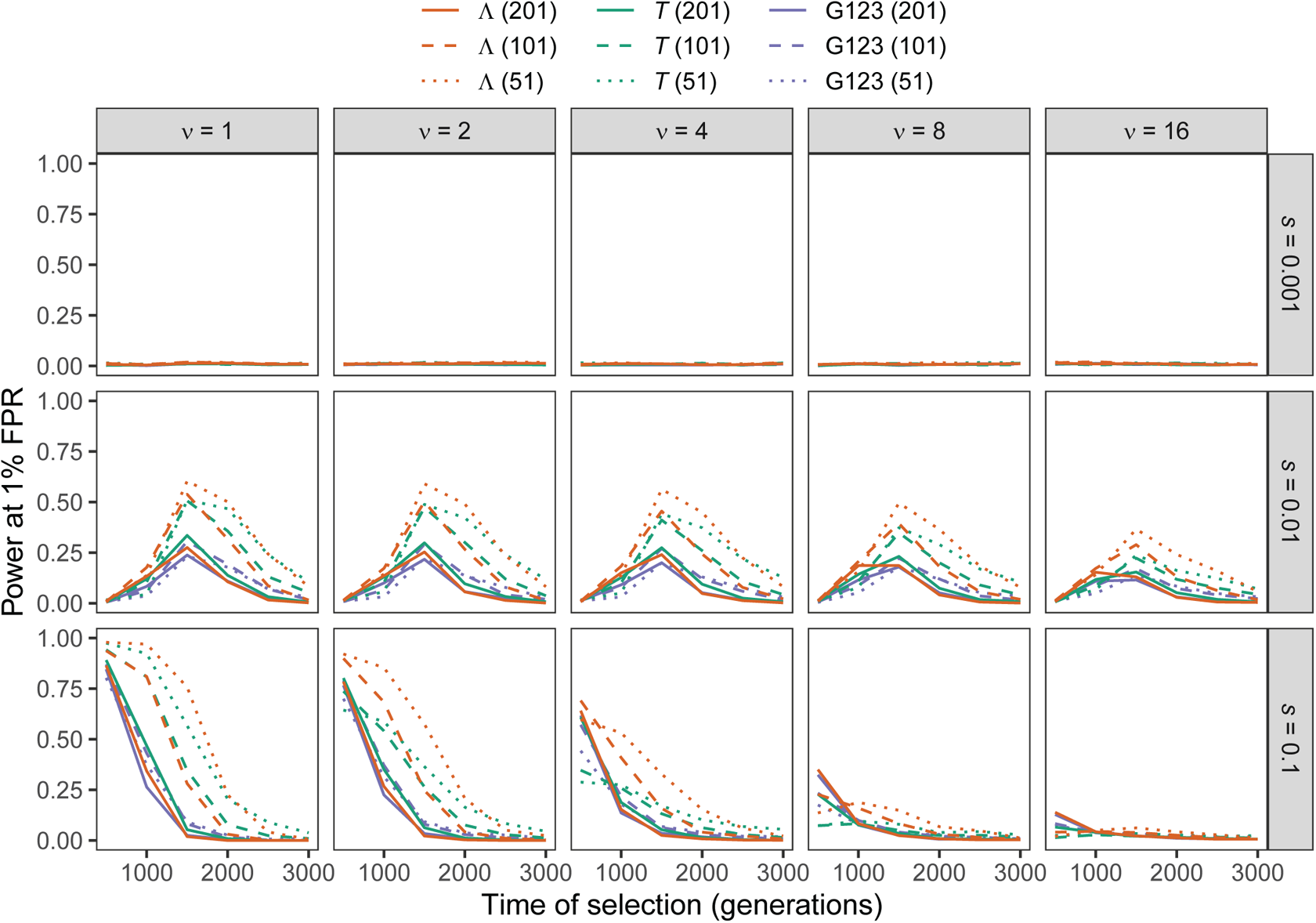
Power at a 1% false positive rate (FPR) as a function of selection start time for applications of Λ, *T*, and G123 with windows of size 51, 101, and 201 SNPs to unphased multilocus genotype input data under simulations of the human central European (CEU) demographic history of Terhorst et al. [2017] with per-site per-generation recombination rate drawn from an exponential distribution with mean of 10*^−^*^8^ for per-generation selection coefficients of *s* 0.001, 0.01, 0.1 on the rows. Classification ability demonstrated for selection start times of *t ∈* {500, 1000, 1500, 2000, 2500, 3000} generations prior to sampling for *ν ∈* {1, 2, 4, 8, 16} initially-selected haplotypes (columns). Results are based on a sample of *n* = 50 diploid individuals and the multilocus genotype frequency spectra for the Λ and *T* statistics truncated at *K* = 10 multilocus genotypes. Plots displaying patterns in estimated sweep softness and footprint size can be found in Figures S17 and S18, respectively.

**Figure S17:**
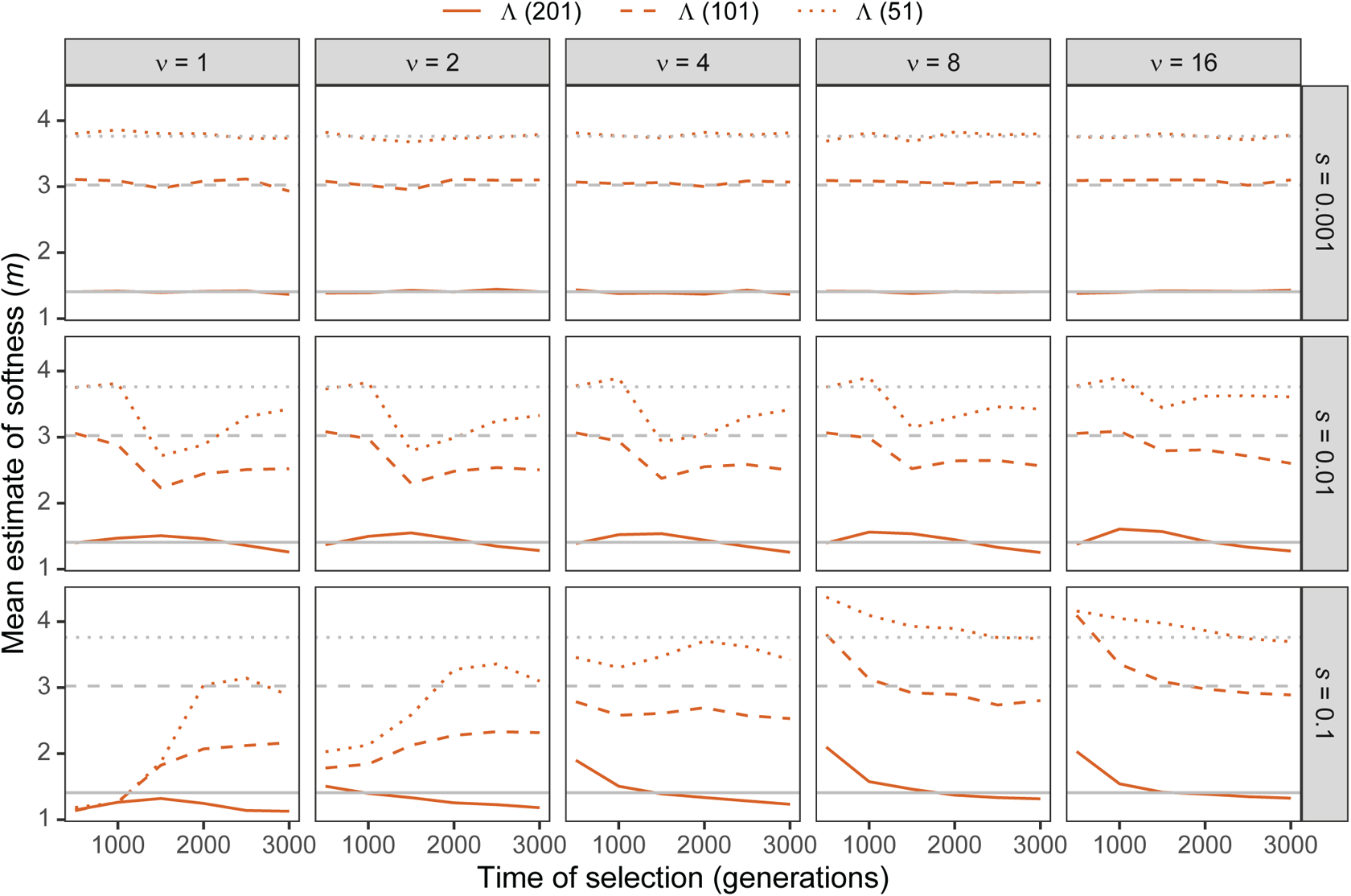
Estimated sweep softness illustrated by mean estimated number of sweeping haplotypes (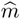) in Λ with windows of size 51, 101, and 201 SNPs applied to unphased multilocus input data under simulations of the human central European (CEU) demographic history of Terhorst et al. [2017] with per-site per-generation recombination rate drawn from an exponential distribution with mean of 10*^−^*^8^ for per-generation selection coefficients of *s ∈* {0.001, 0.01, 0.1} on the rows. Mean estimated softness demonstrated for selection start times of *t ∈* {500, 1000, 1500, 2000, 2500, 3000} generations prior to sampling for *ν ∈* {1, 2, 4, 8, 16} initially-selected haplotypes (columns). Gray solid, dashed, and dotted horizontal lines are the corresponding mean 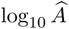 values for Λ applied to neutral simulations. Results are based on a sample of n = 50 diploid individuals and the multilocus genotype frequency spectrum for the Λ statistic truncated at K = 10 multilocus genotypes.

**Figure S18:**
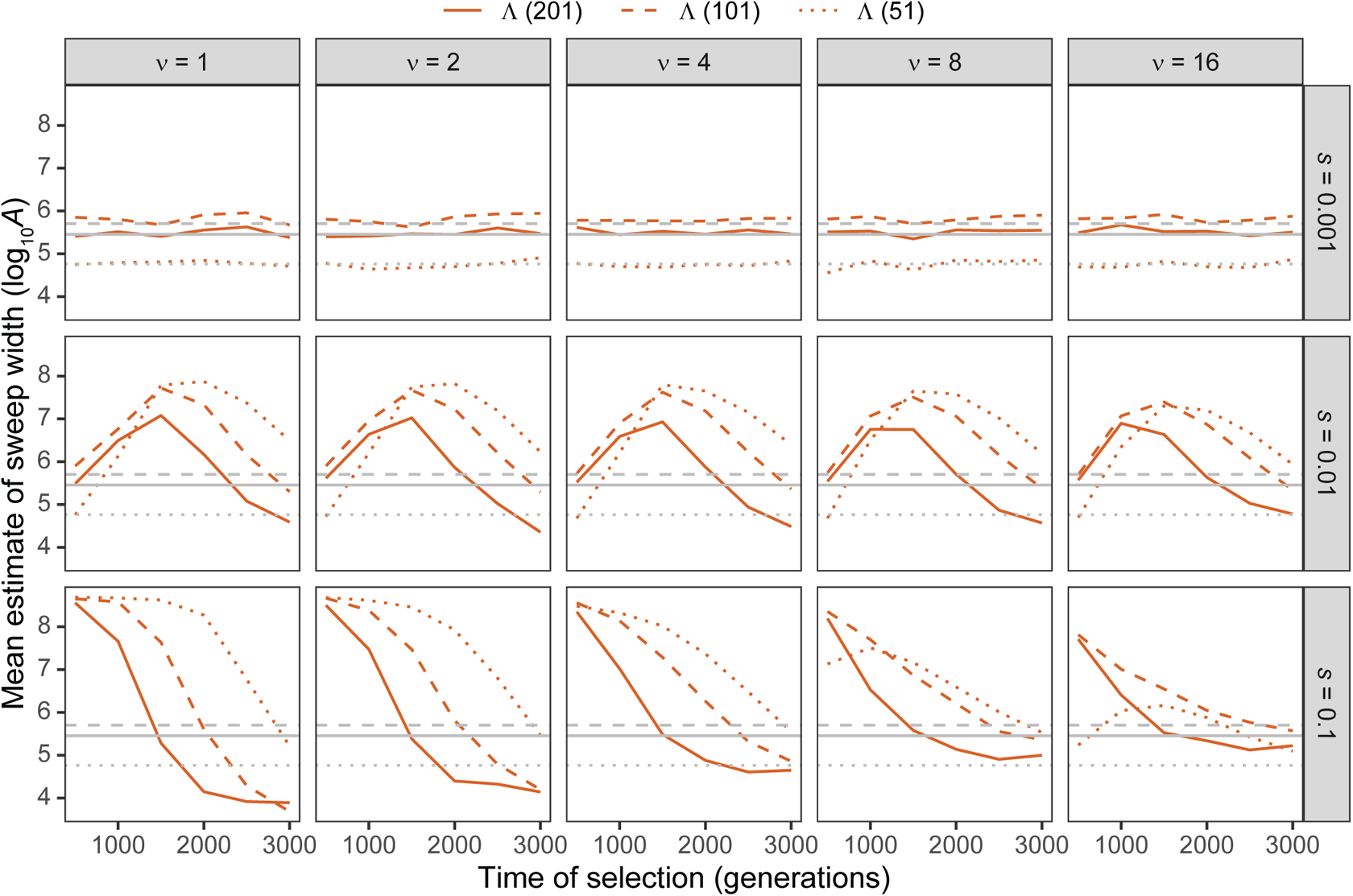
Estimated sweep width illustrated by mean estimated genomic size influenced by the sweep (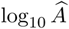) in Λ with windows of size 51, 101, and 201 SNPs applied to unphased multilocus input data under simulations of the human central European (CEU) demographic history of Terhorst et al. [2017] with per-site per-generation recombination rate drawn from an exponential distribution with mean of 10*^−^*^8^ for per-generation selection coefficients of *s ∈* {0.001, 0.01, 0.1} on the rows. Mean estimated genomic size influenced by sweeps demonstrated for selection start times of *t ∈* {500, 1000, 1500, 2000, 2500, 3000} generations prior to sampling for *ν ∈* {1, 2, 4, 8, 16} initially-selected haplotypes (columns). Gray solid, dashed, and dotted horizontal lines are the corresponding mean 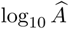 values for Λ applied to neutral simulations. Results are based on a sample of *n* = 50 diploid individuals and the multilocus genotype frequency spectrum for the Λ statistic truncated at *K* = 10 multilocus genotypes.

**Figure S19:**
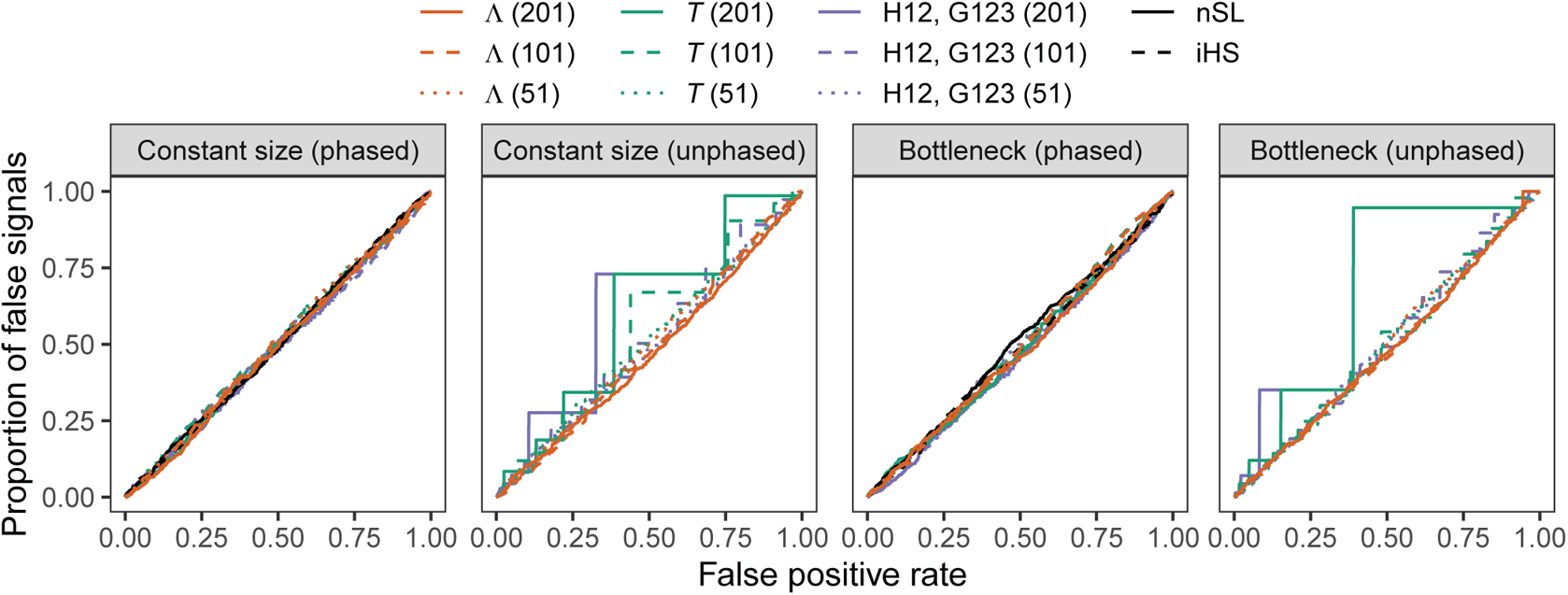
Proportion of false signals as a function of false positive rate for applications of Λ, *T*, H12, and G123 with windows of size 51, 101, and 201 SNPs, as well *nS_L_* and iHS under simulations of a constant-size demographic history and the human central European (CEU) demographic history of Terhorst et al. [2017] (bottleneck scenario) under background selection using either phased haplotype input data (Λ, *T*, H12, *nS_L_*, and iHS) or unphased multilocus genotype input data (Λ, *T*, and G123). Proportion of false signals is computed as the fraction of background selection simulations in which the score computed for Λ, *T*, H12, G123, *nS_L_*, or iHS exceeded the corresponding score threshold defined by a particular false positive rate. Results are based on a sample of *n* = 50 diploid individuals and haplotype and multilocus genotype frequency spectra for the Λ and *T* statistics truncated at *K* = 10 haplotypes or multilocus genotypes.

**Figure S20:**
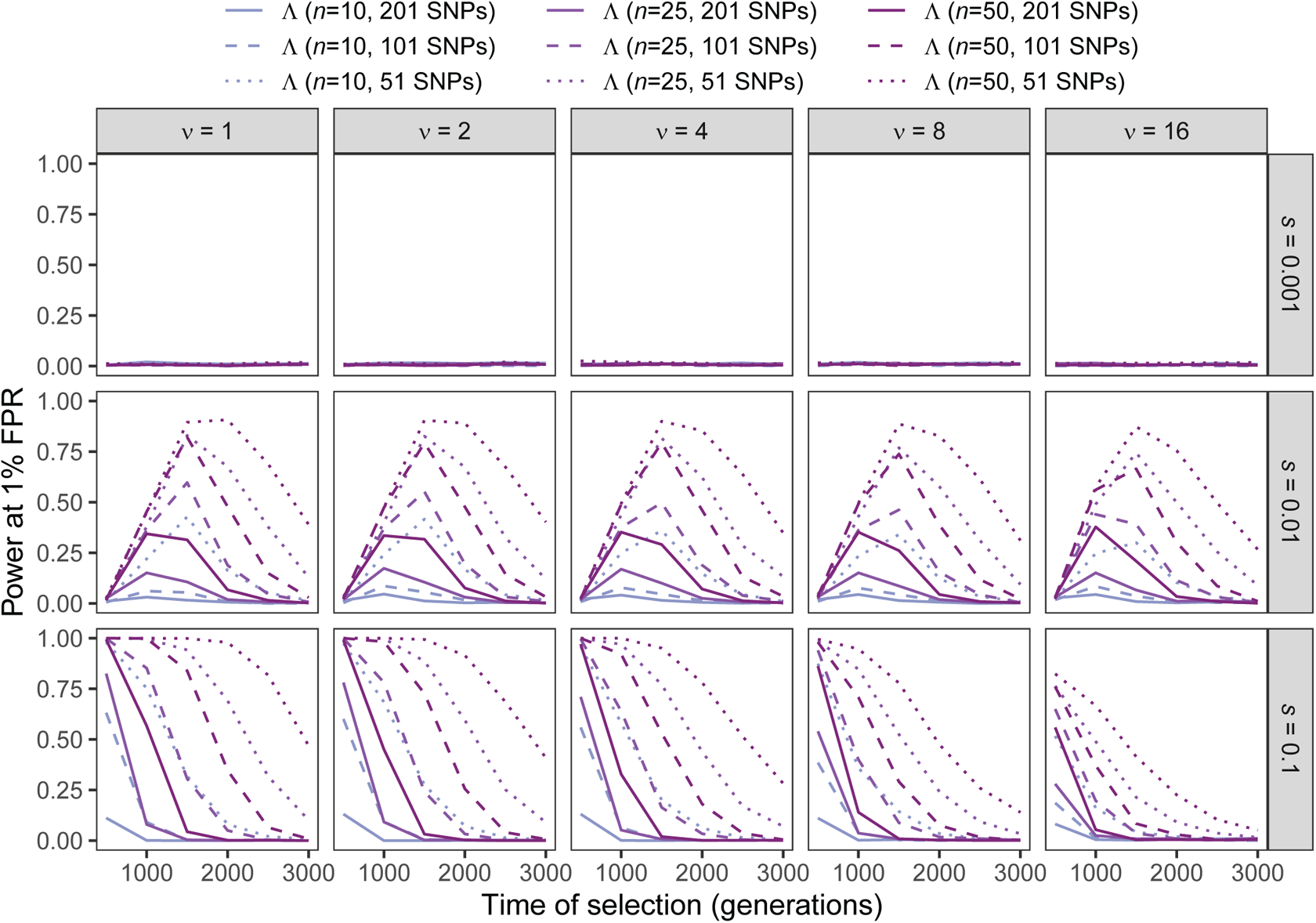
Power at a 1% false positive rate (FPR) as a function of selection start time for applications of Λ with windows of size 51, 101, and 201 SNPs under simulations of a constant-size demographic history and sample size of *n* ∈ {10, 25, 50} diploid individuals for per-generation selection coefficients of *s ∈* {0.001, 0.01, 0.1} on the rows. Classification ability demonstrated for selection start times of *t ∈* {500, 1000, 1500, 2000, 2500, 3000} generations prior to sampling for *ν ∈* {1, 2, 4, 8, 16} initially-selected haplotypes (columns). Results are based on the haplotype frequency spectra for the Λ statistics truncated at *K* = 10 haplotypes.

**Figure S21:**
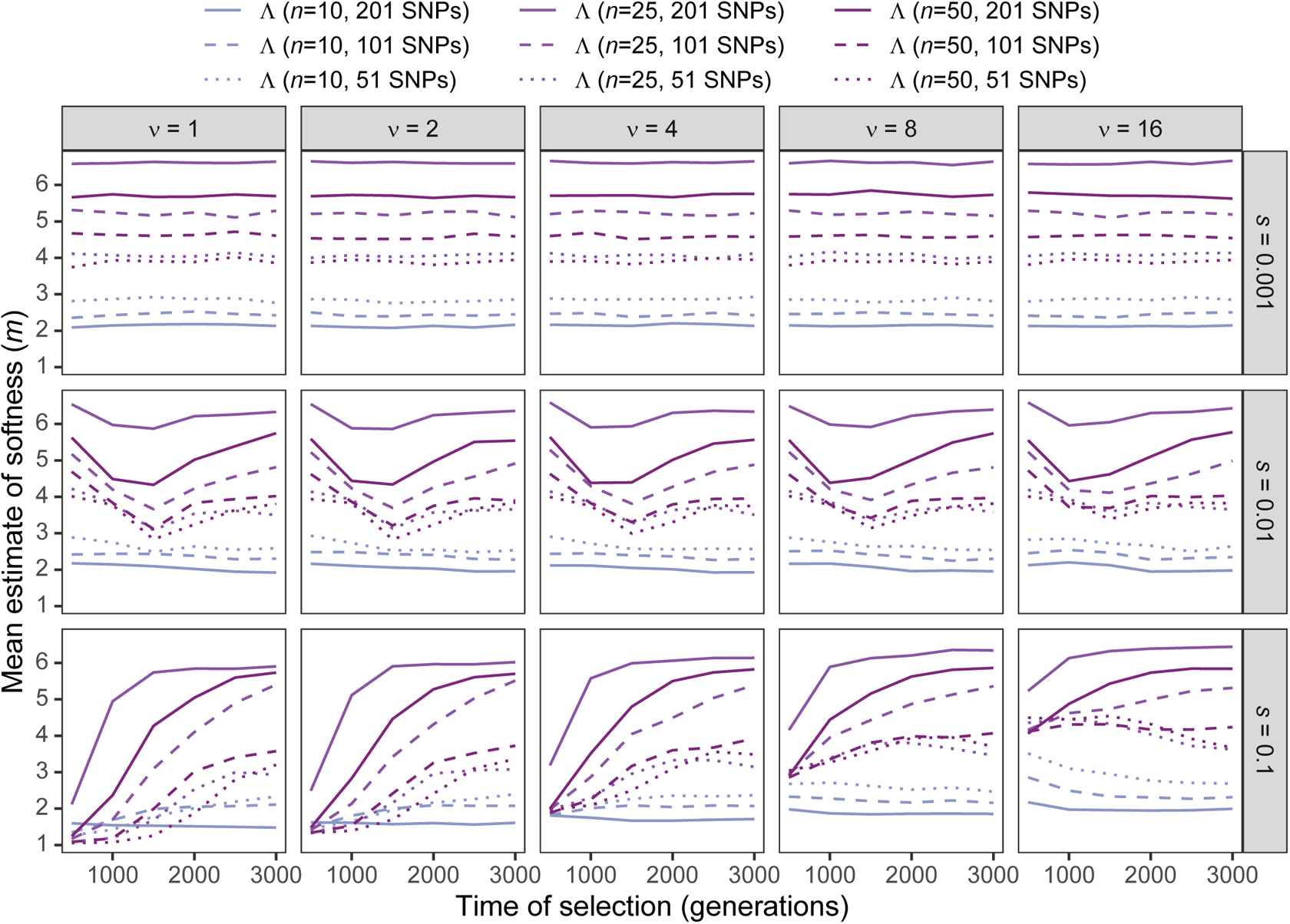
Estimated sweep softness illustrated by mean estimated number of sweeping haplo-types (*m*) in Λ with windows of size 51, 101, and 201 SNPs under simulations of a constant-size demographic history and sample size of *n ∈* {10, 25, 50} diploid individuals for per-generation selection coefficients of *s ∈* {0.001, 0.01, 0.1} on the rows. Mean estimated softness demonstrated for selection start times of *t ∈* {500, 1000, 1500, 2000, 2500, 3000} generations prior to sampling for *ν ∈* {1, 2, 4, 8, 16} initially-selected haplotypes (columns). Results are based on the haplotype frequency spectrum for the Λ statistic truncated at *K* = 10 haplotypes.

**Figure S22:**
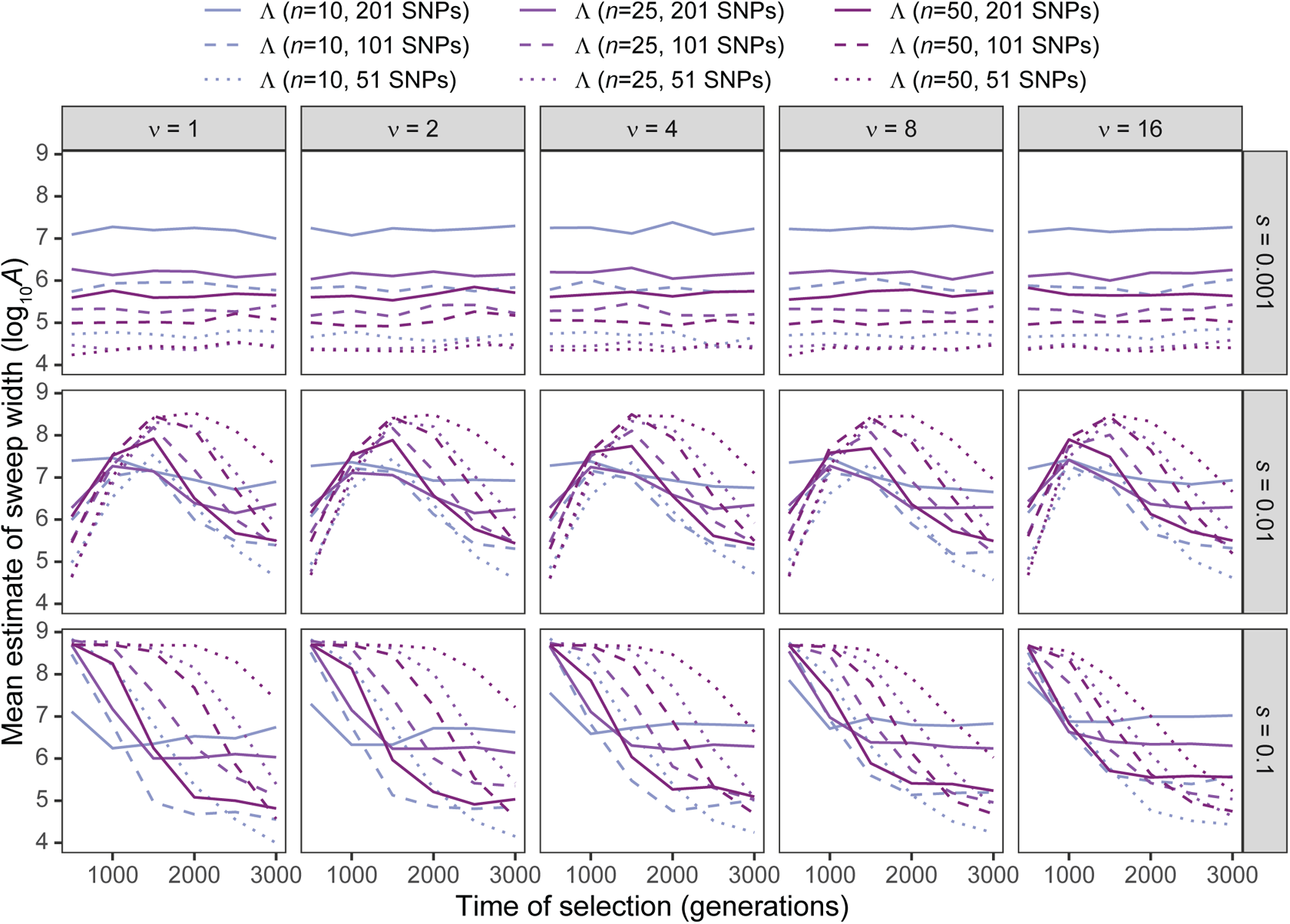
Estimated sweep width illustrated by mean estimated genomic size influenced by the sweep (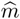) in Λ with windows of size 51, 101, and 201 SNPs under simulations of a constant-size demographic history and sample size of *n ∈* {10, 25, 50} diploid individuals for per-generation selection coefficients of *s ∈* {0.001, 0.01, 0.1} on the rows. Mean estimated genomic size influenced by sweeps demonstrated for selection start times of *t ∈* {500, 1000, 1500, 2000, 2500, 3000} generations prior to sampling for *ν ∈* {1, 2, 4, 8, 16} initially-selected haplotypes (columns). Results are based on the haplotype frequency spectrum for the Λ statistic truncated at *K* = 10 haplotypes.

**Figure S23:**
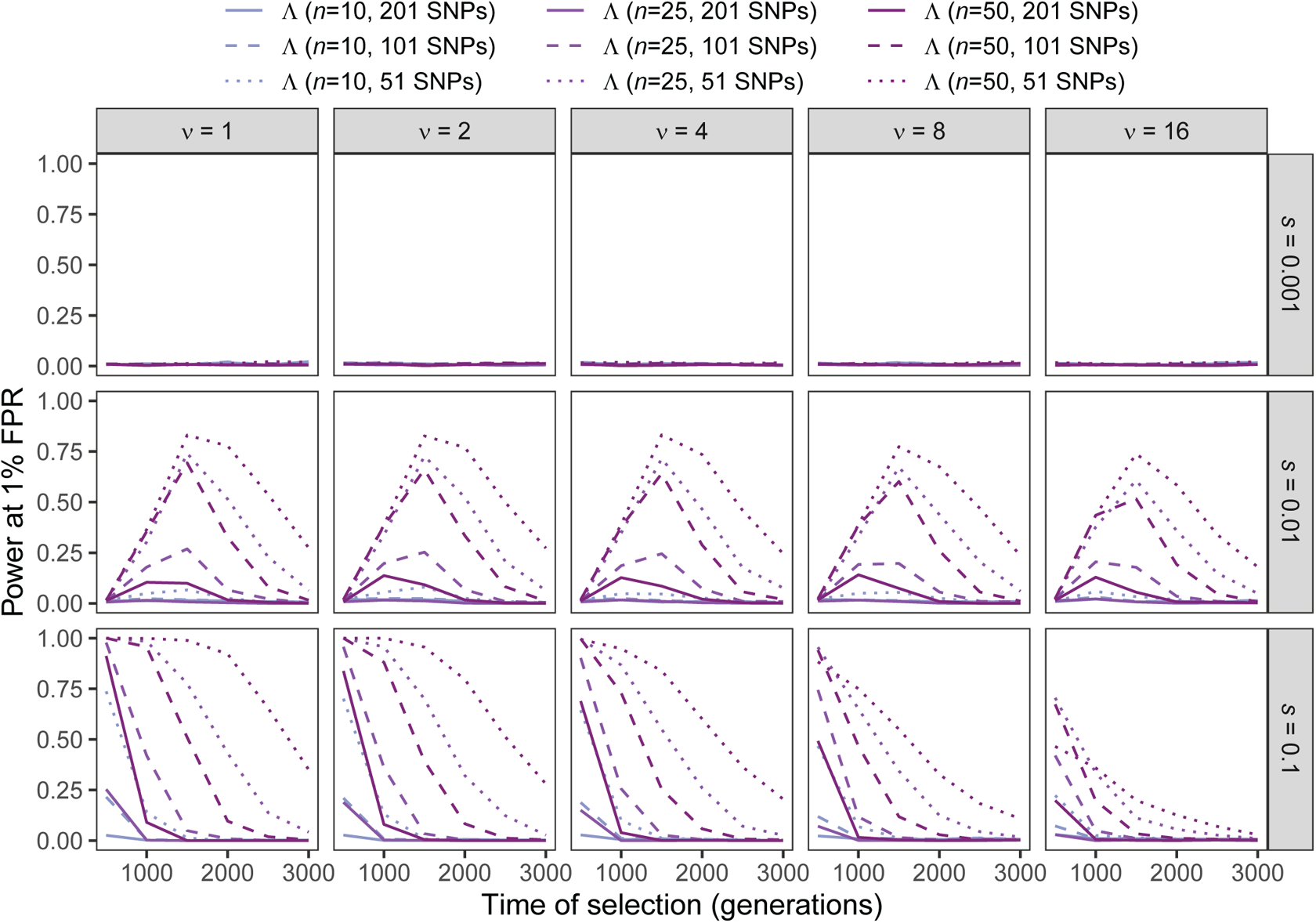
Power at a 1% false positive rate (FPR) as a function of selection start time for applications of Λ with windows of size 51, 101, and 201 SNPs to unphased multilocus genotype input data under simulations of a constant-size demographic history and sample size of *n ∈* {10, 25, 50} diploid individuals for per-generation selection coefficients of *s ∈* {0.001, 0.01, 0.1} on the rows. Classification ability demonstrated for selection start times of *t ∈* {500, 1000, 1500, 2000, 2500, 3000} generations prior to sampling for *ν ∈* {1, 2, 4, 8, 16} initially-selected haplotypes (columns). Results are based on the multilocus genotype frequency spectrum for the Λ statistic truncated at *K* = 10 multilocus genotypes.

**Figure S24:**
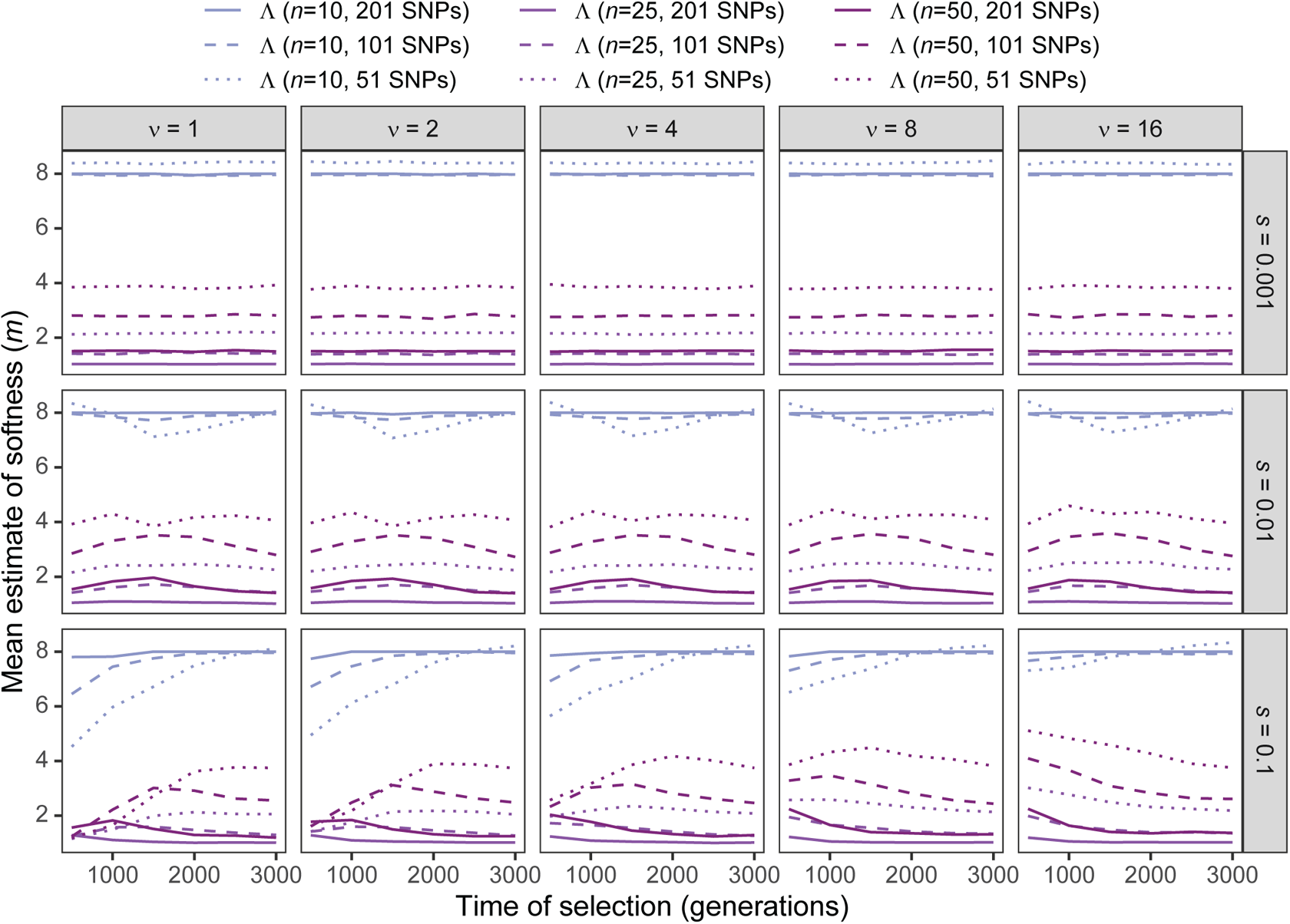
Estimated sweep softness illustrated by mean estimated number of sweeping haplotypes (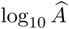) in Λ with windows of size 51, 101, and 201 SNPs applied to unphased multilocus input data under simulations of a constant-size demographic history and sample size of *n ∈* {10, 25, 50} diploid individuals for per-generation selection coefficients of *s ∈* {0.001, 0.01, 0.1} on the rows. Mean estimated softness demonstrated for selection start times of *t ∈* {500, 1000, 1500, 2000, 2500, 3000} generations prior to sampling for *ν ∈* {1, 2, 4, 8, 16} initially-selected haplotypes (columns). Results are based on the multilocus genotype frequency spectrum for the Λ statistic truncated at *K* = 10 multilocus genotypes.

**Figure S25:**
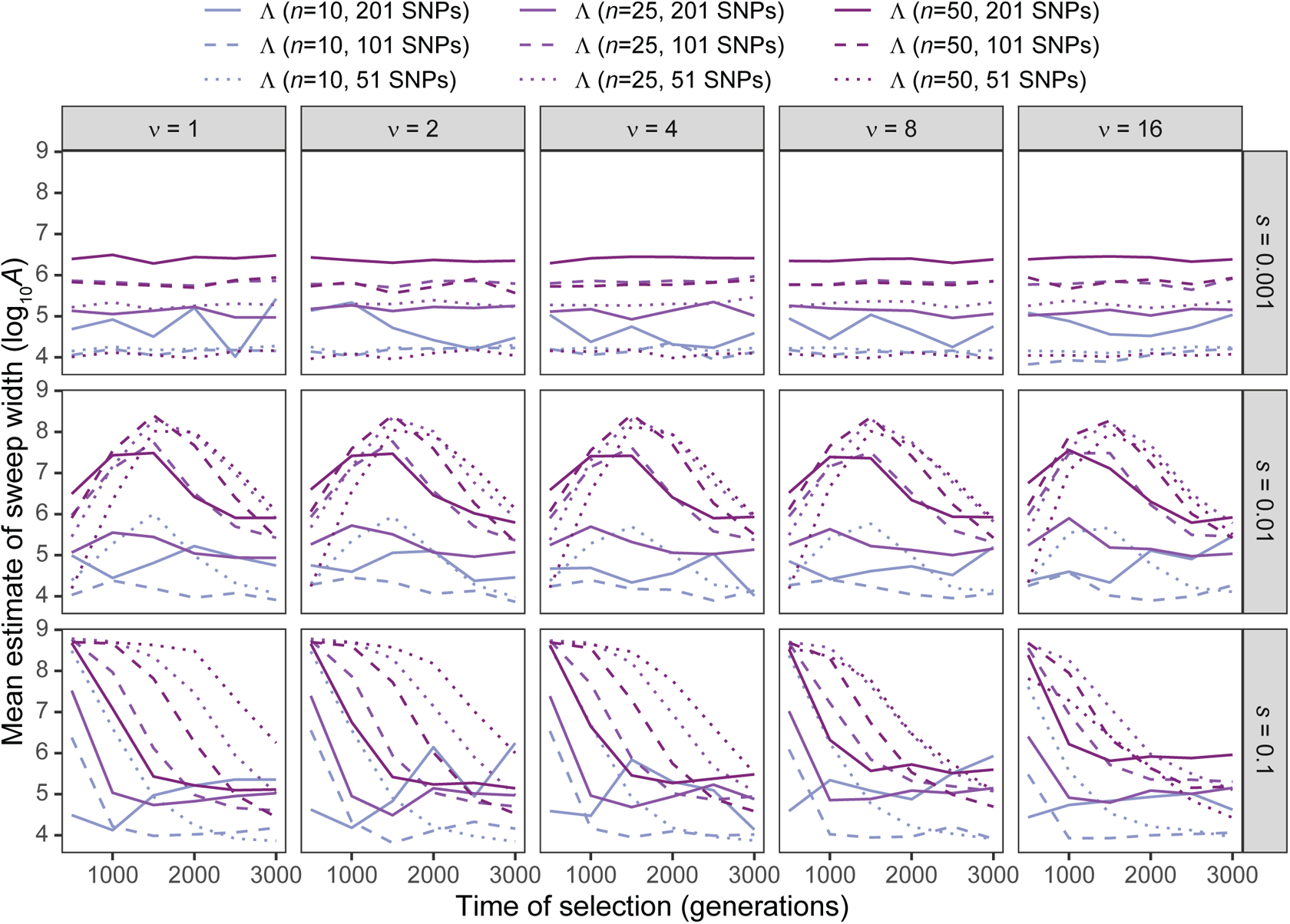
Estimated sweep width illustrated by mean estimated genomic size influenced by the sweep (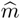) in Λ with windows of size 51, 101, and 201 SNPs applied to unphased multilocus input data under simulations of a constant-size demographic history and sample size of *n ∈* {10, 25, 50} diploid individuals for per-generation selection coefficients of *s ∈* {0.001, 0.01, 0.1} on the rows. Mean estimated genomic size influenced by sweeps demonstrated for selection start times of *t ∈* {500, 1000, 1500, 2000, 2500, 3000} generations prior to sampling for *ν ∈* {1, 2, 4, 8, 16} initially-selected haplotypes (columns). Results are based on the multilocus genotype frequency spectrum for the Λ statistic truncated at *K* = 10 multilocus genotypes.

**Figure S26:**
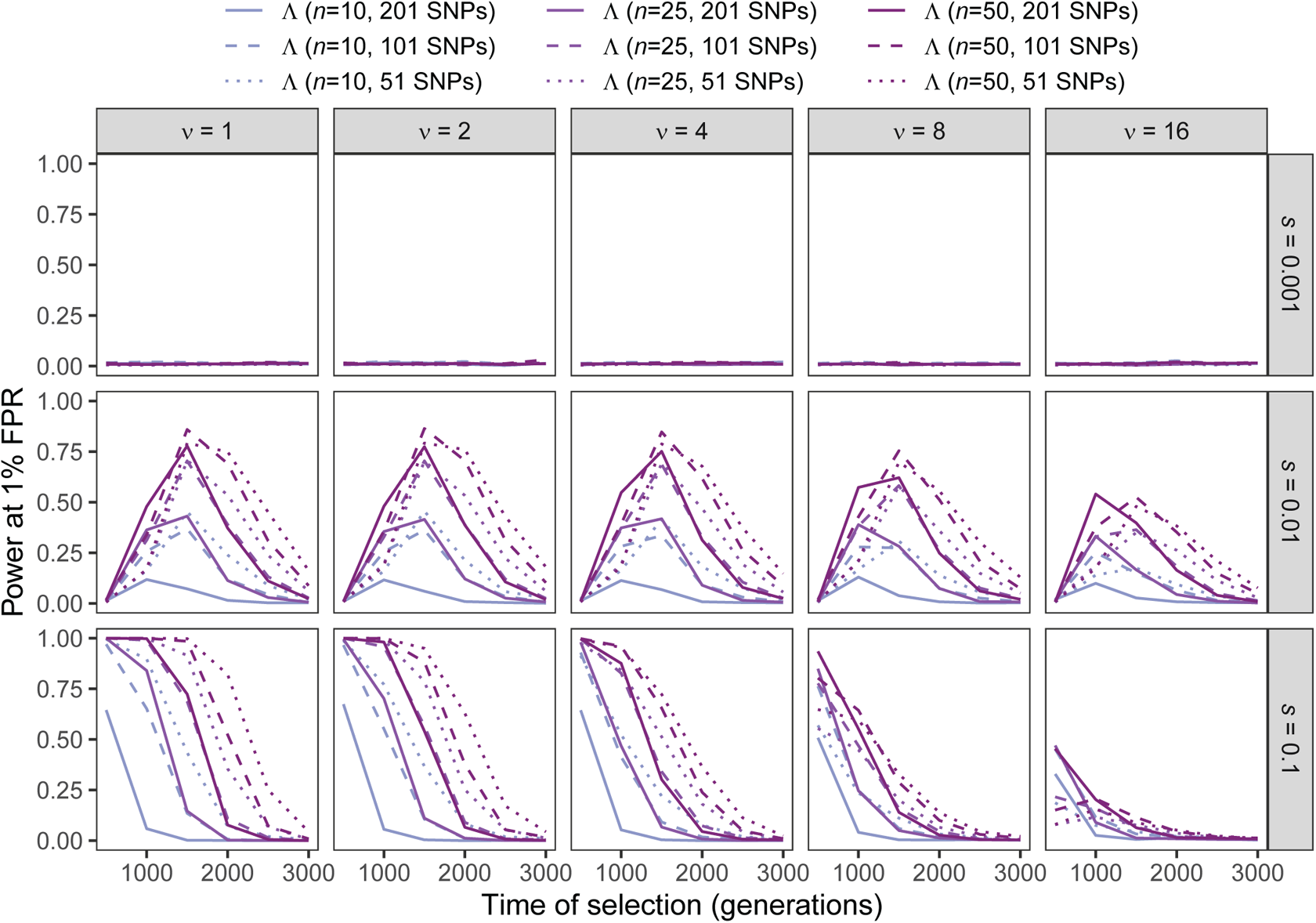
Power at a 1% false positive rate (FPR) as a function of selection start time for applications of Λ with windows of size 51, 101, and 201 SNPs under simulations of the human central European (CEU) demographic history of Terhorst et al. [2017] and sample size of *n ∈* {10, 25, 50} diploid individuals for per-generation selection coefficients of *s ∈* {0.001, 0.01, 0.1} on the rows. Classification ability demonstrated for selection start times of *t ∈* {500, 1000, 1500, 2000, 2500, 3000} generations prior to sampling for *ν ∈* {1, 2, 4, 8, 16} initially-selected haplotypes (columns). Results are based on the haplotype frequency spectra for the Λ statistics truncated at *K* = 10 haplotypes.

**Figure S27:**
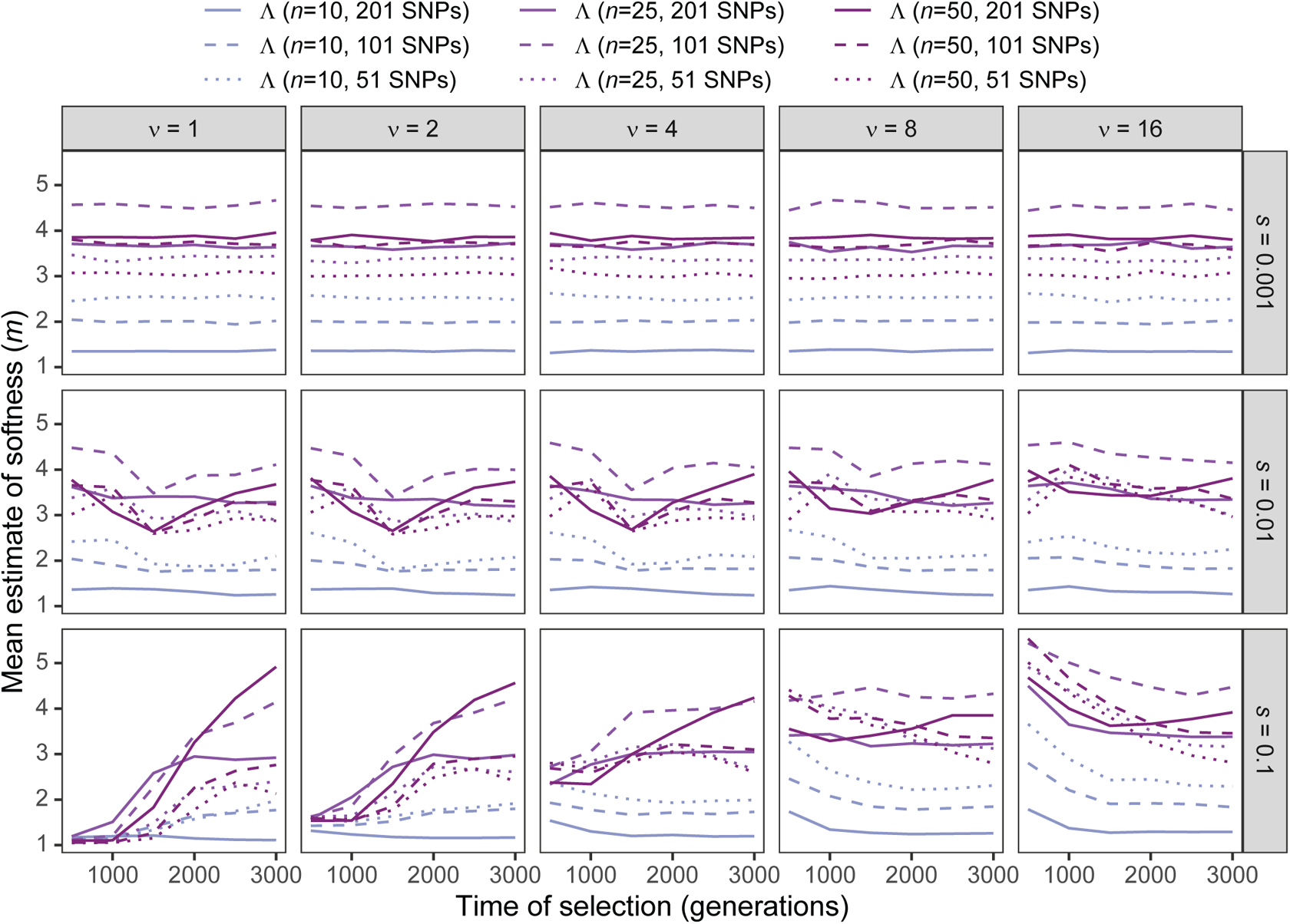
Estimated sweep softness illustrated by mean estimated number of sweeping haplotypes (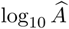) in Λ with windows of size 51, 101, and 201 SNPs under simulations of the human central European (CEU) demographic history of Terhorst et al. [2017] and sample size of *n ∈* {10, 25, 50} diploid individuals for per-generation selection coefficients of *s ∈* {0.001, 0.01, 0.1} on the rows. Mean estimated softness demonstrated for selection start times of *t ∈* {500, 1000, 1500, 2000, 2500, 3000} generations prior to sampling for *ν ∈* {1, 2, 4, 8, 16} initially-selected haplotypes (columns). Results are based on the haplotype frequency spectrum for the Λ statistic truncated at *K* = 10 haplotypes.

**Figure S28:**
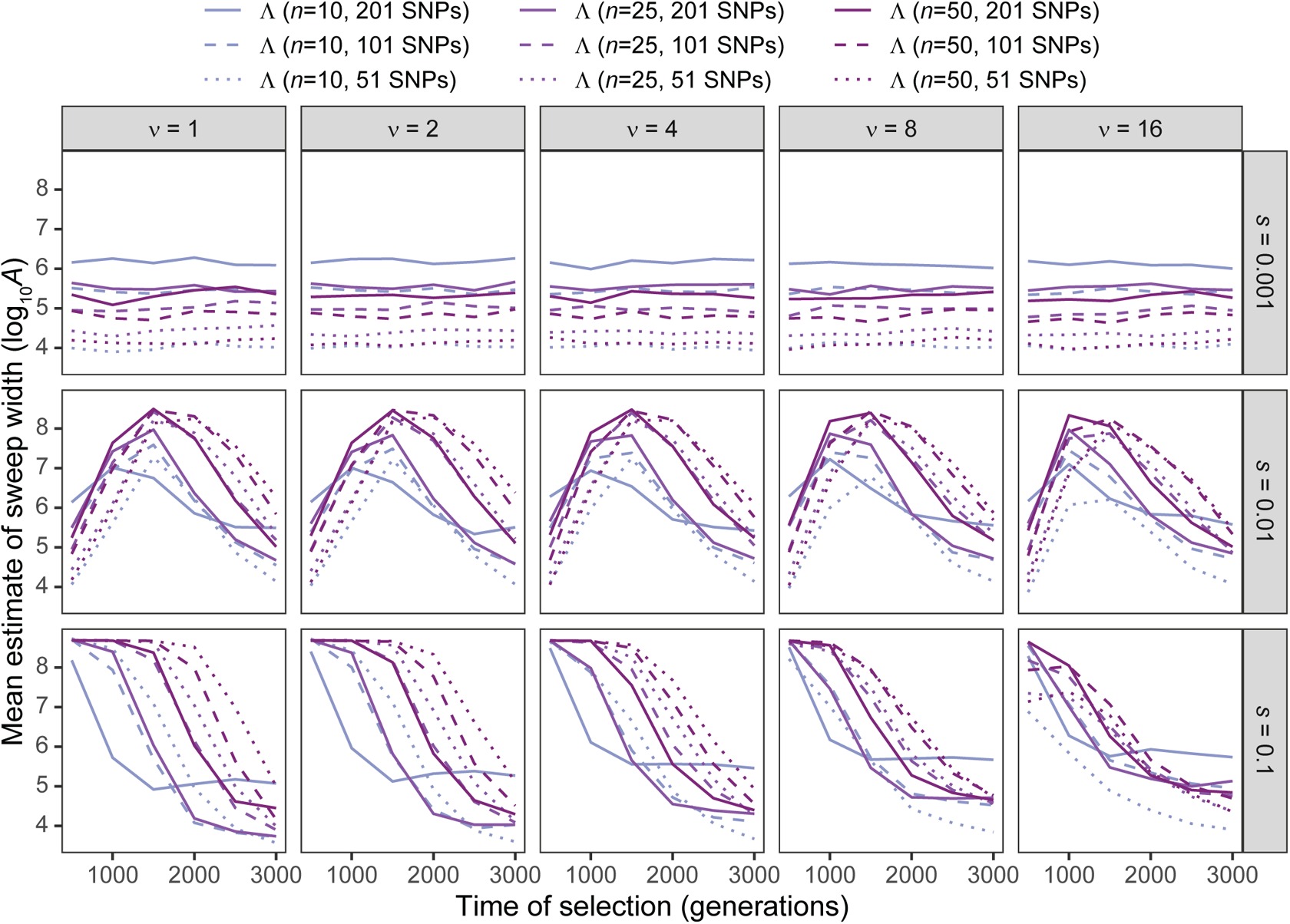
Estimated sweep width illustrated by mean estimated genomic size influenced by the sweep (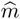) in Λ with windows of size 51, 101, and 201 SNPs under simulations of the human central European (CEU) demographic history of Terhorst et al. [2017] and sample size of *n ∈* {10, 25, 50} diploid individuals for per-generation selection coefficients of *s ∈* {0.001, 0.01, 0.1} on the rows. Mean estimated genomic size influenced by sweeps demonstrated for selection start times of *t ∈* {500, 1000, 1500, 2000, 2500, 3000} generations prior to sampling for *ν ∈* {1, 2, 4, 8, 16} initially-selected haplotypes (columns). Results are based on the haplotype frequency spectrum for the Λ statistic truncated at *K* = 10 haplotypes.

**Figure S29:**
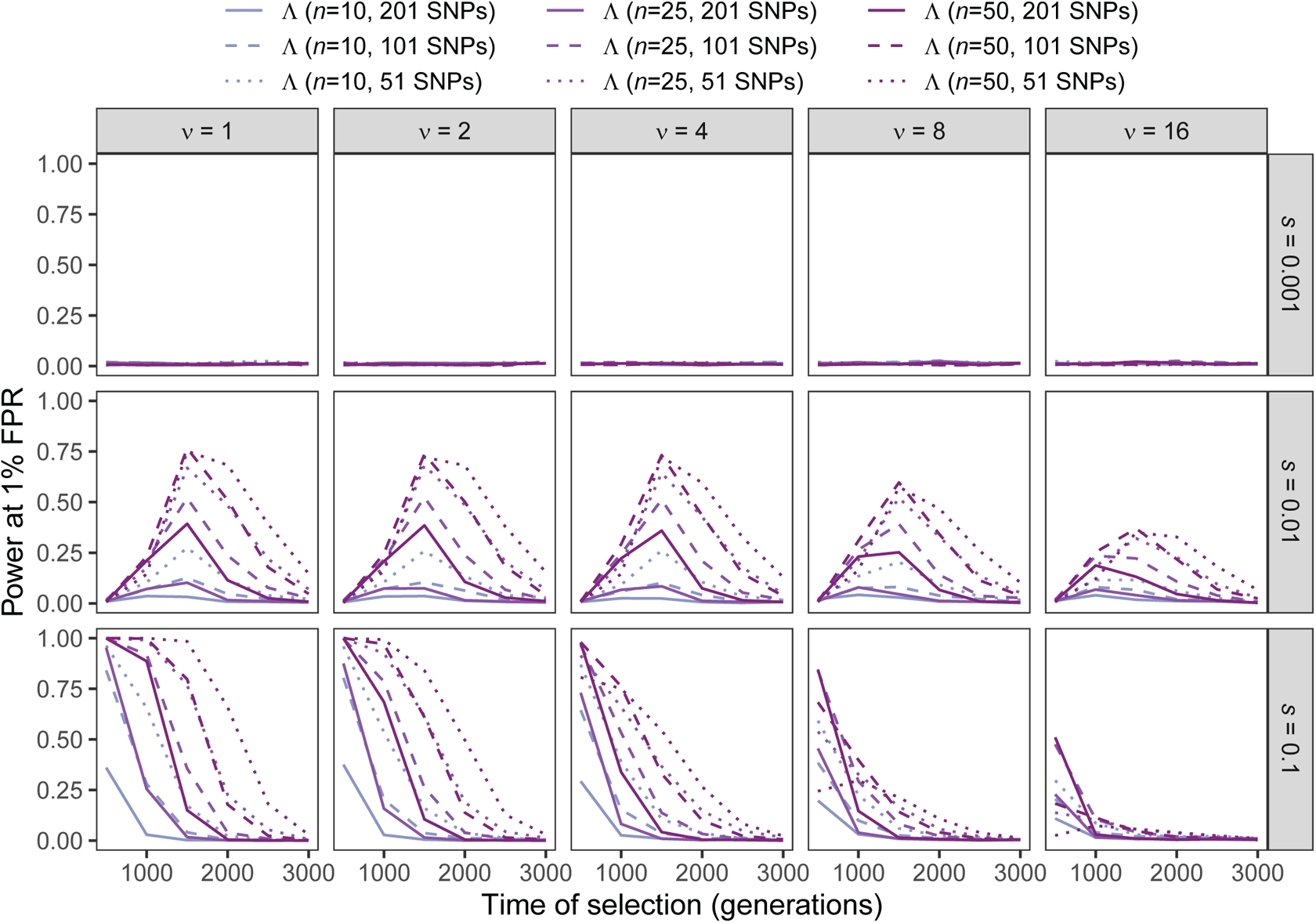
Power at a 1% false positive rate (FPR) as a function of selection start time for applications of Λ with windows of size 51, 101, and 201 SNPs to unphased multilocus genotype input data under simulations of the human central European (CEU) demographic history of Terhorst et al. [2017] and sample size of *n* ∈ {10, 25, 50} diploid individuals for per-generation selection coefficients of *s* 0.001, 0.01, 0.1 on the rows. Classification ability demonstrated for selection start times of *t* ∈ {500, 1000, 1500, 2000, 2500, 3000} generations prior to sampling for *ν ∈* {1, 2, 4, 8, 16} initially-selected haplotypes (columns). Results are based on the multilocus genotype frequency spectrum for the Λ statistic truncated at *K* = 10 multilocus genotypes.

**Figure S30:**
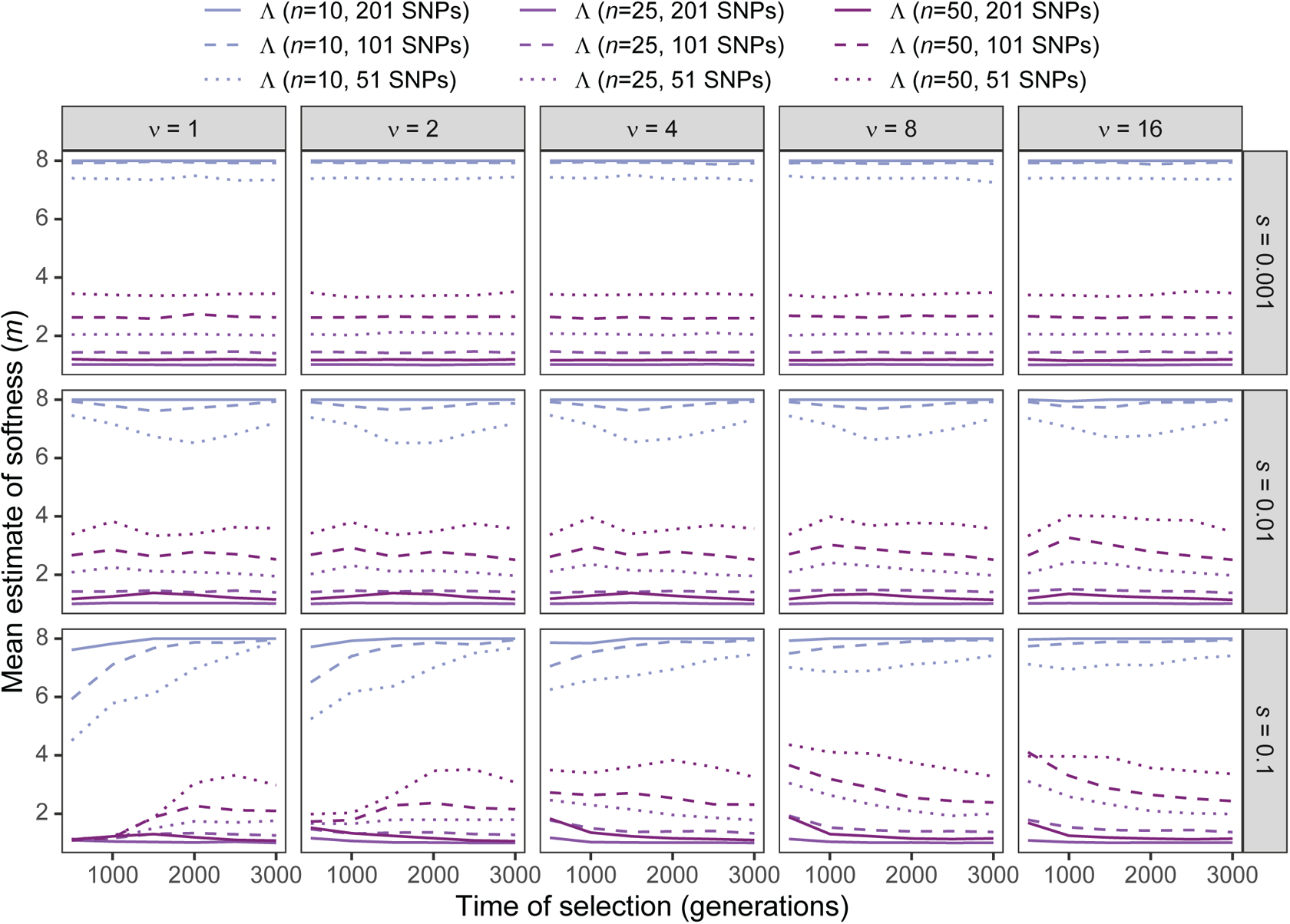
Estimated sweep softness illustrated by mean estimated number of sweeping haplotypes (*m*) in Λ with windows of size 51, 101, and 201 SNPs applied to unphased multilocus input data under simulations of the human central European (CEU) demographic history of Terhorst et al. [2017] and sample size of *n* ∈ {10, 25, 50} diploid individuals for per-generation selection coefficients of *s* 0.001, 0.01, 0.1 on the rows. Mean estimated softness demonstrated for selection start times of *t* ∈ {500, 1000, 1500, 2000, 2500, 3000} generations prior to sampling for *ν ∈* {1, 2, 4, 8, 16} initially-selected haplotypes (columns). Results are based on the multilocus genotype frequency spectrum for the Λ statistic truncated at *K* = 10 multilocus genotypes.

**Figure S31:**
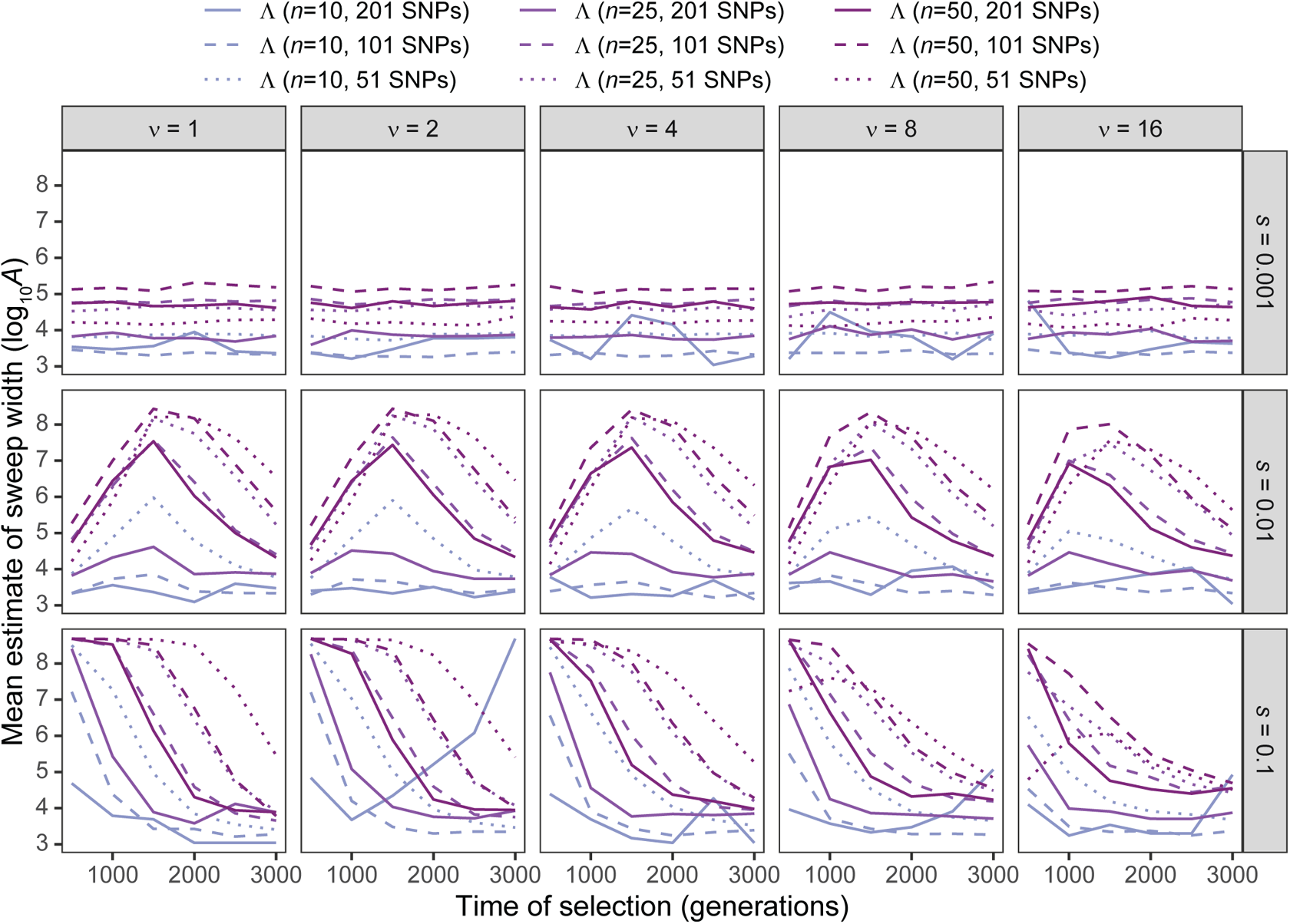
Estimated sweep width illustrated by mean estimated genomic size influenced by the sweep (log_10_ *A*) in Λ with windows of size 51, 101, and 201 SNPs applied to unphased multilocus input data under simulations of the human central European (CEU) demographic history of Terhorst et al. [2017] and sample size of *n* ∈ {10, 25, 50} diploid individuals for per-generation selection coefficients of *s* 0.001, 0.01, 0.1 on the rows. Mean estimated genomic size influenced by sweeps demonstrated for selection start times of *t* ∈ {500, 1000, 1500, 2000, 2500, 3000} generations prior to sampling for *ν ∈* {1, 2, 4, 8, 16} initially-selected haplotypes (columns). Results are based on the multilocus genotype frequency spectrum for the Λ statistic truncated at *K* = 10 multilocus genotypes.

**Figure S32:**
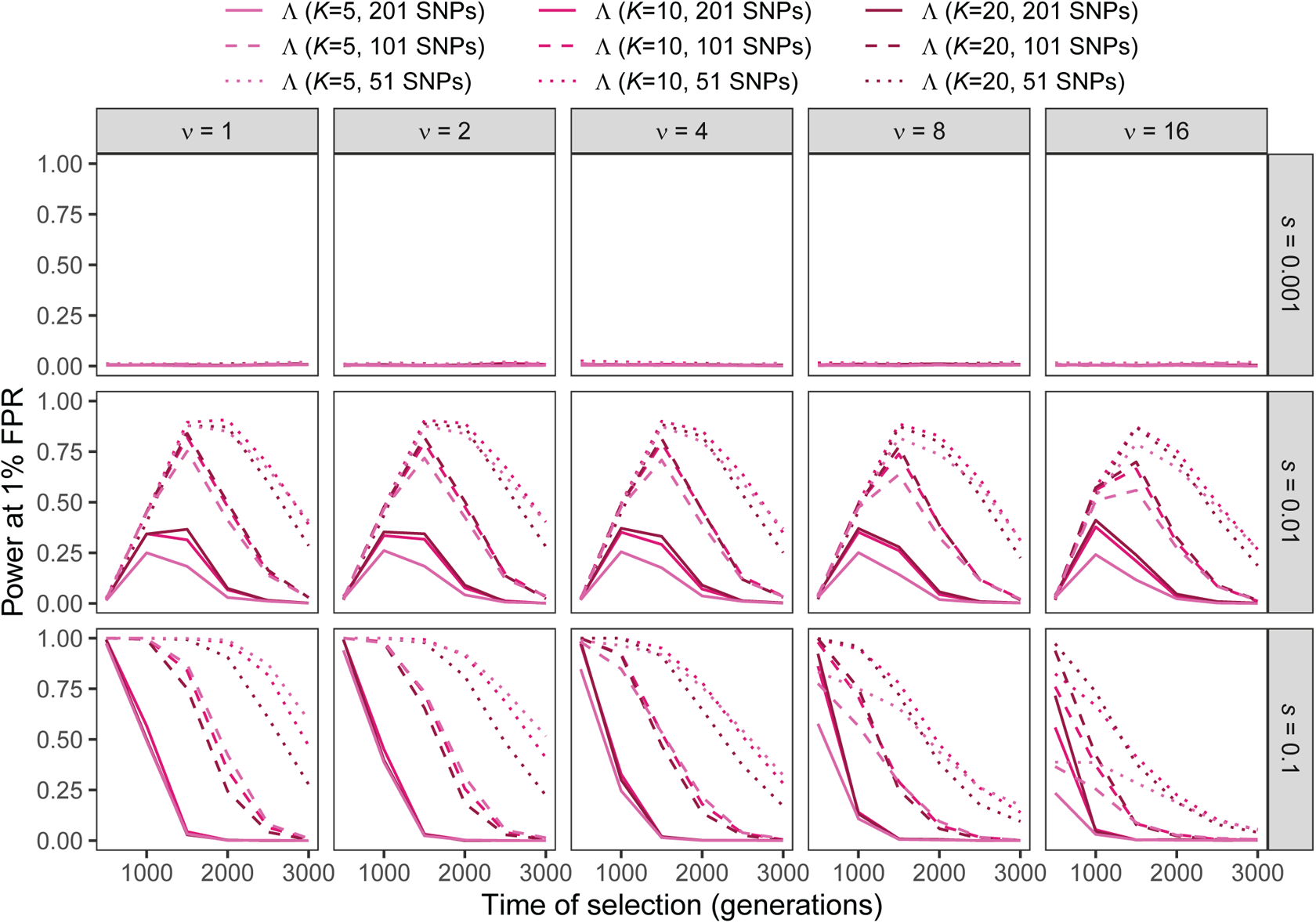
Power at a 1% false positive rate (FPR) as a function of selection start time for applications of Λ with windows of size 51, 101, and 201 SNPs under simulations of a constant-size demographic history and the haplotype frequency spectra for the Λ statistic truncated at *K* 5, 10, 20 haplotypes for per-generation selection coefficients of *s* 0.001, 0.01, 0.1 on the rows. Classification ability demonstrated for selection start times of *t* ∈ {500, 1000, 1500, 2000, 2500, 3000} generations prior to sampling for *ν ∈* {1, 2, 4, 8, 16} initially-selected haplotypes (columns). Results are based on a sample of *n* = 50 diploid individuals.

**Figure S33:**
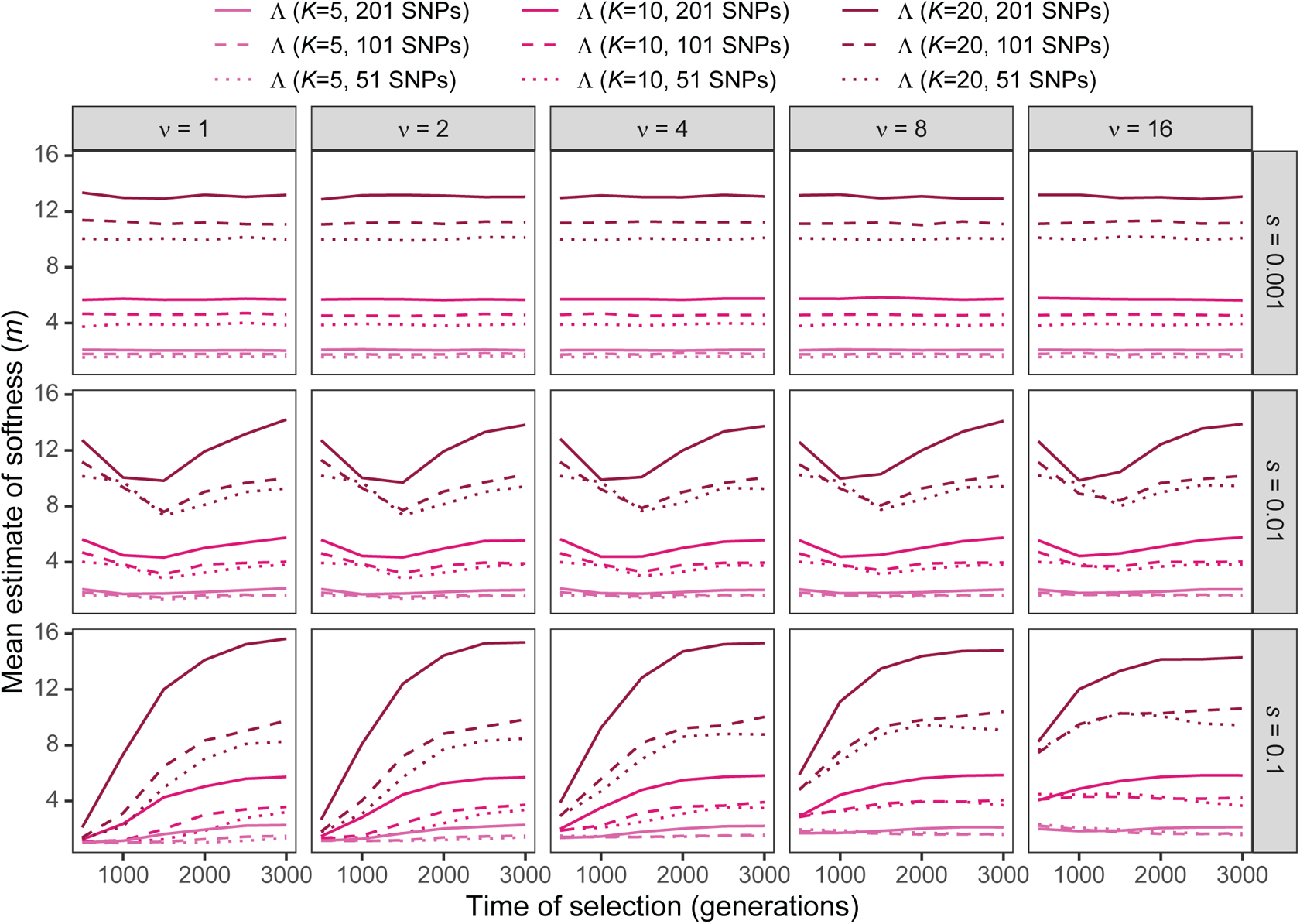
Estimated sweep softness illustrated by mean estimated number of sweeping haplotypes (*m*) in Λ with windows of size 51, 101, and 201 SNPs under simulations of a constant-size demographic history and the haplotype frequency spectra for the Λ statistic truncated at *K* 5, 10, 20 haplotypes for per-generation selection coefficients of *s* 0.001, 0.01, 0.1 on the rows. Mean estimated softness demonstrated for selection start times of *t* ∈ {500, 1000, 1500, 2000, 2500, 3000} generations prior to sampling for *ν ∈* {1, 2, 4, 8, 16} initially-selected haplotypes (columns). Results are based on a sample of *n* = 50 diploid individuals.

**Figure S34:**
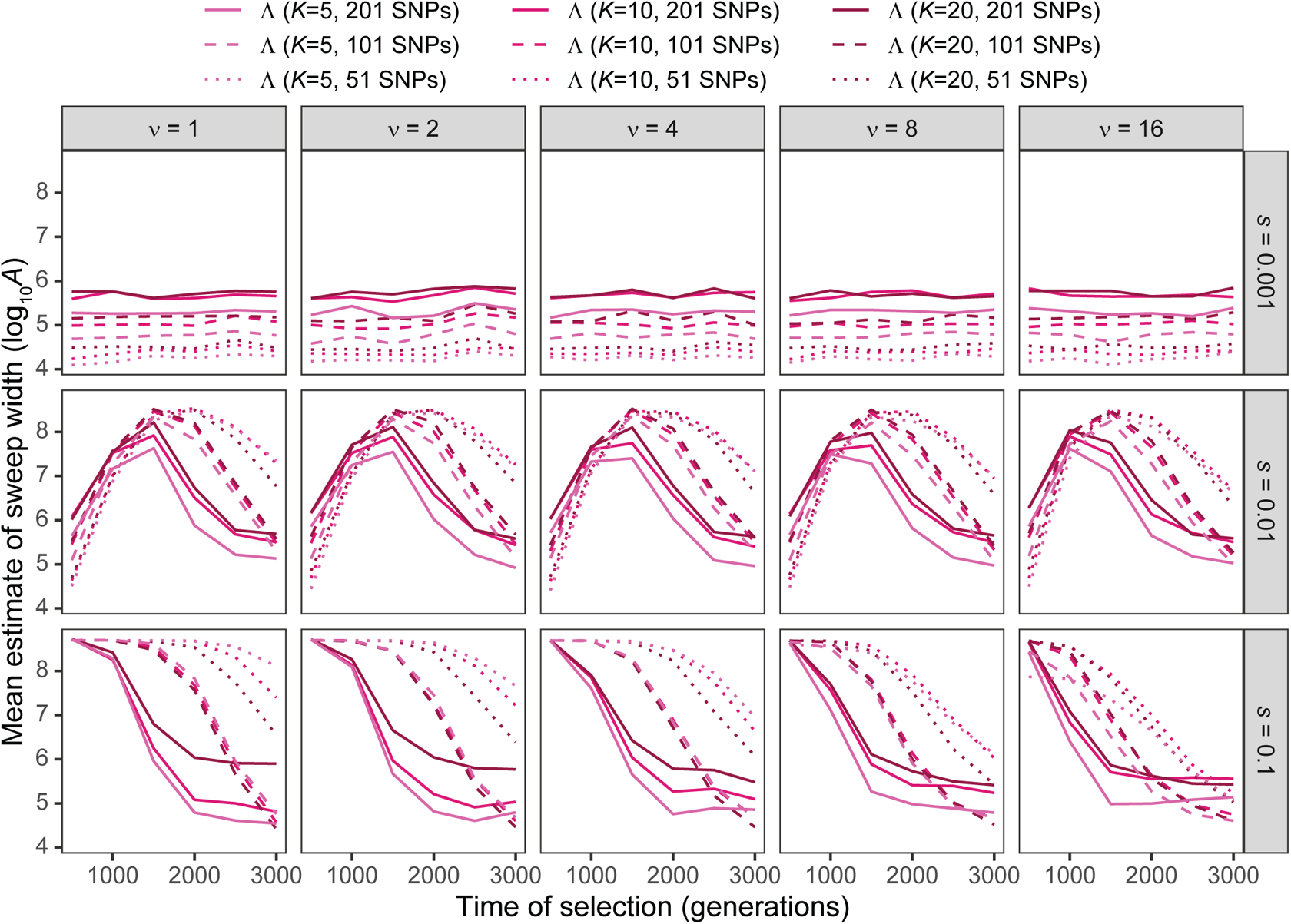
Estimated sweep width illustrated by mean estimated genomic size influenced by the sweep (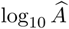) in Λ with windows of size 51, 101, and 201 SNPs under simulations of a constant-size demographic history and the haplotype frequency spectra for the Λ statistic truncated at *K* 5, 10, 20 haplotypes for per-generation selection coefficients of *s* 0.001, 0.01, 0.1 on the rows. Mean estimated genomic size influenced by sweeps demonstrated for selection start times of *t* ∈ {500, 1000, 1500, 2000, 2500, 3000} generations prior to sampling for *ν ∈* {1, 2, 4, 8, 16} initially-selected haplotypes (columns). Results are based on a sample of *n* = 50 diploid individuals.

**Figure S35:**
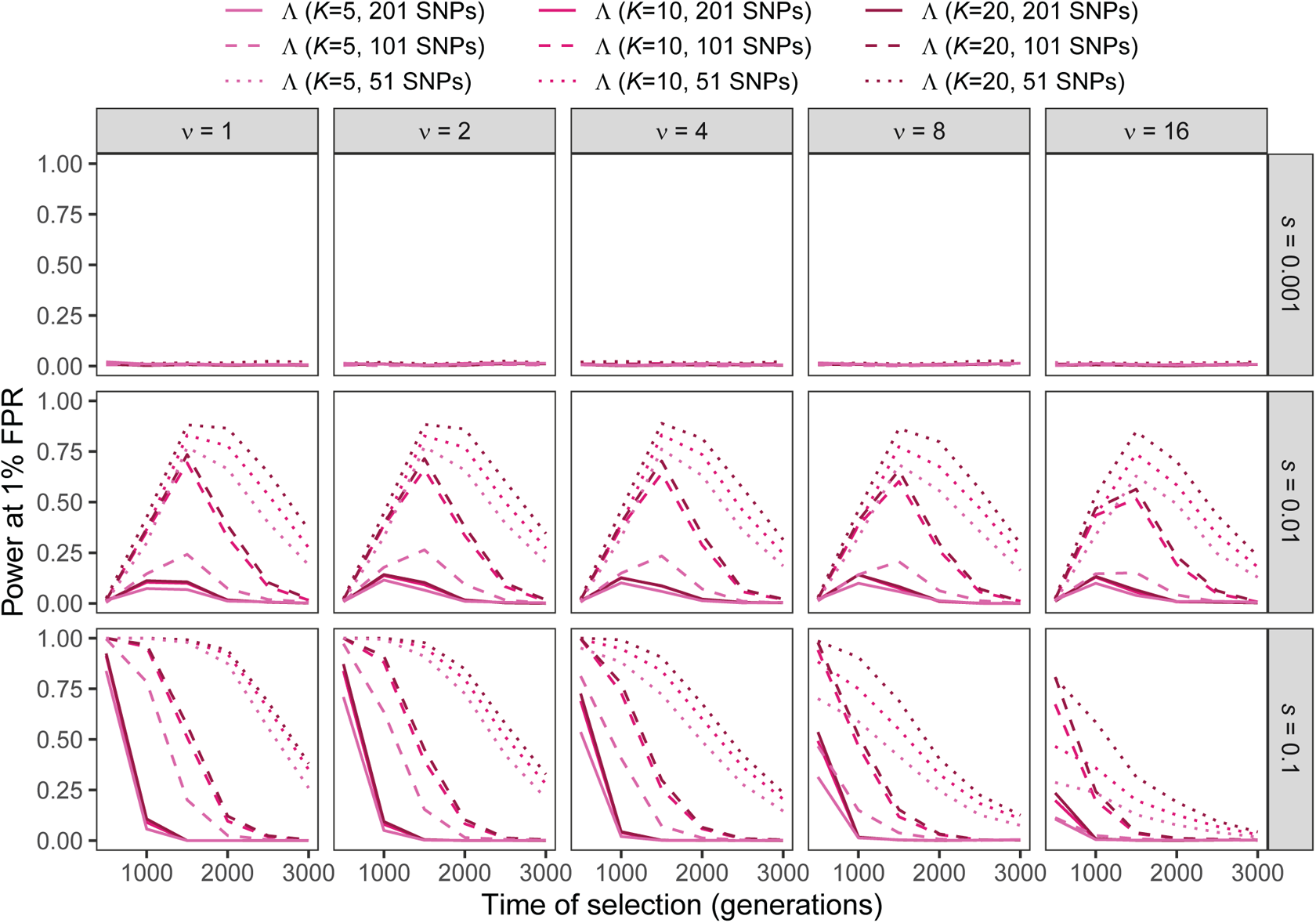
Power at a 1% false positive rate (FPR) as a function of selection start time for applications of Λ with windows of size 51, 101, and 201 SNPs to unphased multilocus genotype input data under simulations of a constant-size demographic history and the multilocus genotype frequency spectra for the Λ statistic truncated at *K* 5, 10, 20 multilocus genotypes for per-generation selection coefficients of *s* 0.001, 0.01, 0.1 on the rows. Classification ability demonstrated for selection start times of *t* ∈ {500, 1000, 1500, 2000, 2500, 3000} generations prior to sampling for *ν ∈* {1, 2, 4, 8, 16} initially-selected haplotypes (columns). Results are based on a sample of *n* = 50 diploid individuals.

**Figure S36:**
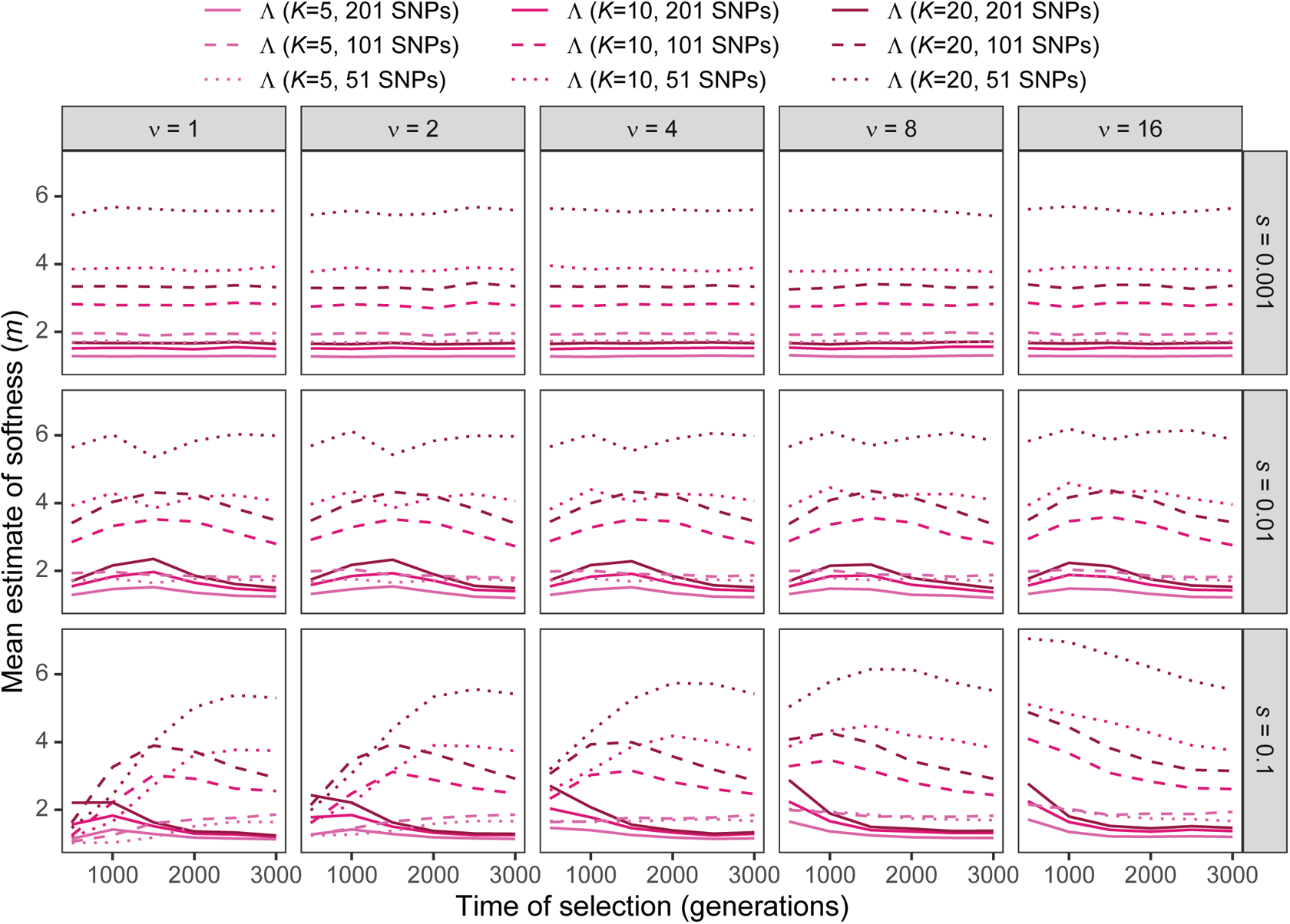
Estimated sweep softness illustrated by mean estimated number of sweeping haplotypes (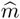) in Λ with windows of size 51, 101, and 201 SNPs applied to unphased multilocus input data under simulations of a constant-size demographic history and the multilocus genotype frequency spectra for the Λ statistic truncated at *K* 5, 10, 20 multilocus genotypes for per-generation selection coefficients of *s* 0.001, 0.01, 0.1 on the rows. Mean estimated softness demonstrated for selection start times of *t* ∈ {500, 1000, 1500, 2000, 2500, 3000} generations prior to sampling for *ν ∈* {1, 2, 4, 8, 16} initially-selected haplotypes (columns). Results are based on a sample of *n* = 50 diploid individuals.

**Figure S37:**
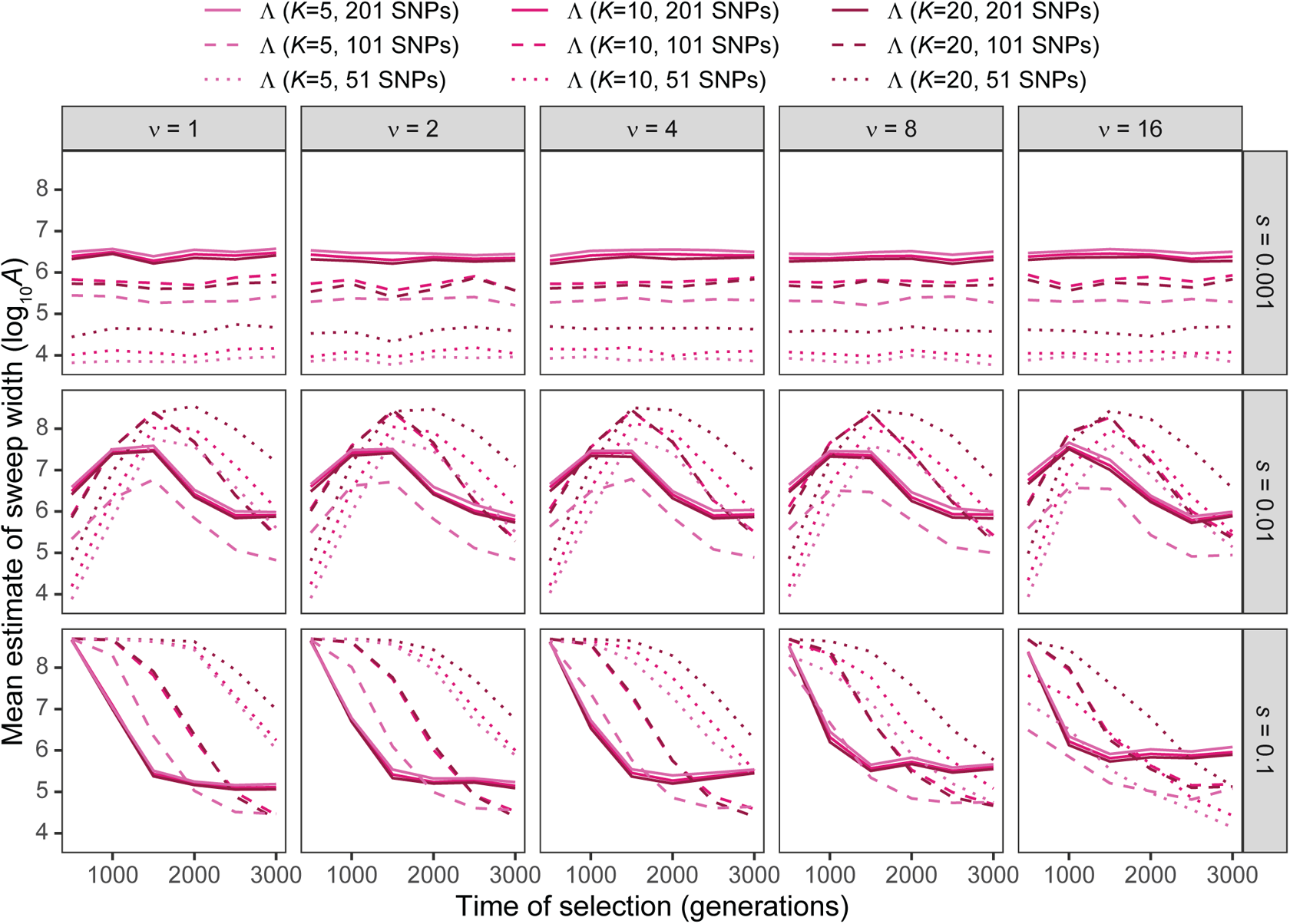
Estimated sweep width illustrated by mean estimated genomic size influenced by the sweep (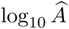) in Λ with windows of size 51, 101, and 201 SNPs applied to unphased multilocus input data under simulations of a constant-size demographic history and the multilocus genotype frequency spectra for the Λ statistic truncated at *K* 5, 10, 20 multilocus genotypes for per-generation selection coefficients of *s* 0.001, 0.01, 0.1 on the rows. Mean estimated genomic size influenced by sweeps demonstrated for selection start times of *t* ∈ {500, 1000, 1500, 2000, 2500, 3000} generations prior to sampling for *ν ∈* {1, 2, 4, 8, 16} initially-selected haplotypes (columns). Results are based on a sample of *n* = 50 diploid individuals.

**Figure S38:**
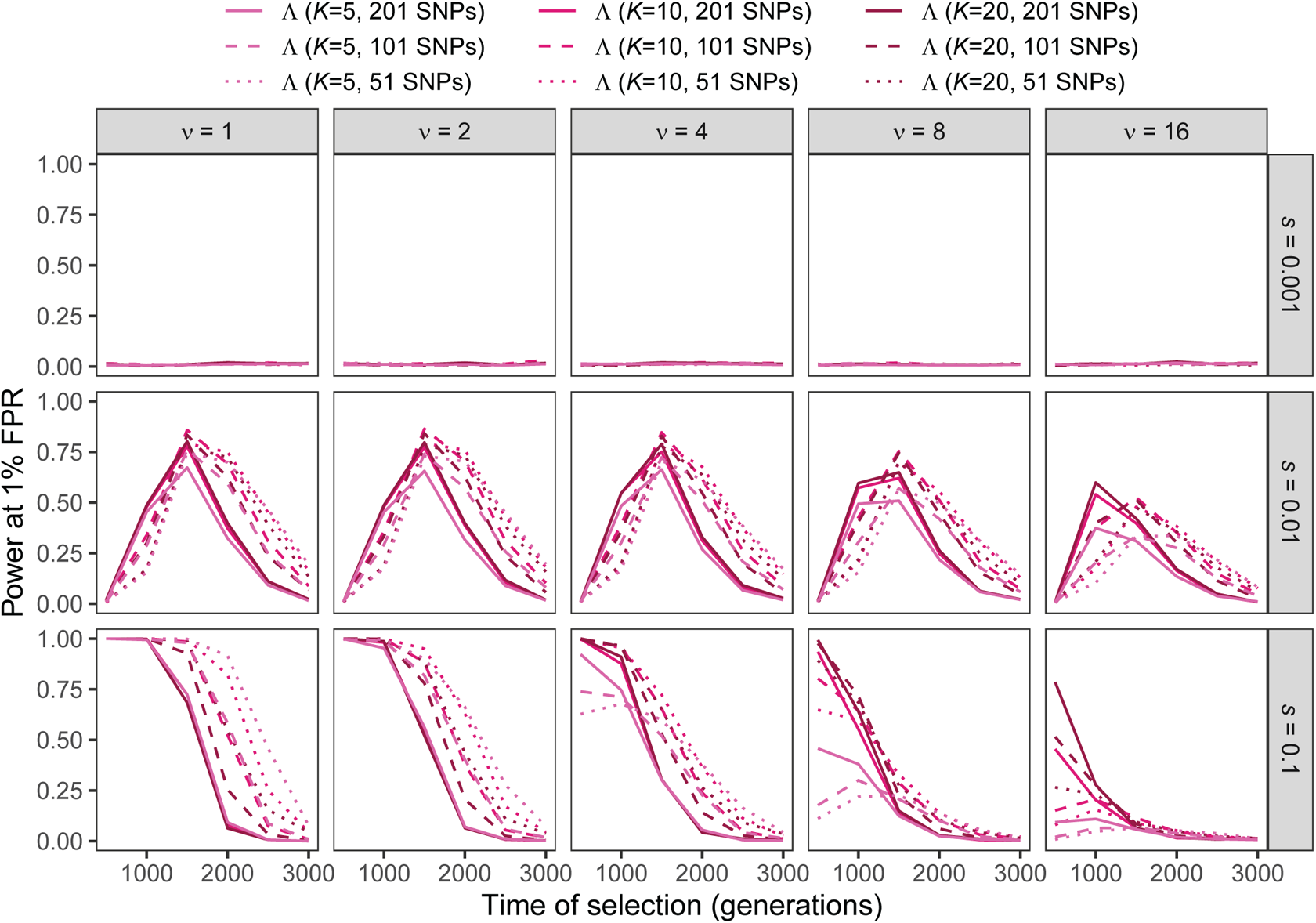
Power at a 1% false positive rate (FPR) as a function of selection start time for applications of Λ with windows of size 51, 101, and 201 SNPs under simulations of the human central European (CEU) demographic history of Terhorst et al. [2017] and the haplotype frequency spectra for the Λ statistic truncated at *K* 5, 10, 20 haplotypes for per-generation selection coefficients of *s* 0.001, 0.01, 0.1 on the rows. Classification ability demonstrated for selection start times of *t* ∈ {500, 1000, 1500, 2000, 2500, 3000} generations prior to sampling for *ν ∈* {1, 2, 4, 8, 16} initially-selected haplotypes (columns). Results are based on a sample of *n* = 50 diploid individuals.

**Figure S39:**
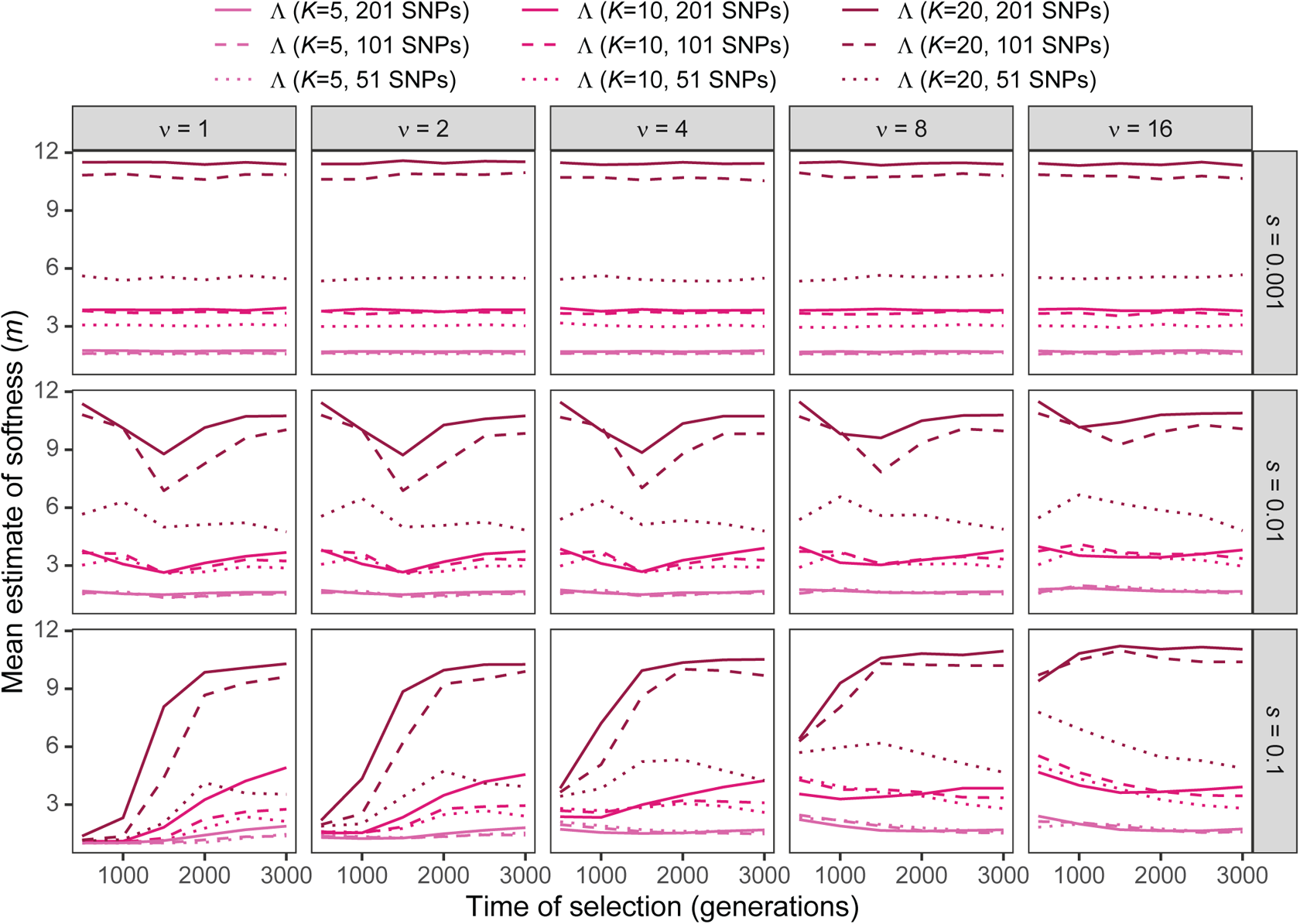
Estimated sweep softness illustrated by mean estimated number of sweeping haplotypes (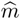) in Λ with windows of size 51, 101, and 201 SNPs under simulations of the human central European (CEU) demographic history of Terhorst et al. [2017] and the haplotype frequency spectra for the Λ statistic truncated at *K* 5, 10, 20 haplotypes for per-generation selection coefficients of *s* 0.001, 0.01, 0.1 on the rows. Mean estimated softness demonstrated for selection start times of *t* ∈ {500, 1000, 1500, 2000, 2500, 3000} generations prior to sampling for *ν ∈* {1, 2, 4, 8, 16} initially-selected haplotypes (columns). Results are based on a sample of *n* = 50 diploid individuals.

**Figure S40:**
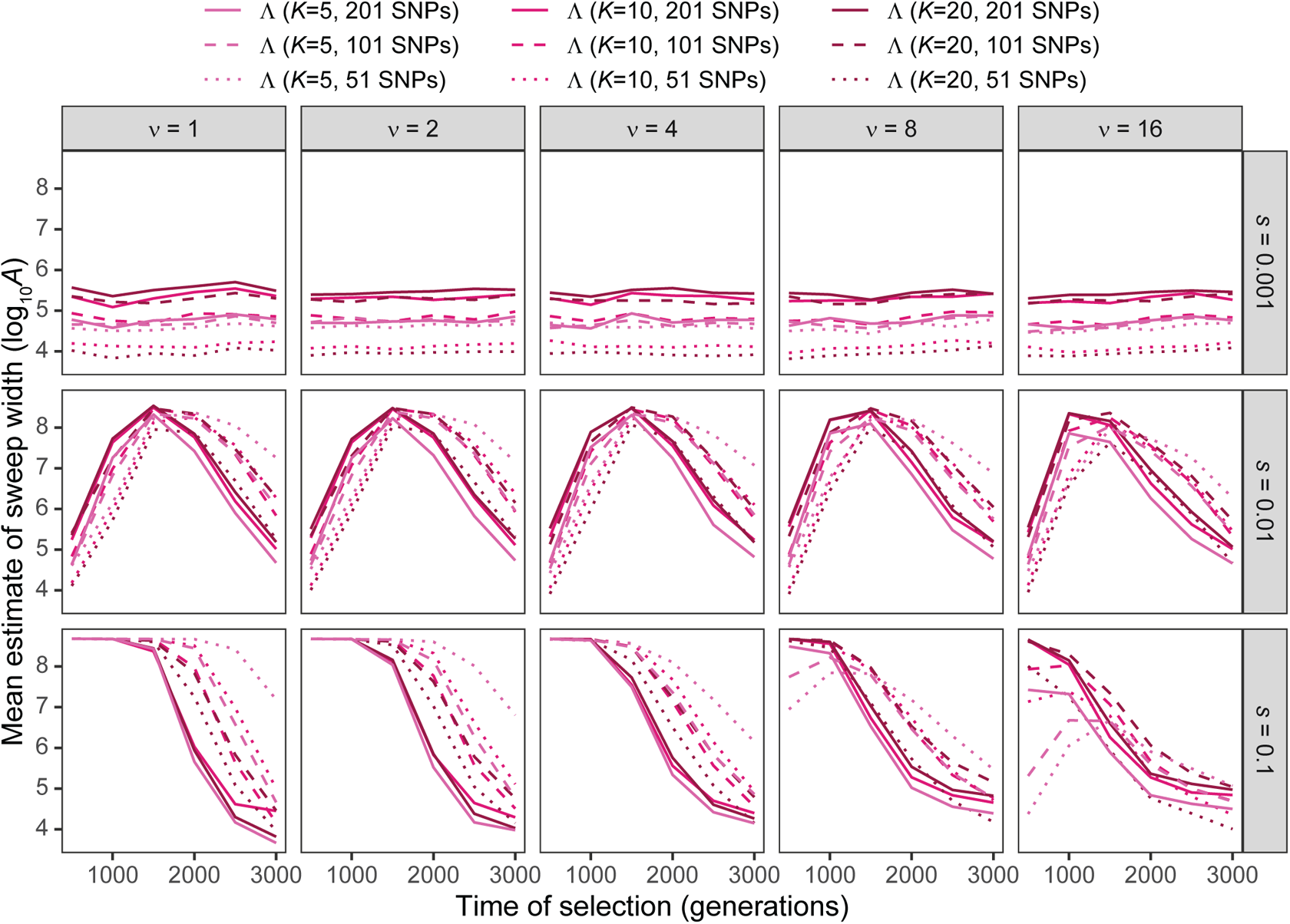
Estimated sweep width illustrated by mean estimated genomic size influenced by the sweep (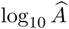) in Λ with windows of size 51, 101, and 201 SNPs under simulations of the human central European (CEU) demographic history of Terhorst et al. [2017] and the haplotype frequency spectra for the Λ statistic truncated at *K* 5, 10, 20 haplotypes for per-generation selection coefficients of *s* 0.001, 0.01, 0.1 on the rows. Mean estimated genomic size influenced by sweeps demonstrated for selection start times of *t* ∈ {500, 1000, 1500, 2000, 2500, 3000} generations prior to sampling for *ν ∈* {1, 2, 4, 8, 16} initially-selected haplotypes (columns). Results are based on a sample of *n* = 50 diploid individuals.

**Figure S41:**
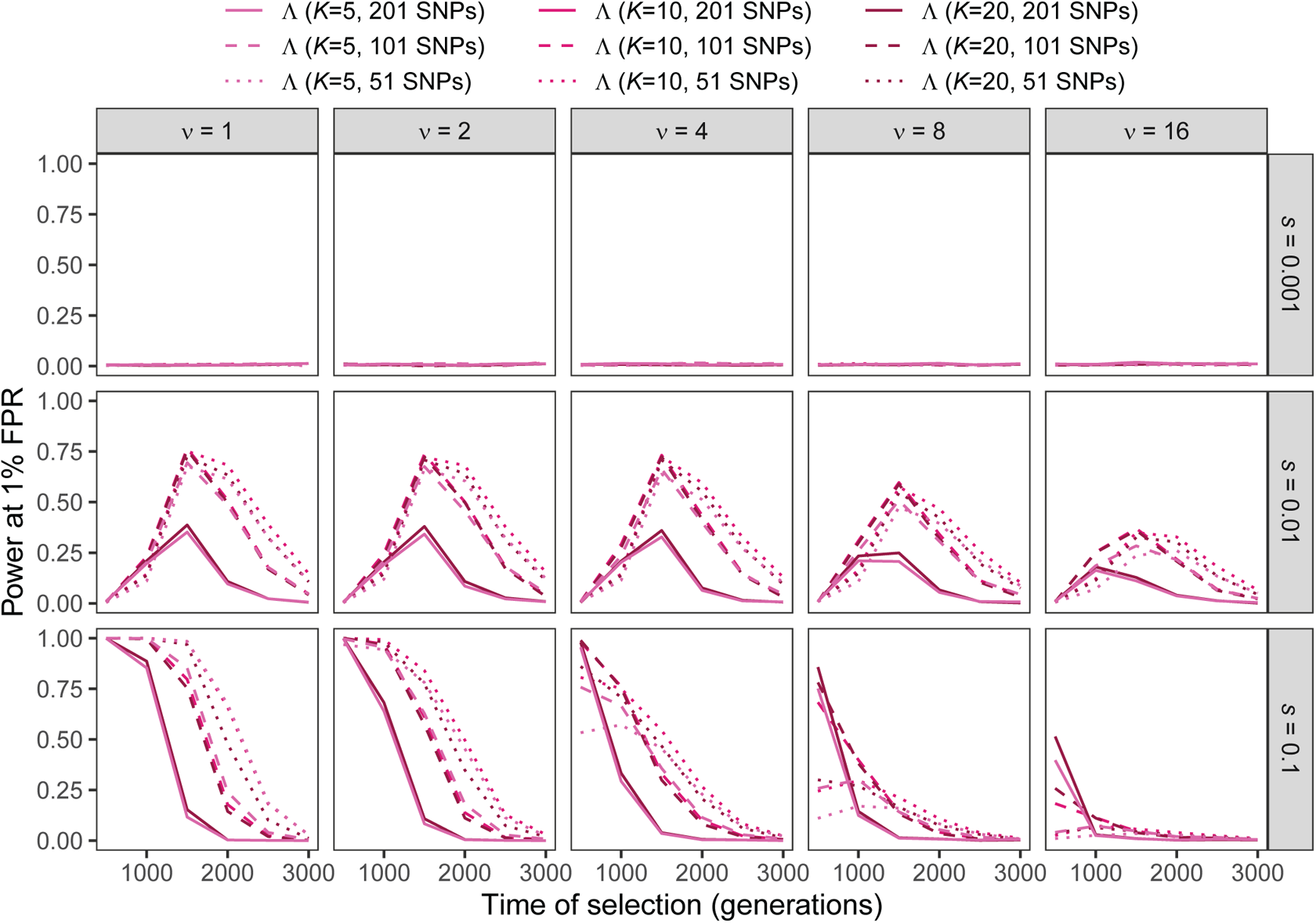
Power at a 1% false positive rate (FPR) as a function of selection start time for applications of Λ with windows of size 51, 101, and 201 SNPs to unphased multilocus genotype input data under simulations of the human central European (CEU) demographic history of Terhorst et al. [2017] and the multilocus genotype frequency spectra for the Λ statistic truncated at *K* 5, 10, 20 multilocus genotypes for per-generation selection coefficients of *s* 0.001, 0.01, 0.1 on the rows. Classification ability demonstrated for selection start times of *t* ∈ {500, 1000, 1500, 2000, 2500, 3000} generations prior to sampling for *ν ∈* {1, 2, 4, 8, 16} initially-selected haplotypes (columns). Results are based on a sample of *n* = 50 diploid individuals.

**Figure S42:**
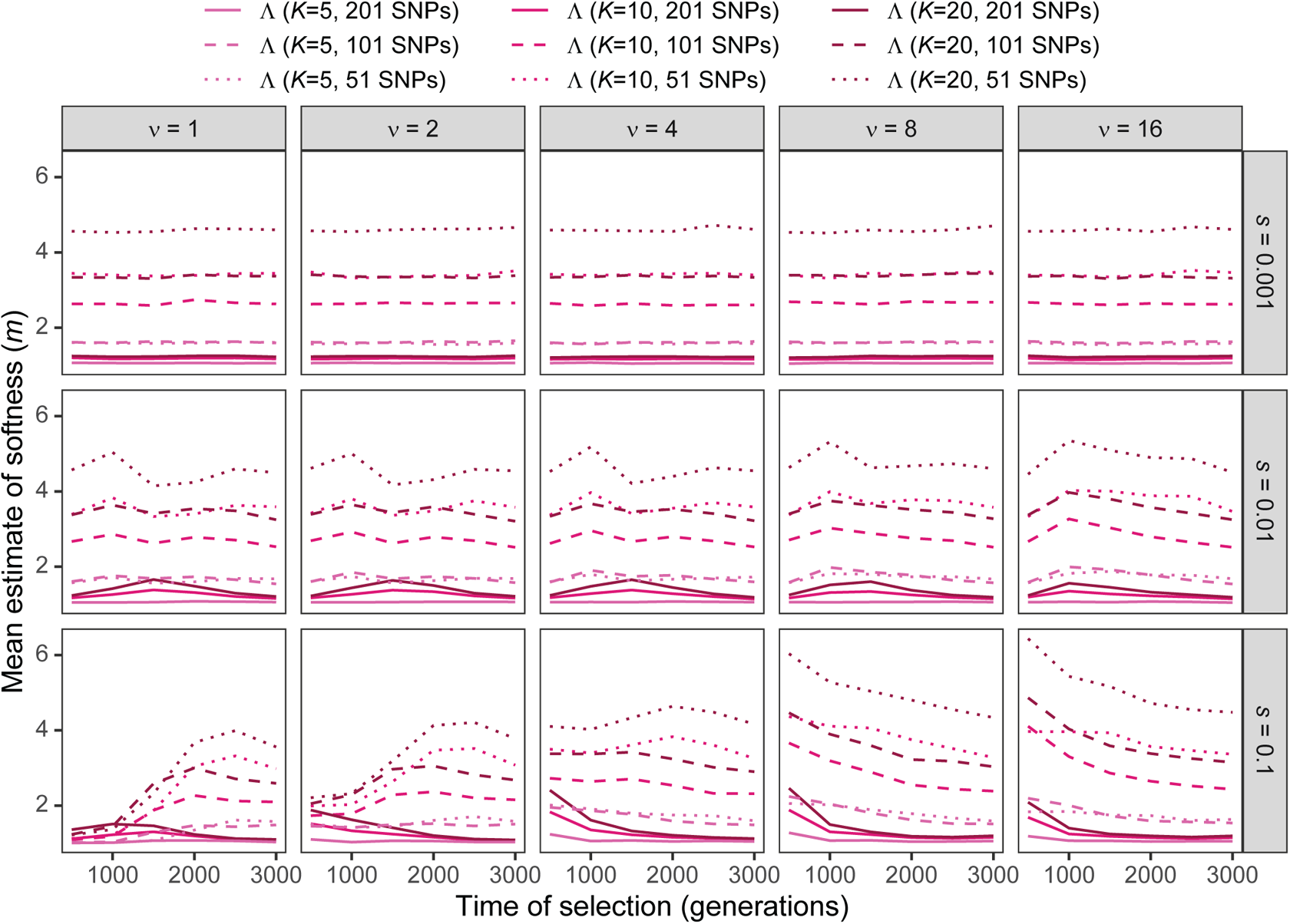
Estimated sweep softness illustrated by mean estimated number of sweeping haplotypes (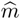) in Λ with windows of size 51, 101, and 201 SNPs applied to unphased multilocus input data under simulations of the human central European (CEU) demographic history of Terhorst et al. [2017] and the multilocus genotype frequency spectra for the Λ statistic truncated at *K* 5, 10, 20 multilocus genotypes for per-generation selection coefficients of *s* 0.001, 0.01, 0.1 on the rows. Mean estimated softness demonstrated for selection start times of *t* ∈ {500, 1000, 1500, 2000, 2500, 3000} generations prior to sampling for *ν ∈* {1, 2, 4, 8, 16} initially-selected haplotypes (columns). Results are based on a sample of *n* = 50 diploid individuals.

**Figure S43:**
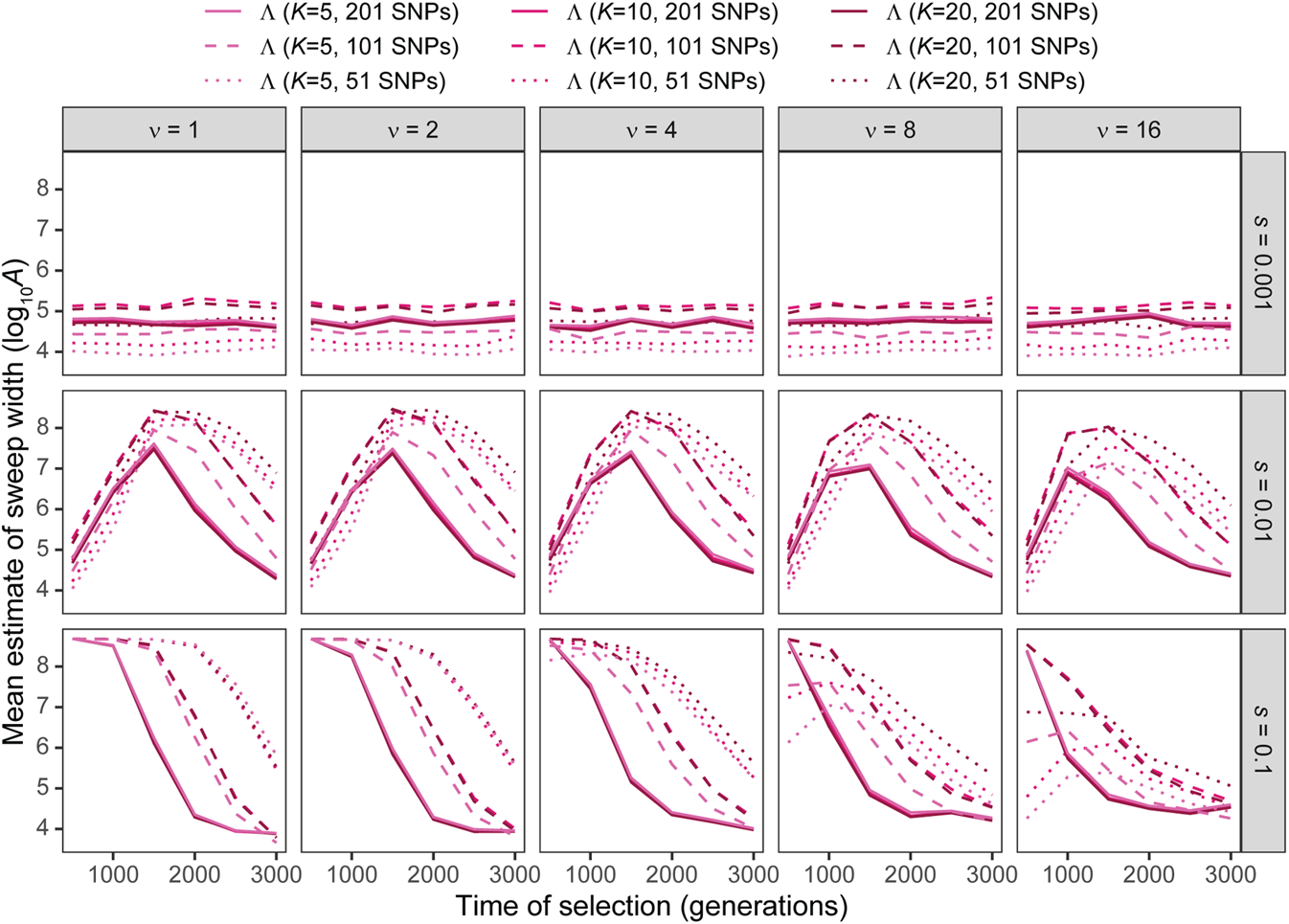
Estimated sweep width illustrated by mean estimated genomic size influenced by the sweep (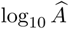) in Λ with windows of size 51, 101, and 201 SNPs applied to unphased multilocus input data under simulations of the human central European (CEU) demographic history of Terhorst et al. [2017] and the multilocus genotype frequency spectra for the Λ statistic truncated at *K* 5, 10, 20 multilocus genotypes for per-generation selection coefficients of *s* 0.001, 0.01, 0.1 on the rows. Mean estimated genomic size influenced by sweeps demonstrated for selection start times of *t* ∈ {500, 1000, 1500, 2000, 2500, 3000} generations prior to sampling for *ν ∈* {1, 2, 4, 8, 16} initially-selected haplotypes (columns). Results are based on a sample of *n* = 50 diploid individuals.

**Figure S44:**
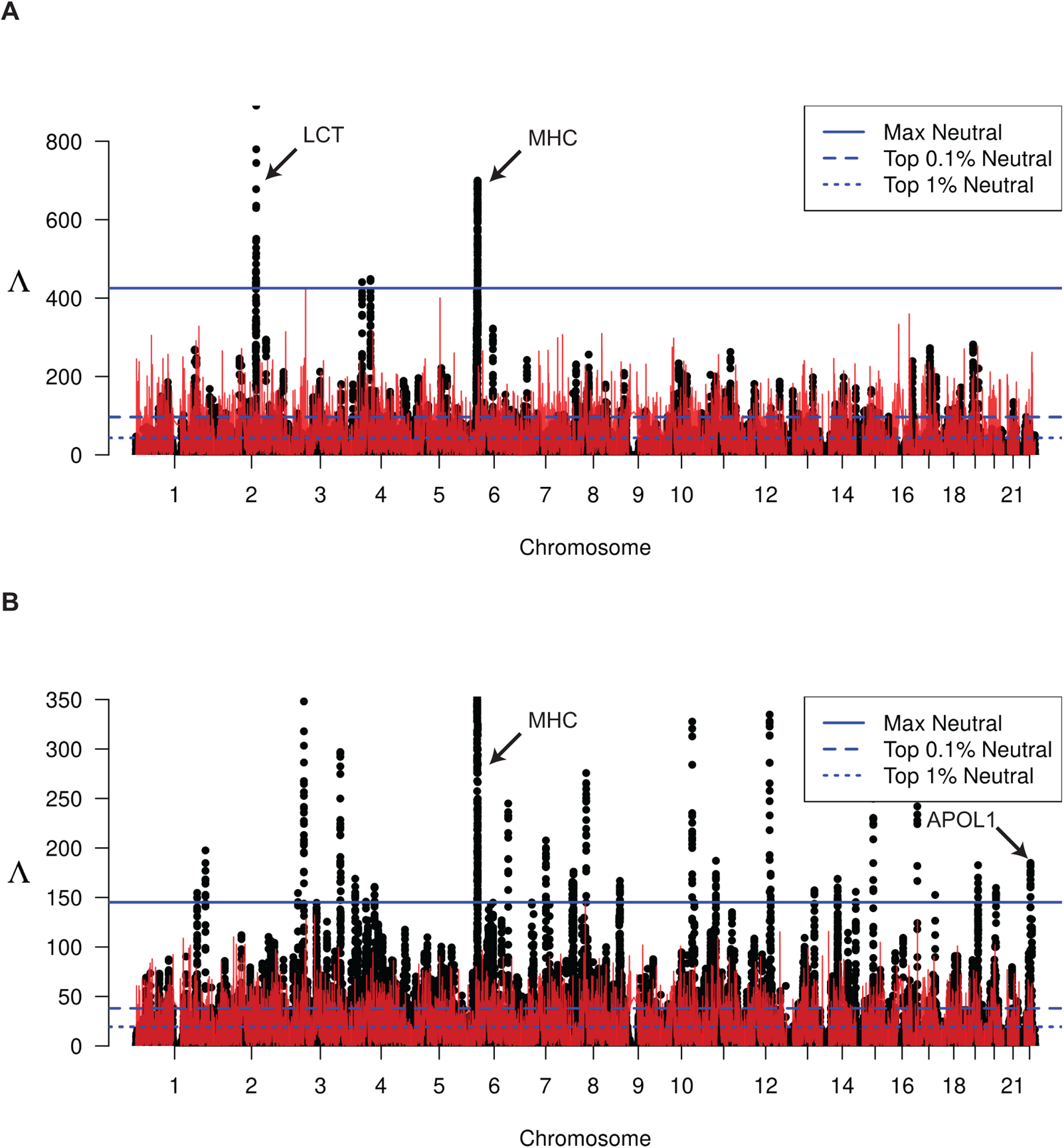
Manhattan plot of unphased multi-locus genotype Λ-statistics for the (A) CEU and (B) YRI populations from the 1000 Genomes Project. Each point represents a single 201-SNP window along the genome. Horizontal lines represent the top 1%, top 0.1%, and maximum observed Λ statistic across all windows in demography-matched neutral simulations. Red line indicates the maximum observed Λ among 100 replicate simulations at that location in the genome.

**Figure S45:**
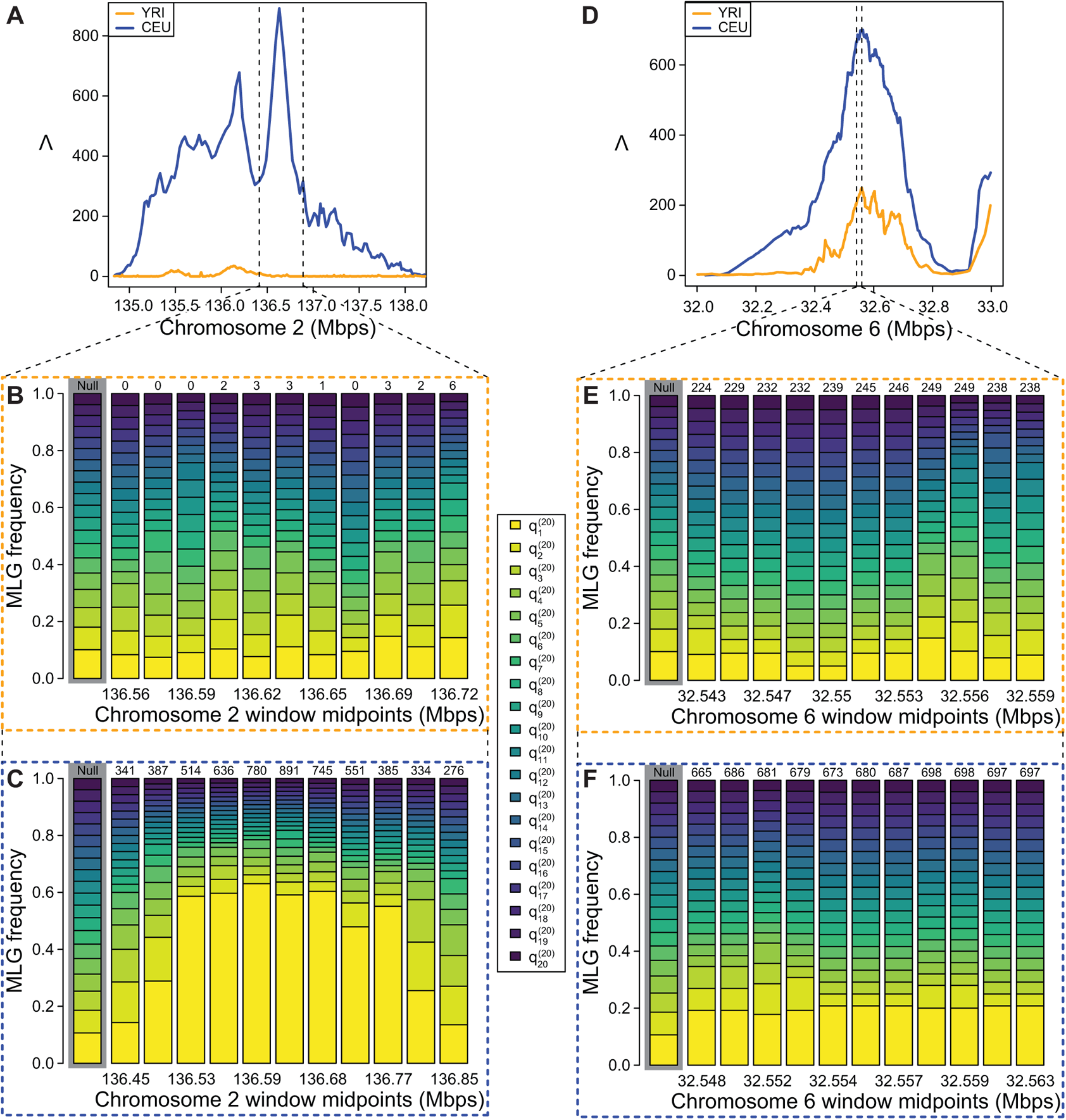
Detailed illustration of Λ statistics and multi-locus genotype frequency spectra in CEU and YRI. (A) Λ plotted in the *LCT* region, vertical dotted lines indicate zoomed region shown in (A) and (C). (B) YRI empirical HFS for 11 windows in the *LCT* region. (C) CEU empirical HFS for 11 windows in the *LCT* region. (D) Λ plotted in the MHC region, vertical dotted lines indicate zoomed region shown in (E) and (F). (E) YRI empirical HFS for 11 windows in the MHC region. (F) CEU empirical HFS for 11 windows in the MHC region. In (B), (C), (E), and (F), numbers above HFS are Λ values for the window rounded to the nearest whole number, and the genome-wide average HFS is highlighted in grey. 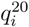 is the frequency of the *i*th most common MLG truncated to *K* = 20.

**Figure S46:**
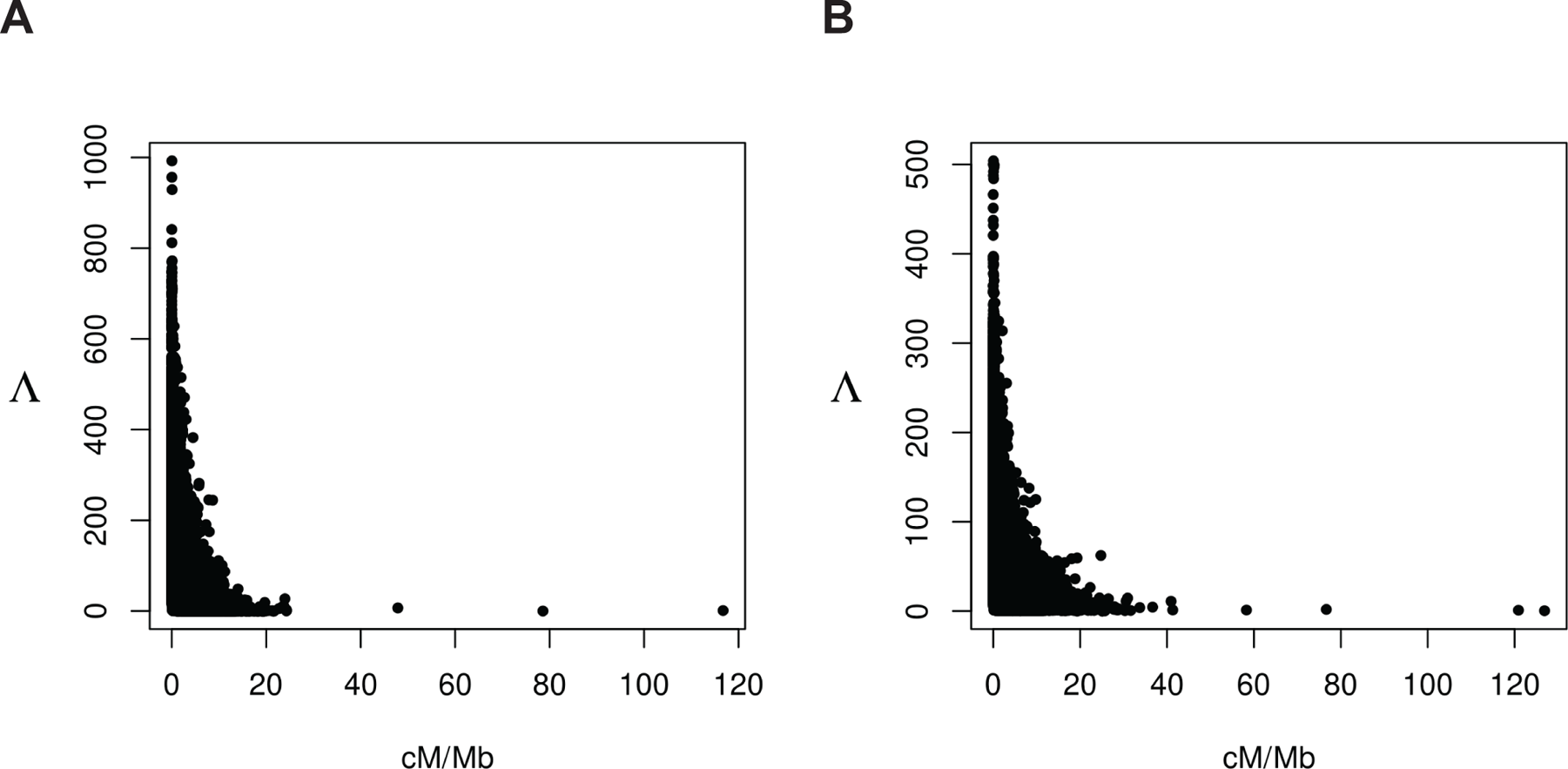
Maximum Λ observed per window across demography-matched neutral simulations versus recombination rate for the (A) CEU and (B) YRI populations.

## Notes

### Competing Interest Statement

The authors have declared no competing interest.

## References

1. JR Adrion, CB Cole, N Dukler, JG Galloway, AL Gladstein, G Gower, CC Kyriazis, AP Ragsdale, G Tsambos, F Baumdicker, J Carlson, RA Cartwright, A Durvasula, I Gronau, BY Kim, P McKenzie, PW Messer, E Noskova, D Ortega-Del Vecchyo, F Racimo, TJ Struck, S Gravel, RN Gutenkunst, KE Lohmueller, PL Ralph, DR Schrider, A Siepel, J Kelleher, and AD Kern. A community-maintained standard library of population genetic models. eLife, 9:e54967, 2020.

2. H Akashi, N Osada, and T Ohta. Weak selection and protein evolution. Genetics, 192:15–31, 2012.

3. A Albrechtsen, I Moltke, and R Nielsen. Natural selection and the distribution of identity-by-descent in the human genome. Genetics, 186:295–308, 2010.

4. N Barton. The effect of hitch-hiking on neutral genealogies. Genet Res, 72:123–133, 1998.

5. AC Beichman, TN Phung, and KE Lohmueller. Comparison of single genome and allele frequency data reveals discordant demographic histories. G3 (Bethesda), 7:3605–3620, 2017.

6. T Bersaglieri, PC Sabeti, N Patterson, T Vanderploeg, T Schaffner, JA Drake, M Rhodes, DE Reich, and JN Hirschhorn. Genetic signatures of strong recent positive selection at the lactase gene. Am J Hum Genet, 74:1111–1120, 2004.

7. AR Boyko, SH Williamson, AR Indap, JD Degenhardt, RD Hernandez, KE Lohmueller, MD Adams, S Schmidt, JJ Sninsky, SR Sunyaev, TJ White, R Nielsen, AG Clark, and CD Bustamante. Assessing the evolutionary impact of amino acid mutations in the human genome. PLoS Genet, 30:e1000083, 2008.

8. B Charlesworth. The role of background selection in shaping patterns of molecular evolution and variation: evidence from variability on the *Drosophila* X chromosome. Genetics, 191:233–2463, 2012.

9. B Charlesworth, MT Morgan, and D Charlesworth. The effect of deleterious mutations on neutral molecular variation. Genetics, 134:1289–1303, 1993.

10. D Charlesworth, B Charlesworth, and MT Morgan. The pattern of neutral molecular variation under the background selection model. Genetics, 141:1619–1632, 1995.

11. H Chen, N Patterson, and D Reich. Population differentiation as a test for selective sweeps. Genome Res, 20:393–402, 2010.

12. X Cheng and M DeGiorgio. Detection of shared balancing selection in the absence of trans-species polymorphism. Mol Biol Evol, 36:177–199, 2019.

13. X Cheng and M DeGiorgio. Flexible mixture model approaches that accommodate footprint size variability for robust detection of balancing selection. Mol Biol Evol, 37:3267–3291, 2020.

14. JM Comeron. Background selection as a baseline for nucleotide variation across the *Drosophila* genome. PLoS Genet, 10:e1004434, 2014.

15. The International HapMap Consortium. A second generation human haplotype map of over 3.1 million SNPs. Nature, 449:841, 2007.

16. M DeGiorgio, KE Lohmueller, and R Nielsen. A model-based approach for identifying signatures of ancient balancing selection in genetic data. PLoS Genet, 10:e1004561, 2014.

17. M DeGiorgio, CD Huber, MJ Hubisz, I Hellmann, and R Nielsen. *SweepFinder2* : Increased sensitivity, robustness, and flexibility. Bioinformatics, 32:1895–1897, 2016.

18. T Derrien, J Estellé, S Marco Sola, DG Knowles, E Raineri, R Guigó, and P Ribeca. Fast computation and applications of genome mappability. PLoS One, 7:e30377, 2012.

19. A Dilthey, C Cox, Z Iqbal, MR Nelson, and G McVean. Improved genome inference in the MHC using a population reference graph. Nat Genet, 47:682–688, 2015.

20. R Durrett and J Schweinsberg. Approximating selective sweeps. Theor Popul Biol, 66:129–138, 2004.

21. D Enard, PW Messer, and DA Petrov. Genome-wide signals of positive selection in human evolution. Genome Res, 24:884–895, 2014.

22. M Fagny, E Patin, D Enard, L Quintana-Murci, and G Laval. Exploring the occurrence of classic selective sweeps in humans using whole-genome sequencing data sets. Mol Biol Evol, 31:1850–1868, 2014.

23. A Ferrer-Admetlla, M Liang, T Korneliussen, and R Nielsen. On detecting incomplete soft or hard selective sweeps using haplotype structure. Mol Biol Evol, 31:1275–1291, 2014.

24. Y Field, EA Boyle, N Telis, Z Gao, KJ Gaulton, D Golan, L Yengo, G Rocheleau, P Froguel, MI McCarthy, and JK Pritchard. Detection of human adaptation during the past 2000 years. Science, 354:760–764, 2016.

25. NR Garud, PW Messer, EO Buzbas, and DA Petrov. Recent selective sweeps in North American *Drosophila melanogaster* show signatures of soft sweeps. PLoS Genet, 11:e1005004, 2015.

26. T Goeury, LE Creary, L Brunet, M Galan, M Pasquier, B Kervaire, A Langaney, J-M Tiercey, MA Fernández-Viña, JM Nunes, and A Sachez-Mazas. Deciphering the fine nucleotide diversity of full HLA class I and class II genes in a well-documented population from sub-Saharan Africa. HLA, 91: 36–51, 2018.

27. S Gravel, BM Henn, RN Gutenkunst, AR Indap, GT Marth, AG Clark, F Yu, R Gibbs, The 1000 Genomes Project, and D Bustamante. Demographic history and rare allele sharing among human populations. Proc Natl Acad Sci USA, 108:11983–11988, 2011.

28. I Gronau, MJ Hubisz, B Gulko, CG Danko, and A Siepel. Bayesian inference of ancient human demography from individuals genomes. Nat Genet, 43:1031–1034, 2011.

29. BC Haller and PW Messer. SLiM 3: Forward genetic simulations beyond the Wright-Fisher model. Mol Biol Evol, 36:632–637, 2019.

30. A Harpak, N Garud, NA Rosenberg, DA Petrov, M Combs, PS Pennings, and J Munshi-South. Genetic adaptation in New York City rats. Genome Biol Evol, 13:evaa247, 2021.

31. AM Harris and M DeGiorgio. A likelihood approach for uncovering selective sweep signatures from haplotype data. Mol Biol Evol, 37:3023–3046, 2020.

32. AM Harris, NR Garud, and M DeGiorgio. Detection and classification of hard and soft sweeps from unphased genotypes by multilocus genotype identity. Genetics, 210:1429–1452, 2018.

33. T Hastie, R Tibshirani, and J Friedman. The elements of statistical learning: data mining, inference, and prediction. Springer, New York, NY, 2nd edition, 2009.

34. J Hermisson and PS Pennings. Soft sweeps. Genetics, 4:2335–2352, 2005.

35. CD Huber, M DeGiorgio, I Hellmann, and R Nielsen. Detecting recent selective sweeps while controlling for mutation rate and background selection. Mol Ecol, 25:142–156, 2015.

36. RR Hudson and NL Kaplan. Deleterious background selection with recombination. Genetics, 141:1605–1617, 1995a.

37. RR Hudson and NL Kaplan. The coalescent process and background selection. Philos Trans R Soc B, 349: 19–23, 1995b.

38. JD Jensen, Y Kim, V Bauer DuMont, CF Aquadro, and CD Bustamante. Distinguishing between selective sweeps and demography using DNA polymorphism data. Genetics, 170:1401–1410, 2005.

39. J Kelleher, AM Etheridge, and G McVean. Efficient coalescent simulation and genealogical analysis for large sample sizes. PLoS Comput Biol, 12:1–12, 2016.

40. AD Kern and DR Schrider. diploS/HIC: an updated approach to classifying selective sweeps. G3 (Bethesda), 8:1959–1970, 2018.

41. Y Kim and W Stephan. Detecting a local signature of genetic hitchhiking along a recombining chromosome. Genetics, 160:765–777, 2002.

42. W-Y Ko, P Rajan, F Gomez, L Scheinfeldt, P An, CA Winkler, A Froment, TB Nyambo, SA Omar, C Wambebe, A Ranciaro, JB Hirbo, and SA Tishkoff. Identifying Darwinian selection acting on different human APOL1 variants among diverse African populations. Am J Hum Genet, 93:54–66, 2013.

43. KM Lee and G Coop. Distinguishing among modes of convergent adaptation using population genomic data. Genetics, 207:1591–1619, 2017.

44. RY Lieu and K Singh. Moving blocks jacknife and bootstrap capture weak dependence, pp. 225–248 in Exploring the “Limits” of the Boostrap. John Wiley and Sons, New York, 1992.

45. K Lin, H Li, C Schlötterer, and A Futschik. Distinguishing positive selection from neutral evolution: boosting the performance of summary statistics. Genetics, 187:229–244, 2011.

46. KE Lohmueller, A Albrechtsen, Y Li, SY Kim, T Koneliussen, N Vinckenbosch, G Tian, E Huerta-Sanchez, AF Feder, N Garup, T Jørgensen, T Jiang, DR Witte, A Sandbæk, I Hellmann, T Lauritzen, T Hansen, O Pedersen, J Wang, and R Nielsen. Natural selection affects multiple aspects of genetic variation at putatively neutral sites across the human genome. PLoS Genet, 7:e1002326, 2011.

47. S Mallick, S Gnerre, and D Reich. The difficulty of avoiding false positives in genome scans for natural selection. Genome Res, 19:922–933, 2009.

48. GA McVean and B Charlesworth. The effects of Hill-Robertson interference between weakly selected mutations on patterns of molecular evolution and variation. Genetics, 155:929–944, 2000.

49. G McVicker, D Gordon, C Davis, and P Green. Widespread genomic signatures of natural selection in hominid evolution. PLoS Genet, 5:e1000471, 2009.

50. Huaiyu Mi, Dustin Ebert, Anushya Muruganujan, Caitlin Mills, Laurent-Philippe Albou, Tremayne Mushayamaha, and Paul D Thomas. PANTHER version 16: a revised family classification, tree-based classification tool, enhancer regions and extensive API. Nucleic Acids Research, 49(D1):D394–D403, 12 2020. ISSN 0305-1048. doi: 10.1093/nar/gkaa1106. URL https://doi.org/10.1093/nar/gkaa1106.

51. MR Mughal and M DeGiorgio. Localizing and classifying selective sweeps with trend filtered regression. Mol Biol Evol, 36:252–270, 2019.

52. MR Mughal, H Koch, J Huang, F Chiaromonte, and M DeGiorgio. Learning the properties of adaptive regions with functional data analysis. PLoS Genet, in press, 2020.

53. LE Nicolaisen and MM Desai. Distortions in genealogies due to purifying selection and recombination. Genetics, 194:221–230, 2013.

54. R Nielsen, Scott Williamson, Y Kim, MJ Hubisz, AG Clark, and C Bustamante. Genomic scans for selective sweeps using SNP data. Genome Res, 15:1566–1575, 2005.

55. M Nordborg, B Charlesworth, and D Charlesworth. The effect of recombination of background selection. Genet Res, 67:159–174, 1996.

56. Yohann Nédélec, Joaquín Sanz, Golshid Baharian, Zachary A. Szpiech, Alain Pacis, Anne Dumaine, Jean-Christophe Grenier, Andrew Freiman, Aaron J. Sams, Steven Hebert, Ariane Pagé Sabourin, Francesca Luca, Ran Blekhman, Ryan D. Hernandez, Roger Pique-Regi, Jenny Tung, Vania Yotova, and Luis B. Barreiro. Genetic ancestry and natural selection drive population differences in immune responses to pathogens. Cell, 167(3):657–669.e21, 2016. ISSN 0092-8674. doi: https://doi.org/10.1016/j.cell.2016.09.025. URL https://www.sciencedirect.com/science/article/pii/S0092867416313071.

57. Pierantonio Parmiani, Cristina Lucchetti, and Gianfranco Franchi. Whisker and nose tactile sense guide rat behavior in a skilled reaching task. Frontiers in behavioral neuroscience, 12:24, 2018.

58. Michael H Parsons, Raimund Apfelbach, Peter B Banks, Elissa Z Cameron, Chris R Dickman, Anke SK Frank, Menna E Jones, Ian S McGregor, Stuart McLean, Dietland Müller-Schwarze, et al. Biologically meaningful scents: a framework for understanding predator–prey research across disciplines. Biological Reviews, 93(1):98–114, 2018.

59. Michael H Parsons, Michael A Deutsch, Dani Dumitriu, and Jason Munshi-South. Differential responses by urban brown rats (Rattus norvegicus) toward male or female-produced scents in sheltered and high-risk presentations. Journal of Urban Ecology, 5(1), 09 2019. ISSN 2058-5543. doi: 10.1093/jue/juz009. URL https://doi.org/10.1093/jue/juz009. juz009.

60. P Pavlidis, S Hutter, and W Stephan. A population genomic approach to map recent positive selection in model species. Mol Ecol, 17:3585–2598, 2008.

61. BA Payseur and MW Nachman. Micorsatelllite variation and recombination rate in the human genome. Genetics, 156:1285–1298, 2000.

62. PS Pennings and J Hermisson. Soft sweeps II—molecular population genetics of adaptation from recurrent mutation or migration. Mol Biol Evol, 23:1076–1084, 2006a.

63. PS Pennings and J Hermisson. Soft sweeps III: the signature of positive selection from recurrent mutation. PLoS Genet, 2:1–15, 2006b.

64. F Pierini and TL Lenz. Divergent allele advantage at human MHC genes: Signatures of past and ongoing selection. Mol Biol Evol, 35:2145–2158, 2018.

65. Marko Piirsoo, Dies Meijer, and Tõnis Timmusk. Expression analysis of the clca gene family in mouse and human with emphasis on the nervous system. BMC developmental biology, 9(1):1–11, 2009.

66. M Przeworski. The signature of positive selection at randomly chosen loci. Genetics, 160:1179–1189, 2002.

67. F Racimo. Testing for ancient selection using cross-population allele frequency differentiation. Genetics, 202:733–750, 2016.

68. T Ruths and L Nakhleh. Boosting forward-time population genetic simulators through genotype compression. BMC Bioinformatics, 14, 2013. doi: 10.1186/1471-2105-14-192.

69. PC Sabeti, DE Reich, JM Higgins, HZP Levine, DJ Richter, SF Schaffner, SB Gabriel, JV Platko, NJ Patterson, GJ McDonald, HC Ackerman, SJ Campbell, D Altshuler, R Cooper, D Kwiatkowski, R Ward, and ES Lander. Detecting recent positive selection in the human genome from haplotype structure. Nature, 419:832–837, 2002.

70. PC Sabeti, P Varilly, B Fry, J Lohmueller, E Hostetter, C Cotsapas, X Xie, EH Byrne, SA McCarroll, R Gaudet, SF Schaffner, ES Lander, and The International HapMap Consortium. Genome-wide detection and characterization of positive selection in human populations. Nature, 449:913–918, 2007.

71. A Scally and R Durbin. Revising the human mutation rate: implications for understanding human evolution. Nat Rev Genet, 13:745, 2012.

72. S Schiffels and R Durbin. Inferring human popualtion size and separation history from multiple genome sequences. Nat Genet, 46:919–925, 2014.

73. DR Schrider. Background selection does not mimic the patterns of genetic diversity produced by selective sweeps. Genetics, 216:499–519, 2020.

74. DR Schrider and AD Kern. S/HIC: robust identification of soft and hard sweeps using machine learning. PLoS Genet, 12:1–31, 2016.

75. J Seger, WA Smith, JJ Prry, J Hunn, ZA Kaliszewska, L La Sala, L Pozzi, VJ Rowntree, and FR Adler. Gene genealogies strongly distorted by weakly interfering mutations in constant environments. Genetics, 184:529–545, 2010.

76. L Śegurel and C Bon. On the evolution of lactase persistence in humans. Ann Rev Genomics Hum Genet, 18:297–319, 2017.

77. D Setter, S Mousset, X Cheng, R Nielsen, M DeGiorgio, and J Hermisson. VolcanoFinder: genomic scans of adaptive introgression. PLoS Genet, 16:e1008867, 2020.

78. S Sheehan and YS Song. Deep learning for population genetic inference. PLoS Comput Biol, 12:1–28, 2016.

79. C Smukowski and M Noor. Recombination rate variation in closely related species. Heredity, 107:496–508, 2011.

80. Aaron J. Stern, Peter R. Wilton, and Rasmus Nielsen. An approximate full-likelihood method for inferring selection and allele frequency trajectories from dna sequence data. PLOS Genetics, 15(9):1–32, 09 2019. doi: 10.1371/journal.pgen.1008384. URL https://doi.org/10.1371/journal.pgen.1008384.

81. ZA Szpiech and RD Hernandez. selscan: an efficient multithreaded program to perform EHH-based scans for positive selection. Mol Biol Evol, 31:2824–2827, 2014.

82. Zachary A. Szpiech. selscan 2.0: scanning for sweeps in unphased data. bioRxiv, 2021. doi: 10.1101/2021.10.22.465497. URL https://www.biorxiv.org/content/early/2021/10/24/2021.10.22.465497.

83. Zachary A. Szpiech, Taylor E. Novak, Nick P. Bailey, and Laurie S. Stevison. Application of a novel haplotype-based scan for local adaptation to study high-altitude adaptation in rhesus macaques. Evolution Letters, 5(4):408–421, 2021. doi: https://doi.org/10.1002/evl3.232. URL https://onlinelibrary.wiley.com/doi/abs/10.1002/evl3.232.

84. N Takahata. Allelic genealogy and human evolution. Mol Biol Evol, 10:2–22, 1993.

85. D Taliun, DN Harris, MD Kessler, J Carlson, ZA Szpiech, R Torres, SA Gagliano Taliun, A Corvelo, SM Gogarten, HM Kang, AN Pitsillides, J LeFaive, S-b Lee, X Tian, BL Browning, S Das, A-K Emde, WE Clarke, DP Loesch, AC Shetty, TW Blackwell, AV Smith, Q Wong, X Liu, MP Conomos, DM Bobo, F Aguet, C Albert, A Alonso, KG Ardlie, DE Arking, S Aslibekyan, PL Auer, J Barnard, RG Barr, L Barwick, LC Becker, RL Beer, EJ Benjamin, LF Bialek, J Blangero, M Boehnke, DW Bowden, JA Brody, EG Buchard, BE Cade, JF Casella, B Chalazan, DI Chasman, IY-D Chen, MH Cho, SH Choi, MK Chung, CB Clish, A Correa, JE Curran, B Custer, D Darbar, M Daya, M de Andrade, DL DeMeo, SK Dutcher, PT Ellinor, LS Emery, C Eng, D Fatkin, T Fingerlin, L Forer, M Fornage, N Franceschini, C Fuchsberger, SM Fullerton, S Germer, MT Gladwin, DJ Gottlieb, X Guo, ME Hall, J He, NL Heard-Costa, SR Heckbert, MR Irvin, JM Johnsen, AD Johnson, R Kaplan, SLR Kardia, T Kelly, S Kelly, EE Kenny, DP Kiel, R Klemmer, BA Konkle, C Kooperberg, A Köttgen, LA Lange, J Lasky-Su, D Levy, X Lin, K-H Lin, C Liu, RJF Loos, L Garman, R Gerszten, SA Lubitz, KL Lunetta, ACY Mak, A Manichaikul, AK Manning, RA Mathias, DD McManus, ST McGarvey, JB Meigs, DA Meyers, JL Mikulla, MA Minear, BD Mitchell, S Mohanty, ME Montasser, C Montgomery, AC Morrison, JM Murabito, A Natale, P Natarajan, SC Nelson, KE North, JR O’Connell, ND Palmer, N Pankratz, GM Peloso, PA Peyser, J Pleiness, WS Post, BM Psaty, DC Rao, S Redline, AP Reiner, D Rode, JI Rotter, I Ruczinski, C Sarnowski, S Schoenherr, DA Schwartz, J-S Seo, S Seshadri, VA Sheehan, WH Sheu, BM Shoemaker, NL Smith, JA Smith, N Sotoodehnia, AM Stilp, W Tang, KD Taylor, M Telen, TA Thornton, RP Tracy, DJ Van Den Berg, RS Vasan, KA Viaud-Martinez, S Vrieze, DE Weeks, BS Weir, ST Weiss, L-C Weng, CJ Willer, Y Zhang, X Zhao, DK Arnett, AE Ashley-Koch, KC Barnes, E Boerwinkle, S Gabriel, R Gibbs, KM Rice, SS Rich, EK Silverman, P Qasba, W Gan, NHLBI Trans-Omics for Precision Medicine (TOPMed) Consortium, GJ Papanicolaou, DA Nickerson, SR Browning, MC Zody, S Zöllner, JG Wilson, LA Cupples, CC Laurie, CE Jaquish, RD Hernandez, TD O’Connor, and GR Abecasis. Sequencing of 53,831 diverse genomes from the NHLBI TOPMed Program. Nature, 590:290–299, 2021.

86. J Tennessen, AW Bigham, TD O’Connor, W Fu, EE Kenny, S Gravel, S McGee, R Do, X Liu, G Jun, HM Kang, D Jordan, SM Leal, S Gabriel, MJ Rieder, G Abecasis, D Altshuler, DA Nickerson, E Boerwinkle, S Sunyaev, CD Bustamante, MJ Bamshad, JM Akey, Broad GO, Seattle GO, and NHLBI Exome Sequencing Project. Evolution and functional impact of rare coding variation from deep sequencing of human exomes. Science, 337:64–69, 2012.

87. J Terhorst, JA Kamm, and YS Song. Robust and scalable inference of population history from hundreds of unphased whole-genomes. Nat Genet, 49:303–309, 2017.

88. The 1000 Genomes Project Consortium. A global reference for human genetic variation. Nature, 526: 68–74, 2015.

89. SA Tishkoff, FA Reed, A Ranciaro, BF Voight, CC Babbitt, JS Silverman, K Powell, HM Mortensen, JB Hirbo, M Osman, M Ibrahim, SA Omar, G Lema, TB Nyambo, J Ghori, S Numpstead, JK Pritchard, GA Wray, and P Deloukas. Convergent adaptation of human lactase persistence in Africa and Europe. Nat Genet, 39:31–40, 2007.

90. R Torres, ZA Szpiech, and RD Hernandez. Human demographic history has amplified the effects of background selection across the genome. PLoS genetics, 14(6):e1007387, 2018.

91. BF Voight, S Kudaravalli, X Wen, and JK Pritchard. A map of recent positive selection in the human genome. PLoS Biol, 4:e72, 2006.

92. HMT Vy and Y Kim. A composite-likelihood method for detecting incomplete selective sweep from population genomic data. Genetics, 200:633–649, 2015.

93. MA Wilson Sayres, KE Lohmueller, and R Nielsen. Natural selection reduced diversity on human Y chromosomes. PLoS Genet, 10:e1004064, 2014.

94. X Yuan, D J Miller, J Zhang, D Herrington, and Y Wang. An overview of population genetic data simulation. J Comput Biol, 19:42–54, 2012.

